# Procrastination partly reflects an evolutionary byproduct of non-planning impulsivity

**DOI:** 10.1101/2025.07.21.665826

**Authors:** Yuanyuan Hu, Jie Xiang, Yuening Jin, Qingchen Fan, Changshuo Wang, Yihan Wu, Dang Zheng, Bowen Hu, Tingyong Feng, Yuan Zhou, Zhiyi Chen

## Abstract

Procrastination has immediately visible repercussions on health and survival resilience, yet shows stably heritable and remains increasingly pervasive across human societies. Despite a paradox, this behavior is theoretically explained to represent a byproduct of evolutionary advantages underlying impulsivity, yet not deciphered well by scientific evidences. After adjusting psychometric endogeneity, we demonstrate the unique predictive roles of non-planning impulsivity (NPI) during late adolescence and early adulthood uniquely predicts procrastination in later adulthood in a twin cohort (*N* = 154). This association was further replicated in two independent cohorts (*N* = 327, *N* = 1,543). Using AE models, in conjunction of single-paper meta-analytic synthesis (*N* = 3,656 twin pairs), we observed significant shared genetic contributions underlying this NPI-procrastination association (*r*_g_ = 0.51, *95% CI*: 0.18 - 0.84). Beyond to the phenotypic heritability, employing a Genome-Wide-Association Study (GWAS), six NPI-procrastination overlapping SNPs are identified, functionally accounting for neural dysregulation. Thus, leveraging neurodevelopmental normative modelling (*N* = 37,407), online meta-analytic estimations (*k* = 198, loci = 5,855) and seed-based *d* mapping estimates (*N* = 893), cortical deviations in the left dorsolateral prefrontal cortex (DLPFC) - the brain region showing highest probabilistic overlap mapping NPI to procrastination, partly explains their shared genetic variants, but are substantially independent in genetic contribution. Mendelian Randomization analysis finally indicates causal roles of NPI and procrastination both, to DLPFC deviations. Our findings empirically clarified this theory that procrastination partly derives from NPI as an evolutionary byproduct indeed, but is still unique in neurogenetic entities.

**Brief summary:** Procrastination is a puzzling human behavior that compromises survival-relevant outcomes yet remains both widespread and heritable. Although theorized as a byproduct of impulsivity’s evolutionary advantages, empirical support for this account has been limited. Here, we provide converging evidence across psychometric, genetic, and neuroimaging modalities to show that non-planning impulsivity during late adolescence and early adulthood uniquely predicts later procrastination, and that the two traits share significant genetic overlap. We further identify specific genetic variants and morphological deviations in the DLPFC that link, but also partially dissociate, their biological pathways. These findings clarify the evolutionary and neurogenetic architecture of procrastination and underscore its partial derivation from impulsivity alongside distinct developmental origins.

## Introduction

Procrastination, the irrational delay of intended actions despite foreseeable negative consequences (1), is a remarkably pervasive problem in modern societies. The 75% of over 50,000 college students report chronic procrastination, while employees lose 1.5–3.0 hours daily in procrastination-related activities (2–4). Yet, procrastination’s repercussions extends far beyond its prevalence, imposing significant psychological distress and financial strain (5) whilst draining societal resources through lost productivity and economic inefficiencies, as well even deteriorating maladaptive lifelong coping patterns in response to environment demands (6–9). However, though procrastination casts plainly visible ramifications over individuals and society, it is stably heritable and is observed for the genomic loci (10, 11). Thus, it has yielded a long-lasting evolutionary paradox for why procrastination uncontrollably continues in this human populations, given no perceived societal and bioadaptive benefits.

To address this historical puzzle, many psychological theoretical evidence emerged, such as undervalued task execution motivation (12), goal-management failure (13, 14), emotional repair (15) and short-term behavior reinforcement (16), yet gaps remain in fully interpreting this paradox. One theory posits that procrastination, is not a cognitive or behavioral ingredient but an evolutionary byproduct stemming from impulsivity (17). This evolutionary conceptualization suggests that impulsivity is bioadaptive in premodern societies, strengthening survival and reproduction by promoting swift actions to immediate threats or rewarding opportunity (18). Although such evolutionary advantages have been overridden in modern environments that demand goal-directing behavior and long-term planning, these impulsive tendencies remain genetically ingrained, predisposing individuals to procrastinate by prioritizing immediate temptations. Supporting this argument, a bulk of studies reveal a positively phenotypic correlation between the two traits (*r*_*s*_= 0.23 - 0.70) (19–21). Furthermore, cross-sectional twin cohort studies show that procrastination and impulsivity share nearly identical genetic variation (*r*_g_ = 0.95 - 1.00), indicating their homologous evolutionary origins (13, 22). However, these observations deviate from this theoretical constructs, parsing impulsivity as a compound trait and thus misaligning to the most relevant structure — non-planning action and immediate rewarding priority. Loehlin and Martin (2014) demonstrate the prominently lower genetic association between both traits (*r*_g_ = 0.30) in a large-scale twin cohort (*N* > 3,000), appealing clarification into currently unexplained confounding mechanisms, particularly in the intricate neurogenetic architecture.

Liu and Feng (2017) demonstrate shared brain morphological covariance between impulsivity and procrastination in left dorsolateral prefrontal cortex (DLPFC), an evolutionary hub in the neocortex (20, 23, 24). This evolutionary advantage in brain organization is further supported by functional neuroimaging evidence: both traits are commonly overlapped onto DLPFC-centric central executive network (CEN), which is functionally implicated in integrating goal-direct action and long-term planning (25), reflecting an evolutionary hierarchy, echoing the notion that “we evolved to be procrastinators” (26–29). While the phenotyping of DLPFC as an evolutionary brain locus shared in both traits is gaining recognition, it remains more largely discussed conceptually rather than empirically elucidated. No neurogenetic evidence emerge to probe into this conceptualization underlying DLPFC-mediate evolutionary origins. Notably, shared brain morphological underpinnings are not exclusive within DLPFC. For instance, procrastinators and individuals with trait impulsivity exhibit common neuroanatomical atrophy in left middle frontal cortex and right anterior cingulate cortex (ACC), while showing increased regional volumes in the right parahippocampus cortex (PHC) (30–32). However, compared to the DLPFC, evidence for shared cortical structural changes in other brain regions remains inconsistent—these regions show neither clear opposite effects nor consistent lateralization patterns (33–35). While gaining increasing recognition for the shared neurogenetic origins underlying both traits, this evolutionary conceptualization has not yet empirically studied and the mechanistic understanding remains substantially unexplored.

As a proof-of-concept to this evolutionary theory, we first examined whether non-planning impulsivity (NPI), is predictive of procrastination **(Figure 1a)**. Despite conducting this analysis, one notable concern warrants caution in interpreting this conception: NPI may share the same psychological constructs as procrastination itself (20, 36). To address this issue, we further carried out a Gaussian graph-theoretical topological analysis to identify psychometric similarities in the specific items between the two traits, adjusting for these similarities as covariates of no interests to calibrate their unique phenotypic association **(Figure 1b)**. Upon confirming this conception, the multivariate AE (Additive gene - unique Environment) model was built up to clarify their shared genetic contributions, along with a single-paper meta-analytic estimate to synthesize this effect size **(Figure 1c)**. Beyond to behavioral genetic association, we conducted a Genome-Wide Association Study (GWAS) to capture their shared SNP signatures, enriching biologically mechanistic understandings **(Figure 1d)**. To clarify neurogenetic markers, we conducted two independent meta-analytic approaches: a seed-based d mapping (SDM) model and a separate analysis using the NeuroSynth platform. Both analyses systematically identified neural loci associated with the two traits, which were subsequently mapped from volumetric space to surface-based parcels **(Figure 1e)**. Once those shared neural loci were determined, we further employed neuroimaging normative modeling to investigate whether and how shared genetic associations may be underpinned by brain morphology, with a particular focus on the DLPFC, a region that has undergone rapid evolutionary changes **(Figure 1f)**. Finally, leveraging Mendelian Randomization (MR) analysis, we illustrated shared causal neurogenetic pathways underlying NPI and procrastination **(Figure 1g)**. Through this closed-loop behavior-gene-brain architecture, we aim to decode the neurogenetic underpinnings of procrastination, shedding light on its paradoxical persistence in modern societies.

**Figure 1.**
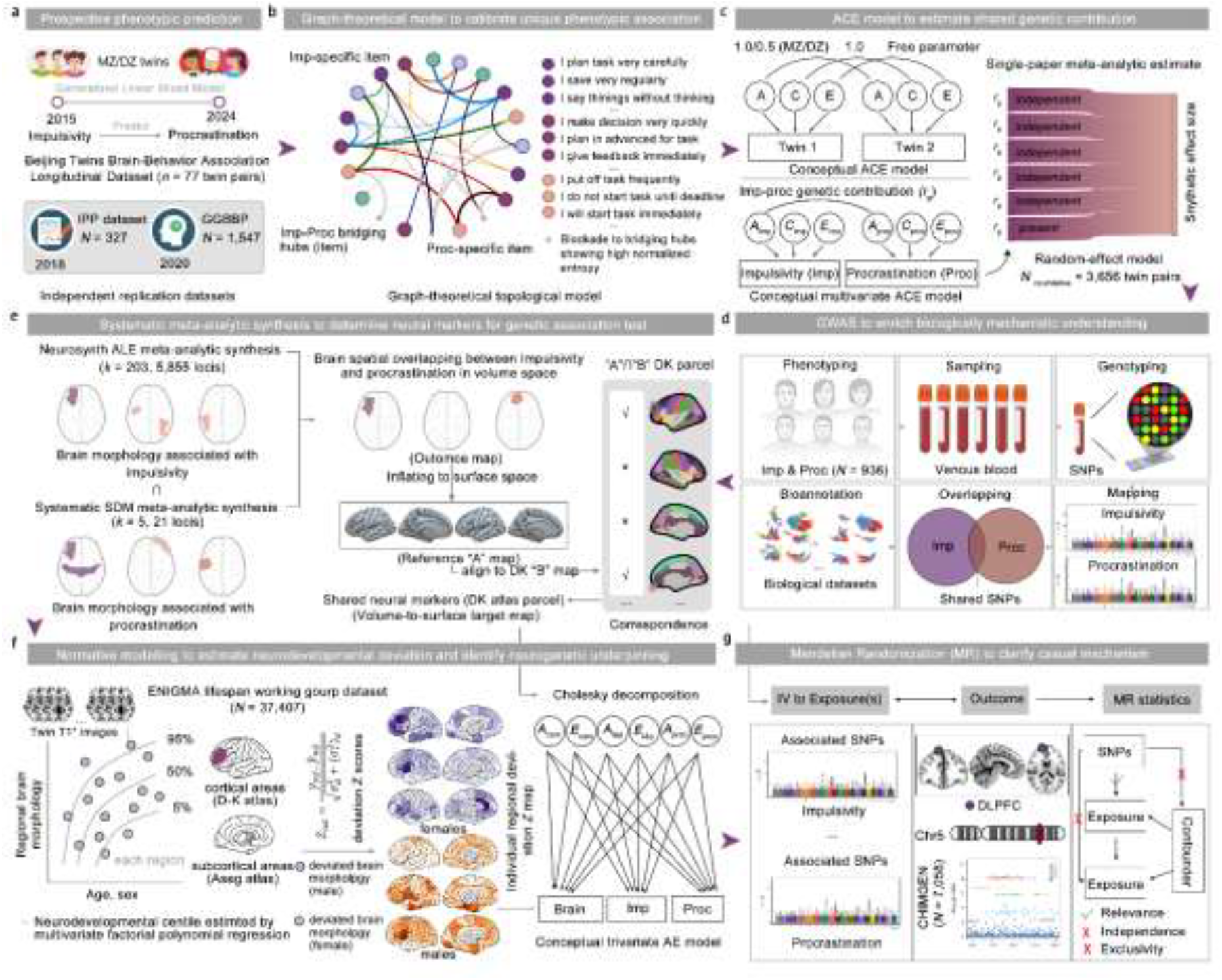
The integrative framework testing the evolutionary neurogenetic basis of procrastination. **a** Using an 9-year longitudinal twin cohort and two independent samples, we first tested whether adolescent NPI prospectively predicts procrastination in early adulthood. **b** To control the psychometric overlap between impulsivity and procrastination, a graph-theoretical topological model was applied to item-level data to identify bridging constructs and adjust for redundant variance. **c** Univariate and bivariate ACE twin models were constructed to estimate shared genetic contributions between NPI and procrastination, with single-paper meta-analytic synthesis to aggregate effect sizes. **d** A GWAS identified overlapping SNPs and enriched functional annotations in an independent sample, offering molecular-level insights into shared biological mechanisms. **e** Meta-analytic neuroimaging synthesis combined NeuroSynth and SDM approaches to identify convergent brain regions associated with both traits. **f** Normative neurodevelopmental deviation was quantified using structural MRI from a population reference dataset and applied to the twin cohort to test whether shared morphological loci explain phenotypic and genetic associations. **g** MR was performed to test the causal relationships between genetically predicted impulsivity/procrastination and structural deviations in the DLPFC, a region implicated in future-oriented planning and uniquely human cognition.

## Results

### Samples and designs

This study first utilized a 8-year Longitudinal Twin Cohort that included 154 participants (77 pairs: monozygotic twins, MZ = 40 pairs; dizygotic twins, DZ = 37 pairs) for phenotypic and genetic association analysis. Considering reproducibility of these findings, two independent samples were included (Cross-sectional Adolescent Cohort: *N* = 327; Cross-sectional Large-scale Adult Cohort: *N* = 1,543) **(*Supplementary Material* Tab. S1)**. Given notable heterogeneity reporting genetic associations between impulsivity and procrastination (*r*_g_ = 0.29 - 1.00) (13, 37), a single-paper meta-analytic sample synthesizing 3,656 twin pairs from existing empirical studies was curated to estimate this pooled effect size (*k* = 4, ***see Methods***) **(*Supplementary Material* S3 Tab S2)**. Upon demonstrating these shared genetic associations, we enrolled an additional sample for GWAS, encompassed by 936 participants with Asian ancestry **(*Supplementary Material* Tab. S1)**. To localize potential neurogenetic marker(s) shared between the both traits, a systematic neuroimaging meta-analytic sample and an online meta-analytic decoding were generated by 508 and 198 independent primary studies, respectively **(*see Methods, Supplementary Material* S5)**. Moving forward to enrich their neurogenetic understanding, the ENIGMA lifespan sample consisted of 37,407 participants with T1* neuroimaging data, aged from 3 to 90 years, was integrated to calculate neurodevelopmental deviations in Twin Adolescent Cohort in a region-by-region way **(*Supplementary Material* Tab. S1)**. Upon revealing shared neurogenetic substrates, we finally included 7,058 Chinese Han participants from CHIMGEN-GWAS dataset for capturing causally neurogenetic pathways in the MR analysis **(*Supplementary Material* Tab S1)**.

### NPI significantly predicts procrastination at the phenotypic level

Using this 8-year longitudinal twin cohort, we found that NPI during late adolescence and early adulthood uniquely predicts procrastination in later adulthood (*β* = 0.38, *p* = 0.016, *R*^2^ = 0.44), significantly surpassing other subconstructs underlying impulsivity (i.e., Attentional Impulsivity and Motor Impulsivity) **(Figure 2a and *Supplementary Material* S1 Tab S1)**. This finding was replicated in two independent samples, showing high reproducibility **(*Supplementary Material* S1 Tab. S1)**. Using single-paper meta-analytic model to synthesize these effect sizes, it demonstrated a statistically robust association between NPI and procrastination (*d* = 0.61, 95% CI: [0.44, 0.77], α = 0.99, *p* < .001) **(*Supplementary Material* S1 Tab S3)**. At a phenotypic level, this evolutionary theory was empirically supported by showing predictive powers of NPI to procrastination.

**Figure 2.**
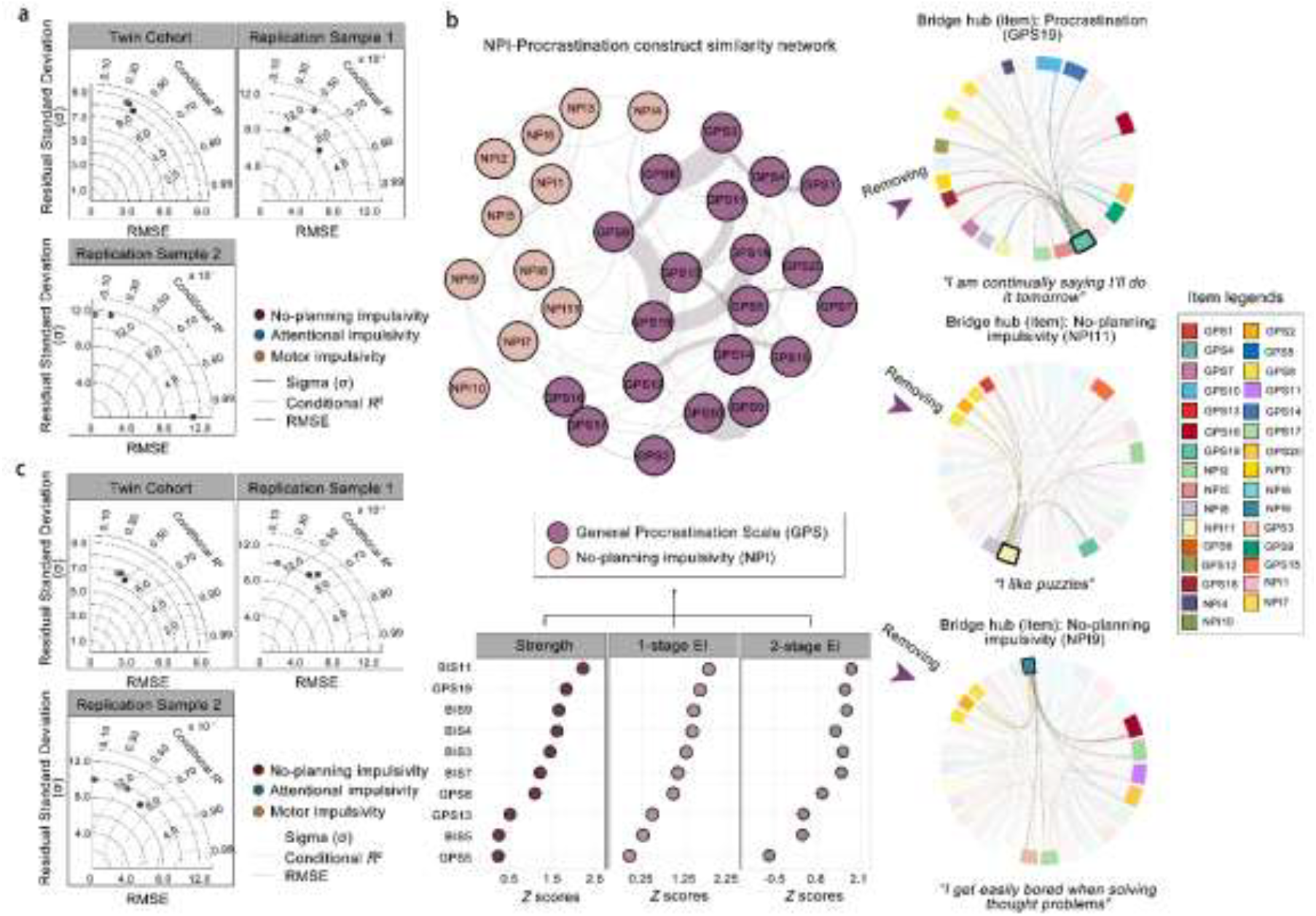
NPI prospectively predicts procrastination and remains robust after accounting for psychometric overlap. **a** In a longitudinal adolescent twin cohort, NPI significantly predicted procrastination, outperforming other impulsivity subcomponents. This effect was replicated in two independent samples, and standardized residuals confirmed the model’s robustness across Sigma, RMSE and conditional R^2^. **b** To evaluate potential psychometric overlap between NPI and procrastination, a graph-theoretical item similarity network was constructed. Three bridge items— one from the GPS and two from the BIS-11—were identified as shared hubs and removed to control for between-scale psychometric similarity. **c** After removing overlapping items, the predictive association between NPI and procrastination remained significant and replicable across samples.

Nevertheless, one notable concern to question this conclusion is psychometric similarity between the two traits, that is to observe psychological construct’s resemblance for what both scales measured (20, 36). To clarify, we established a graph-theoretical network model by incorporating all items from both scales meanwhile for identifying which items were measured similarly in item-centered statistical structures **(*see Methods*)**. In this model, items showed statistically higher bridge centrality once they were central to connect between-scale items than other peripheral ones, thus implying psychometric similarity. We identified three “bridge” items (hubs), with two items *“I get easily bored when solving thought problems”* and *“I like puzzles”* from the BIS Non-planning subscale and with one *“I am continually saying I’ll do it tomorrow”* from the GPS scale **(Figure 2b, *Supplementary Material* S2)**.

After removing these items to control between-scale psychometric similarity, we calibrated unique phenotypic associations between NPI and procrastination, showing the similar findings (*β*_original_ **=** 0.38, *β*_calibrated_ **=** 0.34). This calibrated result was fully replicated as well **(Figure 2c, *Supplementary Material* S1-S3**), Leveraging the single-paper meta-analytic model, this synthetic effect size to their unique phenotypic associations exhibited slightly decreased but robust (*d* = 0.50, 95% CI: [0.38 - 0.63], α = 0.99, *p* < .001). Compared to overlapped factor structures between original scales, both measurements were observed statistically separate after removing these three items **(*Supplementary Material* S2 Tab S3)**. Taken together, the impulsivity, particularly in the NP portrait, prospectively and robustly predicted procrastination, theoretically indicating that procrastination may be its resultant byproduct in the phenotypic layer, at least.

### NPI and procrastination share significant genetic contributions

Upon identifying the robust phenotypic association between NPI and procrastination, we further investigated their genetic underpinning using ACE structural equation modeling method **(*see Methods*)**. Here, the nested AE model was established as benchmark for estimating quantitative genetic statistics because it showed best fits **(*Supplementary Material* S3 Tab S1)**. We found that NPI exhibited a heritability of 37.6% (*h*^2^, 95% CI: [0.08 - 0.60]), while procrastination demonstrated a higher heritability of 63.01% (95% CI: [0.41 - 0.77]) **(Figure 3a)**. More importantly, beyond to show separately heritable, both traits were identified to share genetic contributions (*r*_*g*_ = 0.42, 95% CI: [0.10, 0.70]; **Figure 3a**). Similarly, using a single-paper meta-analytic model, this genetic correlation was calibrated to be 0.66 by synthesizing four effect sizes from existing studies (*p* < .001**; Figure 3c-d and *Supplementary Material* S3 Tab S2**). Therefore, these empirical evidences demonstrating the moderate genetic correlation between NPI and procrastination, partly supports this theoretical conceptualization that both traits shared a significant but nonidentical genetic contribution.

**Figure 3.**
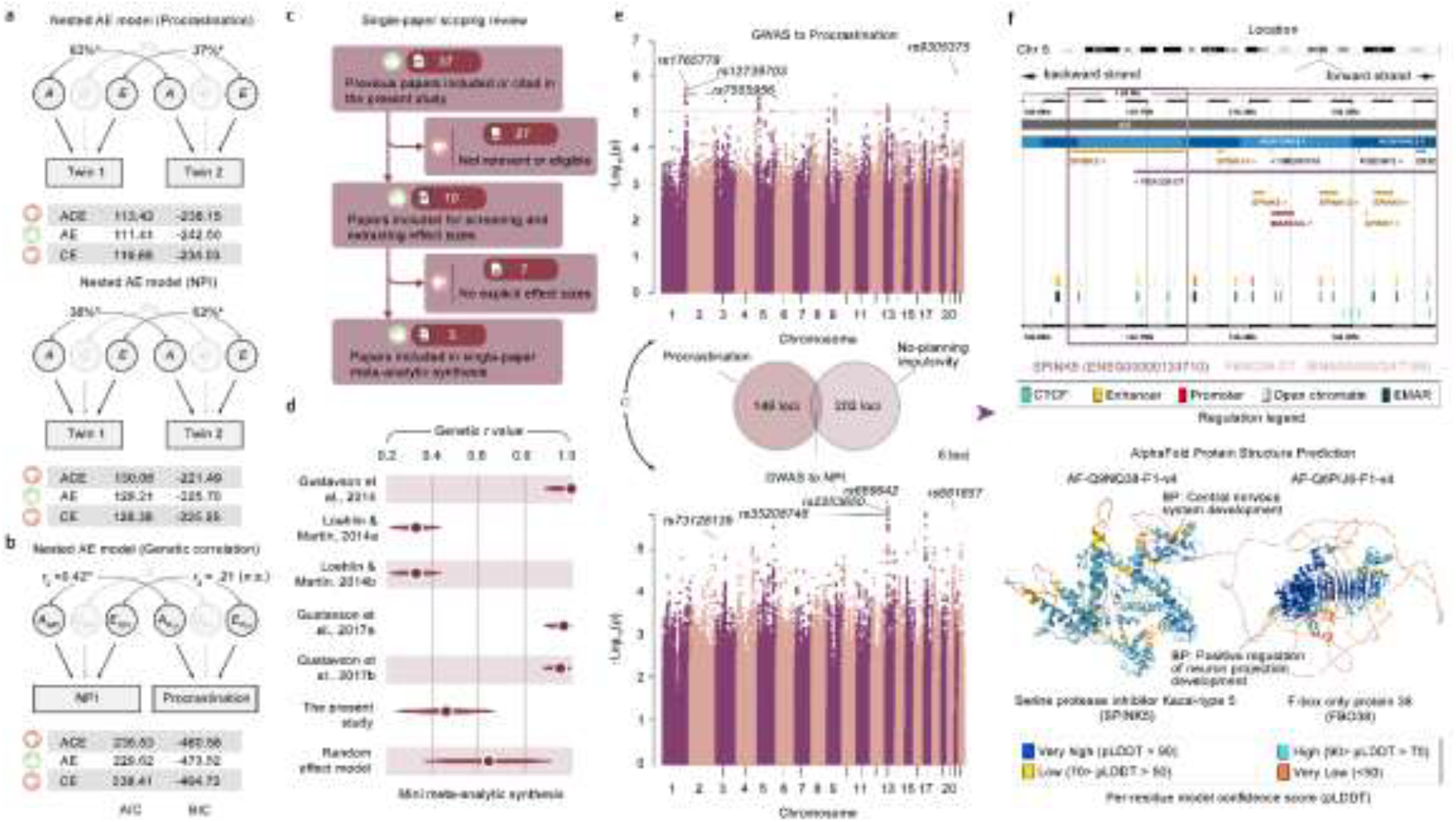
NPI and procrastination share genetic contributions and overlapping neurodevelopmental loci. **a** Univariate ACE models revealed that both NPI and procrastination are heritable traits. **b** Bivariate ACE model showed significant genetic overlap between the two traits, suggesting partially shared heritable influences. **c** Genetic correlations between impulsivity and procrastination from three independent studies were synthesized using a single-paper meta-analytic model. **d** Forest plot summarizing the meta-analytic results, confirming the robustness of the genetic association between NPI and procrastination. **e** GWAS identified multiple candidate SNPs for each trait, with six loci overlapping across NPI and procrastination. **f** Shared loci were functionally annotated to genes involved in neurodevelopmental processes, including *SPINK5* and *FBXO38-DT*, with protein structures predicted using *AlphaFold*.

### NPI and procrastination overlap in the specific genetic loci associated with neurodevelopment

Despite manifesting shared genetic contributions between NPI and procrastination, the specific genomic entities underlying their association remain unexplored. Based on GWAS, we identified 149 loci candidates for procrastination (*p* < 5 × 10^−5^; ***Supplementary Material* S7 Tab S2**), such as in *rs9305375, rs1765779, rs12739703* and *rs7555956*, while discovered 202 candidates for NPI, particularly in *rs681657, rs669642, rs2253650 and rs35208748* **(Figure 3e and *Supplementary Material* S7 Tab S3)**. Therefore, six loci were found to be overlapped between the two traits, including *rs9325070, rs9791022, rs111890148, rs72660259, rs17107747* and *rs6887442* **(Figure 3e)**. Leveraging Ensembl Variant Effect Predictor (VEP, https://asia.ensembl.org/Tools/VEP) for biological ontology annotation ***(see Methods)***, these loci were found to be at closest in *SPINK5* (Serine protease inhibitor Kazal-type 5) and *FBXO38-DT* (F-box only protein 38), with partial functions to the specific protein coding, which all located in Chromosome 5 **(Figure 3f)**. Thus, their protein structures were estimated via AlphaFold (https://alphafold.ebi.ac.uk), which were annotated in associating with biological processes underlying “Central nervous system developments” and “Positive regulation of neuron projection developments” **(Figure 3f, *Supplementary Material* S7 Tab S4-S5)**. In sum, these findings extend beyond to demonstrate a general genetic correlation by capturing specific genomic loci candidates that overlapped between NPI and procrastination, thereby elucidating their potential biological functions in neurodevelopment.

### DLPFC and mPFC contribute to neurodevelopmental loci shared between the both traits

Given the involvement in neurodevelopment, we further examined specific neural localizers that may be shared between impulsivity and procrastination. Leveraging online NeuroSynth meta-analytic estimations (*k* = 198, *loci* = 5,855), for impulsivity, we clarified significant likelihood activation in several key parcels of frontal cortex, including left DLPFC (MNI coordinate: −40, 46, 4), right caudate (MNI coordinate: 10, 14, −4), as well as left ventromedial prefrontal cortex (vmPFC, MNI coordinate: −6, 13, −4) **(Figure 4a)**. Full results including peak coordinates, activation strength, cluster size, and anatomical labels are provided in ***Supplementary Material* S5 Tab. S3**. Furthermore, as the meta-analytic *d* mapping model identified from a systematic neuroimaging synthesis, neural signatures in procrastination included clusters of left DLPFC (MNI Coordinate: −34, 42, 30), right PHC (MNI coordinate: 20, −10, −30), and left medial prefrontal cortex (mPFC, MNI coordinate: 0, 46, −10) **(Figure 4a and *Supplementary Material* S5 Tab S2)**. Therefore, we identified spatial overlapping between the both traits as reference map representing their shared neural loci, including left DLPFC (MNI coordinate: −40, 42, 20) and mPFC (MNI coordinate: 0, 46, −10) **(Figure 4a and *Supplementary Material* S5 Tab S4)**. Finally, mapping these shared loci from volumetric space to cortical surfaces ***(see Methods)***, they are identified to overlap in rostral middle frontal cortex (i.e., DLPFC) and medial orbitofrontal cortex (i.e., mPFC), as the Desikan-Killiany cortical atlas parceled. Taken together, it resonates this notion that the DLPFC and mPFC may be potential neural markers representing shared neurodevelopmental vulnerability between impulsivity and procrastination.

**Figure 4.**
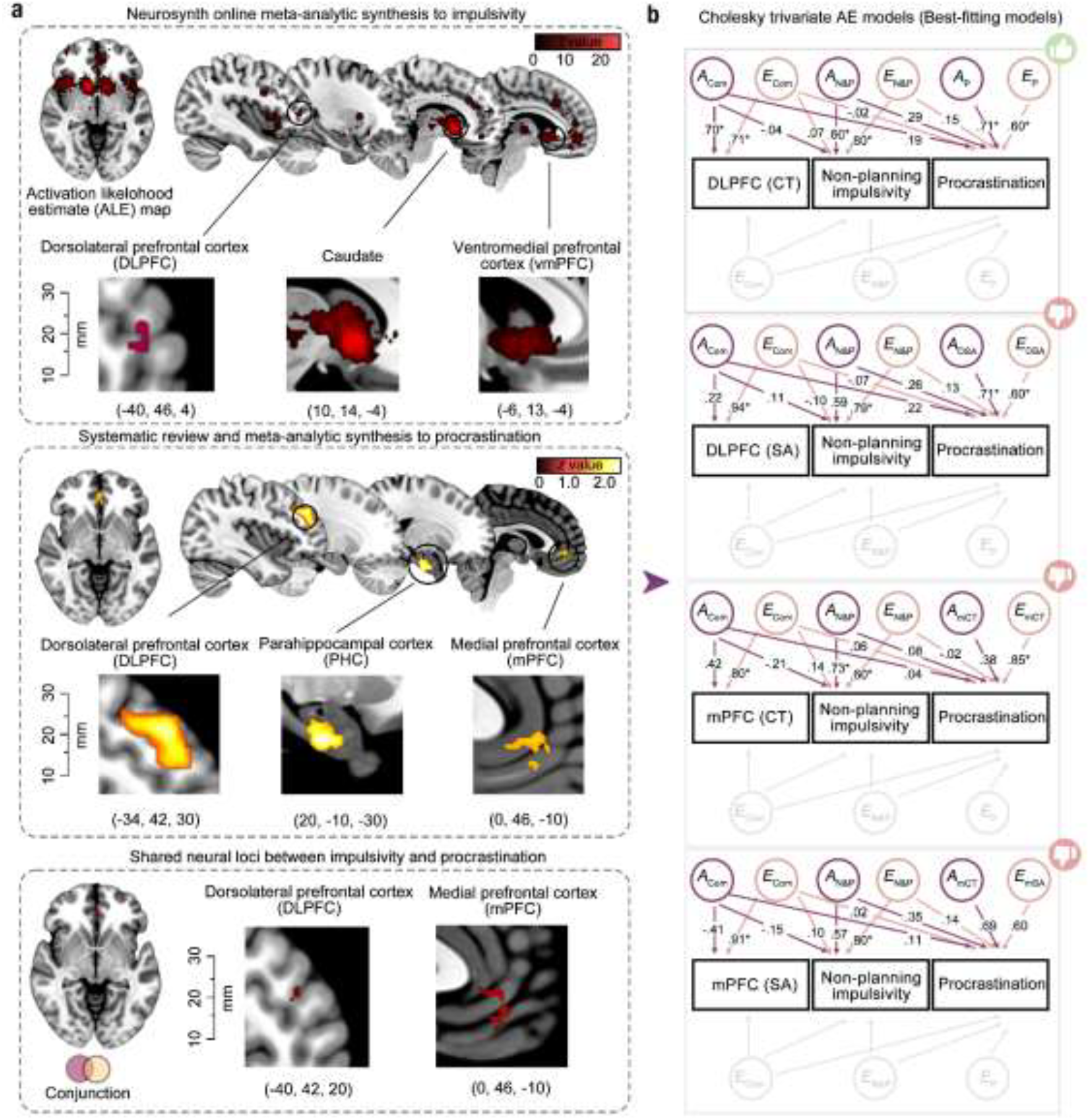
DLPFC and mPFC reflect shared neural loci underlying impulsivity and procrastination. **a** Meta-analytic neuroimaging synthesis identified overlapping brain regions associated with impulsivity and procrastination. Shared volumetric loci were mapped onto surface-based cortical parcels, revealing convergence in the left DLPFC and mPFC. **B** Trivariate ACE model showed that deviations in DLPFC cortical thickness partly explained the genetic variance shared between impulsivity and procrastination. However, procrastination also exhibited unique genetic contributions, and surface area deviations in DLPFC or mPFC did not significantly account for their shared genetic variance.

### Neurodevelopmental deviations in left DLPFC partly explain their shared genetic association, while procrastination poses uniquely genetic variants

Based on these shared neural loci identified above (i.e., DLPFC and mPFC), we further modeled their individual morphological deviations to investigate neurogenetic overlap between the both phenotypes, particularly in DLPFC. Using a normative modelling approach in the Longitudinal Twin Cohort, we identified neurodevelopmental deviations in both cortical surface areas (SA) and cortical thickness (CT), especially in the frontal areas **(see *Supplementary Material* S6)**. Employing a Cholesky trivariate AE model **(see *Methods* and *Supplementary Material* S4 Tab S1-S3)**, we found that: the latent genetic factor (*A*_com_) accounted for a substantial proportion of the variations in cortical thickness deviations in left DLPFC (a_1_ = 0.70, 95% CI: [0.48 - 0.90]). When the genetic contribution from deviations in the left DLPFC was constrained, the genetic factor shared between NPI and procrastination (*A*_N&P_) explained a significant portion of variance in NPI (a_2_ = [0.60, 95% CI: [0.21- 0.85]), but not in procrastination. Notably, procrastination exhibited a substantial unique genetic component (a_3_ = 0.71, 95% CI: [0.40-.90]) **(Figure 4b and *Supplementary Material* S4 Tab S7)**. However, neither SA deviations in DLPFC nor in mPFC significantly explained their shared genetic variants **(Figure 4b and *Supplementary Material* S4 Tab S7)**. These findings collectively indicate that while DLPFC deviations partially account for shared genetic variants between NPI and procrastination, the procrastination still pose uniquely genetic contribution.

### NPI and procrastination causally links with left DLPFC morphology

While these Cholesky decomposition models conceptually identify directional associations between brain morphology and phenotypes, their causally neurogenetic roles should be clarified. Employing bidirectional two-sample MR analyses by incorporating brain image-derived phenotype GWAS data from CHIMGEN (38) **(*see Methods*)**, we examined whether DLPFC causally shared as the neurogenetic marker to NPI and procrastination. The inverse-variance weighted (IVW) analysis revealed significant causal roles of both NPI and procrastination to left DLPFC morphology (*β*_*NPI*_ = −0.05, *p* = .044; *β*_*Procrastination*_ = −0.01, *p* = .052) **(Figure 5 and S7 Tab S1)**. No statistically significant heterogeneity and horizontal pleiotropy were identified in both IVW models (for NPI as exposures, Cochran’s *Q* tests, *Q* = 0.83, *p* = .661, MR-Egger regression test, *intercept* = 0.02, *p* = .530; for procrastination as exposures, *Q* = 0.72, *p* = .867, *intercept* = 0.03, *p* = 0.698) **(*Supplementary Material* S7.3)**. In addition, sensitivity analyses and reverse causation tests demonstrated computational robustness of these findings **(*Supplementary Material* S7.3)**. Thus, while these findings fail to directly support this hypothesis, it may delineate this notion that NPI and procrastination do not fully share in neurogenetic substrates but render the similar neurobiological outcomes in the DLPFC.

**Figure 5.**
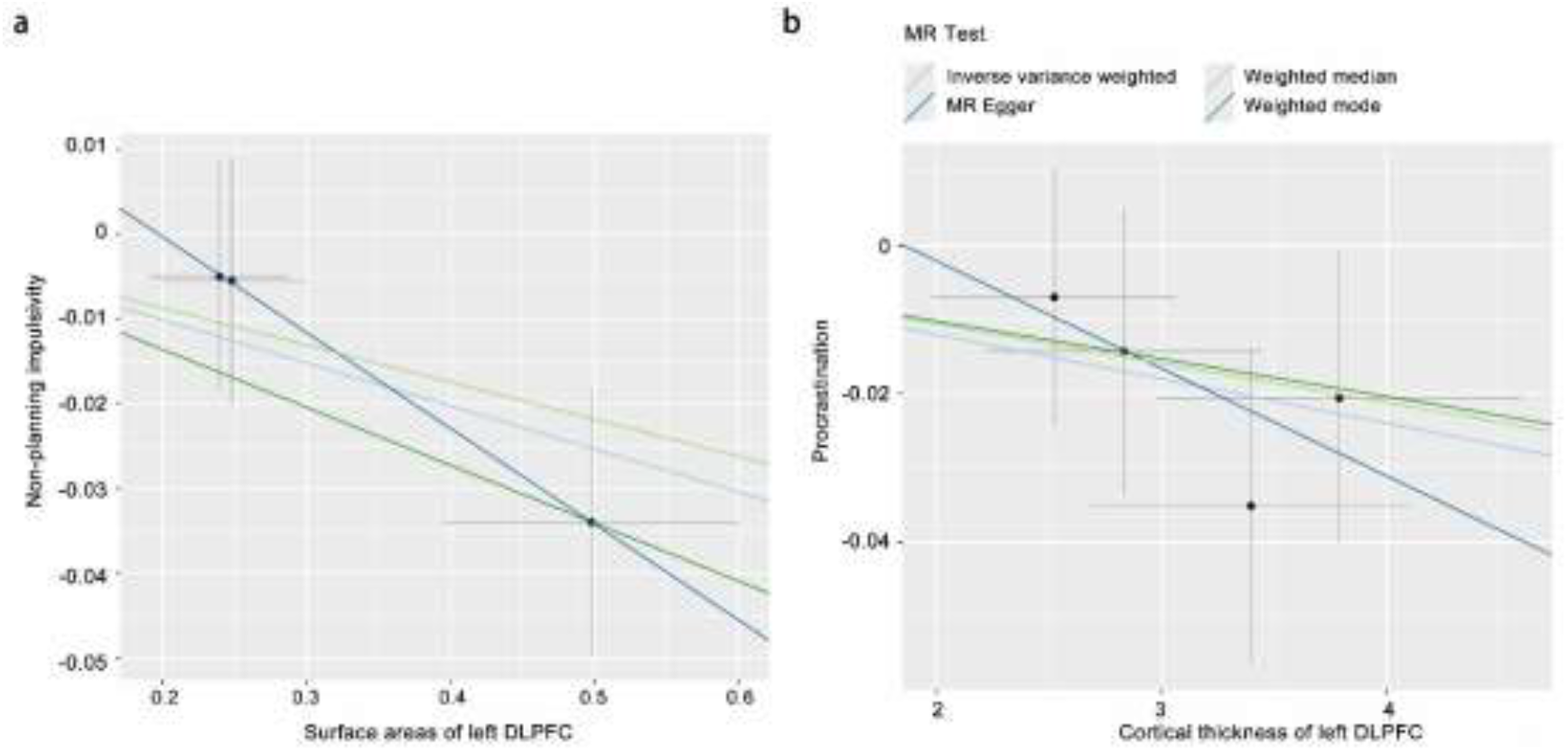
NPI and procrastination exhibit causal influences on DLPFC morphology. Bidirectional MR analyses assessed whether NPI and procrastination causally impact structural features of the DLPFC. The plot visualizes instrument-level associations, while causal estimates were derived using the IVW method as the primary analytical approach. Each dot represents a single SNP, with *x-* and *y-*values indicating its *β* on the exposure and outcome, respectively. Error bars denote standard errors. Results suggest both NPI and procrastination showed significant negative causal effects on DLPFC structure.

## Discussion

In this study, to address this long-lasting theoretical debate - whether human procrastination derives from impulsivity as an evolutionary byproduct, we employed a longitudinal twin design for examining neurogenetic underpinning between impulsivity and procrastination. Our findings revealed that NPI in late adolescence and early adulthood prospectively predicted their procrastination trait in later adulthood. This impulsivity-procrastination prediction indicated a significant phenotypic association, and has been replicated in independent samples. Furthermore, using AE model, both traits exhibited moderate-to-high heritability, and significantly shared genetic influences in accounting their phenotypic association, partly supporting this notion that procrastination may share as a byproduct. To delineate genetic mechanism underlying this association, we have performed GWAS, and demonstrated statistically significant SNPs overlapped between the both traits, which were involved into specific protein coding on neurodevelopment. Therefore, through systematic meta-analytic synthesis, DLPFC and mPFC were identified as shared impulsivity-procrastination loci. By calculating neuroanatomical developmental deviations from normative modelling for them, we found that neurodevelopmental deviations in DLPFC partly explained NPI-procrastination shared genetic variants, while procrastination posed substantially unique genetic contribution. Though this implied DLPFC as a neurogenetic mechanism underlying NPI-procrastination mapping, MR analyses demonstrated that both NPI and procrastination causally predicted DLPFC deviations, rather than the other way round. Taken together, this study systematically clarified neurogenetic underpinning of NPI-procrastination mapping, thus supporting this theoretical notion that procrastination may partly reflects a byproduct from impulsivity.

This study reveals a robust phenotypic and genetic association between procrastination and impulsivity, particularly in non-planning component, thereby providing partial empirical support for the theoretical proposition that procrastination may represent an evolutionary byproduct of impulsivity. The hypothesis that impulsivity—characterized by diminished foresight and reduced goal-directed deliberation—may have conferred adaptive advantages through rapid decision-making in resource competition has long been posited in evolutionary psychology (39–41). According to this framework, the cognitive architecture underlying short-sighted goal pursuit, which prioritizes immediate over long-term outcomes, is thought to also underpin procrastination (42–44). However, a bulk of evidence supporting this theoretical notion remain two fundamental gaps, with one for confounding psychometric similarities between the two traits and with another for obscuring conceptual structure. Despite ample empirical evidence corroborating shared phenotypic and genetic underpinning between impulsivity and procrastination (13, 22, 29, 45, 46), this statistical association was critiqued for potential overestimation from similar measurements, particularly where the impulsivity was identified as a complicated construct covering multifaceted traits and behaviors, such as procrastination (47, 48). Therefore, by adjusting for psychometric bridging items from statistical modeling in the present study, we provide clear empirical evidence to support the “pure” association between the two traits. While procrastination has been theorized to co-evolve with impulsivity through rapid, short-sighted decisions favoring immediate rewards, existing empirical studies have predominantly assessed impulsivity as a multifaceted construct (e.g., using measures such as BIS-11 or UPPS) (20, 22, 37, 49), rather than focusing on this specific theoretically-derived component implicated in procrastination. Rather than merely replicating prior findings, this study narrows impulsivity measurement to its non-planning dimension, bridging the gap between theoretical conceptualization and statistical estimation. Our results provide straightforward evidence suggesting that procrastination may emerge as a behavioral byproduct of impulsive traits.

Beyond clarifying the shared behavioral genetic underpinning, we further identify six overlapping genomic loci between NPI and procrastination, shedding light on potential biological mechanisms that contribute to their common genetic architecture. Here, the identified SNPs map to two nearest genes—*SPINK5* and *FBXO38-DT*. Although neither gene shows direct functional enrichment for NPI or procrastination, both have been implicated in protein encoding or regulatory processes for neurodevelopments (50–52). Specifically, as a serine protease inhibitor (53), *SPINK5* expression determines maintenance of neuronal homeostasis, regulation of synaptic plasticity, and modulation of neuroinflammation, which are all involved in neurodevelopmental processes (54–56). It has been echoed that the *FBXO38-DT*, a regulatory long non-coding RNA, has been substantially identified in modulating central nervous system development and positive neuronal projection, posing close associations with neurological disorders (52, 57). Thus, as articulated above, these findings suggest that the shared loci identified may influence neurodevelopmental processes that underlie both NPI and procrastination. Supporting this argument, carriers of the *COMT 158Val* allele exhibit higher levels of procrastination compared to 158Met carriers—a polymorphism robustly linked to dopaminergic modulation in neurodevelopmental conditions (10, 58, 59). Taken together, these findings enriches our understanding of genetic mechanisms shared between the NPI and procrastination, highlighting biologically regulatory roles of neurodevelopmental processes in their association.

A key finding is the identification of the left DLPFC as a shared neurogenetic locus between impulsivity and procrastination—a region critically involved in evolutionary neurodevelopment. As a major cortical expansion from primate to human, the DLPFC reflects an evolutionary functional specialization to undergird high-order cognition, particularly where temporal evolution in DLPFC has been captured in shaping self-regulation and goal-direct decisions (23, 60, 61). Thus, procrastination and impulsivity may represent behavioral outcomes of DLFPC evolutionary neurodevelopment meantime, rather to separate or sequential traits (62, 63). Supporting that, genetic variations linked to the DLPFC have been implicated in the regulation of decisions involving long-term rewards and goals, these cognitive components that substantially underpin both impulsivity and procrastination (64, 65). Notably, in addition to the shared neurogenetic basis in the DLPFC, procrastination exhibits substantial unique genetic variation, pointing to partially distinct evolutionary origins. This point has been previously examined through evidence of limited genetic correlation between the two traits (*r* = 0.23) (37), a finding corroborated in the present study, which reveals a moderate genetic correlation between non-planning impulsivity and procrastination. Therefore, taking together, these findings indicate that although the DLPFC serves as a neurogenetic locus for procrastination, its evolutionary mechanism cannot be fully explained as a mere “byproduct” of adaptive self-regulation strategies linked to impulsivity. Instead, it may be more appropriately framed within a reward-encoding system rooted in limbic regions (66, 67). On balance, the present study not only identifies a shared neurogenetic locus underlying the both traits, supporting the theoretical notion that procrastination may represent an evolutionary “byproduct” of non-planning impulsivity, but also reveals substantial unique neurogenetic contributions specific to procrastination.

An unexpected finding worthy to discuss is that both procrastination and non-planning impulsivity causally contribute to DLPFC morphology, rather than the reverse. It has long been argued that neurodevelopmental anomalies in left DLPFC, particularly in the neuroanatomical deviations, result in procrastination tendency due to DLPFC-specific flaws in top-down regulation and cognitive control (29, 30, 34, 68). Supporting this point, Xu and colleagues (2023) demonstrated that anodal neuromodulation to the DLPFC causally attenuates procrastination willingness by enhancing the perceived value of task outcomes, thereby providing strong evidence for a directional influence from DLPFC neural dynamics to behavioral phenotype (69). Given the rapid evolution in neocortex, particularly in DLPFC, the neurobiological functions of this region has emerged as key determinants of psychological, cognitive and behavioral outcomes in modern society (70, 71). Therefore, procrastination has theoretically modeled as a outcome from DLPFC-specific dysfunction (72, 73). Despite this, our finding recasts a reconsideration of the brain-behavior relationship in procrastination, particularly in light of evidence that DLPFC morphology is causally influenced by both traits. In conjunction with prior observations showing that procrastination carries substantial unique genetic variation, these results suggest that the trait may contribute to epigenetic modifications in neuroanatomy (74, 75). In other words, a new conceptual framework for understanding the evolutionary origins of procrastination emerges: rather than being shaped by passive genetic regulation alone, procrastination and impulsivity may actively influence brain plasticity in response to the evolving demands of modern society (76–78). In summary, while we demonstrate a shared neurogenetic basis between procrastination and non-planning impulsivity, our findings also suggest that the two traits may reciprocally regulate the brain locus through behavior–brain epigenetic mechanisms.

While this study offers valuable insights into the neurogenetic mechanisms underlying non-planning impulsivity and procrastination, several limitations should be warranted. Due to the absence of baseline procrastination measurements, our findings are limited to providing evidence for prospective prediction rather than causal inference regarding the association between the two traits. Despite compromising sample size from this longitudinal design in the present study, enlarging twin neuroimaging sample size could undoubtedly increase robustness and reliability for our findings in the future. Furthermore, based on a hypothesis-driven framework, we narrow neurogenetic analyses on discrete brain loci identified through systematic meta-analytic synthesis of neuroimaging overlap between procrastination and non-planning impulsivity. Thus, expanding investigations to encompass broader brain regions and even large-scale networks beyond those limited loci may provide a more comprehensive understanding of the underlying neurogenetic architecture. Moreover, compared to the standard GWAS significance threshold, we adopted a more lenient threshold (*p* < 5 × 10^−5^), which nevertheless meets an acceptable level of statistical credibility (*q* < .05). As a result, genomic loci that identified in the present study reach “suggestive statistical significance” rather “genome-wide statistical significance”. Finally, though observing unique genetic and environmental contributions, the AE model that we used in the present study fails to quantitatively estimate how they interacted in regulating behavioral phenotype (i.e., G × E effect). To comprehend roles of G × E interplay in shared neurogenetic underpinning between the two traits, twin interaction model and GWAS-based G × E model could be added in the future.

In conclusion, this study provides empirical support for ongoing theoretical debates regarding the origins of procrastination—demonstrating that it may indeed partly emerge as an evolutionary byproduct of non-planning impulsivity, with neurodevelopmental deviations in the DLPFC accounting for their shared neurogenetic architecture. Moreover, procrastination exhibits substantial unique genetic variants and demonstrates causal influences on shaping DLPFC morphology, suggesting the distinct behavior-brain epigenetic mechanism that contribute to its neurogenetic origins.

## Materials and Methods

### Human participants and ethic statements

This study includes multiple human samples, encompassed by phenotyping data, twin neuroimaging data and genomic sequencing data. The 8-year longitudinal twin neuroimaging cohort derives from Beijing Twin Study (BeTwiSt), consisted of 77 twin pairs acquiring T1* neuroimaging data at 2015 (monozygotic twins, MZ = 40 pairs; dizygotic twins, DZ = 37 pairs) and measuring procrastination at 2024 in the follow-up. Two independent cohorts acquiring phenotyping data are included as replication samples (*N* = 327, *N* = 1,543). A total of 936 samples from participants of Asian ancestry were initially included for genomic analysis phenotyping for genome of impulsivity and procrastination. Moreover, a large-scale neuroimaging dataset established by the ENIGMA has been included to model normative neurodevelopmental trajectory (*N* = 37,407, with 19,826 females, aged from 3 to 90 years ago). For bridging causal loops of macroscale neuroimaging loci to molecular genomic variants, in the MR analysis, the normative CHIMGEN-GWAS dataset to mark significant SNPs underlying each regional cortical morphology, consisted of 7,058 Chinese Han participants, and GWAS dataset phenotyping for genome of impulsivity and procrastination has been included. For additional information, see ***Supplementary Material* Tab. S1**.

Written informed consents have been acquired preceding to conduct this study. The Institutional Review Board (IRB) of Chinese Academy of Sciences (Institute of Psychology, #H15024 and #H20037), the Southwest University (Faculty of Psychology, #H24121), the Research Ethics Committee of Southwest University and the Medical Ethics Committee of Chongqing Ninth People’s Hospital (CNPH2019016) reviewed and formally approved these studies involving into human participant data.

### Measurement of Impulsivity and Procrastination

#### Barratt Impulsiveness Scale (BIS-11)

We use the Barratt Impulsiveness Scale (BIS-11) (79) to assess impulsivity in the Longitudinal Twin Adolescent Cohort and Cross-sectional Adolescent Cohort. The BIS-11 is a 30-item self-report instrument that quantifies trait impulsivity across three distinct domains: (1) attentional impulsivity (e.g. I don’t “pay attention”?); (2) motor impulsivity (e.g. I act “on impulse”?); and (3) non-planning impulsivity (e.g. I plan tasks carefully?). Participants rated each item on a 5-point Likert scale (1 = rarely/never, 5 = almost always/always), with higher scores indicating greater levels of impulsivity. Besides, we also use the Chinese version of BIS-11 (80) in the independent adult cohort to evaluate measurement consistency and generalizability across cultural and linguistic contexts.

#### General Procrastination Scale (GPS)

Procrastination was assessed using the 20-item General Procrastination Scale (GPS) (81), a widely used self-report measure developed to capture chronic tendencies to postpone intended actions despite potential negative consequences. The scale includes items such as “I generally delay before starting on work I have to do”, rated on a 5-point Likert scale, with higher scores indicating greater procrastinatory behavior.

### Extended Bayesian information criterion graphic lasso (EBICglasso) analysis

Given the potential influence of high-bridge-centrality items on the impulsivity–procrastination relationship, we conducted an Extended Bayesian information criterion graphic lasso (EBICglasso) analysis to identify bridging nodes connecting these phenotypes. Partial correlations were calculated across all items from the BIS and GPS scales to define the network’s edges. The network was then estimated using the EBICglasso algorithm, which penalizes spurious connections to enhance interpretability and control for false positives (82). A tuning parameter of 0.5 was selected to balance network sparsity and model fit, consistent with standard practices in network analysis. To identify key bridging symptoms, we calculated multiple centrality metrics, including 1-step and 2-step Bridge Expected Influence (BEI) and Bridge Strength, which quantify the extent to which nodes connect distinct symptom clusters. To address potential inconsistencies arising from the reliance on a single metric, these metrics were normalized and combined into a composite score to provide a comprehensive assessment of each node’s bridging role.

To ensure the robustness and replicability of our findings, we conducted permutation tests with 1,000 iterations. Due to the high degree of node isolation in the original network, bootstrap analyses were restricted to the fully connected network. To address the inherent instability and inconsistencies that may arise in networks that are not fully connected, we calculated Shannon entropy (SE) for each node. This metric captures the diversity and stability of its bridging influence, providing a more robust assessment of its role within the network. By focusing on nodes exceeding the 90% in Shannon entropy, we identified bridging symptoms that play a pivotal role in maintaining inter-cluster connectivity, thereby enhancing the interpretability and reliability of results in non-fully connected networks (83).

### ACE modeling

We used structural equation modeling implemented in the *OpenMx* package in *R* (84) to estimate the heritability of NPI and procrastination. Univariate ACE models were first applied to decompose the variance of each trait into additive genetic (A), shared environmental (C), and non-shared environmental (E) components. Heritability was defined as the proportion of variance attributable to genetic factors. We then employed bivariate ACE models to examine the genetic and environmental correlations between NPI and procrastination. These models allowed us to estimate the genetic correlation (*r*_*g*_), which reflects the extent of shared genetic influences between the two traits, as well as the phenotypic correlation (*r*_*ph*_), which captures the combined genetic and environmental covariance. For each model, we began with the full ACE specification and compared nested sub-models (AE, CE, and E) using likelihood-ratio tests, the Akaike Information Criterion (AIC), and the Bayesian Information Criterion (BIC) to identify the best-fitting model.

To investigate the role of individual morphological deviation in the genetic relationship between NPI and procrastination, we extended our analysis to multivariate modeling (85). A trivariate Cholesky decomposition was used to sequentially partition the covariance among morphological deviation, NPI and procrastination into A, C, and E components. Trait ordering was set as morphological deviation→ NPI→ procrastination, consistent with our hypothesis that procrastination emerges as a genetic byproduct of impulsivity, and their genetic associations may be underpinned by brain morphology. However, given the theoretical ambiguity regarding causal ordering, we also tested the alternative model specifying the sequence as NPI→ procrastination→ morphological deviation (***Supplementary Material S8)***. In this framework, the first set of latent factors (A1, C1, E1) captures influences on morphological deviation and potentially on the other two traits, while the second (A2, C2, E2) accounts for residual influences on NPI and morphological deviation, and the third (A3, C3, E3) represents unique variance in procrastination. We additionally tested alternative multivariate structures, including the Independent Pathway and Common Pathway models, which differ in how shared variance is modeled across traits **(*Supplementary Material* S4)** (86). Model comparison was conducted using −2 log-likelihood ratio tests and AIC values, and all models were fitted using the CSOLNP optimizer. Likelihood-based 95% confidence intervals were computed for all parameter estimates.

### Single-paper meta-analytic synthesis

To generate precise and stable estimates of effect sizes across multiple independent samples reported within the current study, we implemented single-paper meta-analytic synthesis ((87)). This approach treats each dataset within the manuscript as a separate but conceptually aligned study, and aggregates their effect sizes using meta-analytic techniques to improve statistical precision and mitigate the risk of overinterpreting sampling variability. Random-effects models were used under the assumption that the presence of both conceptual alignment and modest methodological heterogeneity across datasets.

In the first application, we examined the association between NPI and procrastination across three independent cohorts. Each study provided a standardized effect size (*Hedges’ g*) quantifying the relationship between NPI and procrastination. In the second application, we used single-paper meta-analysis to synthesize genetic correlations (*r*_*g*_) between impulsivity and procrastination, integrating four estimates—one derived from our twin sample using ACE modeling, and three from prior twin studies with matched phenotypic constructs. For both meta-analyses, these effect sizes were synthesized using a random-effect meta-analysis implemented in *R*. Post hoc power analyses were conducted to estimate the achieved power given the synthesized effect sizes and combined sample sizes. A priori power calculations were also conducted to determine required sample sizes for detecting the pooled effect with 80% power at α = .05. This analytic approach provides a principled framework for integrating evidence across sub-studies within a single manuscript and strengthens the reliability and reproducibility of the reported findings.

### Genome-wide association analysis

Whole-genome sequencing was performed on 936 samples from the adult cohort using the BGI Genomics at a mean sequencing depth of 27×. Genomic DNA was extracted from peripheral blood using the QIAGEN DNeasy Blood & Tissue Kit. Library preparation included random fragmentation via Covaries, size selection, adapter ligation, and amplification through ligation-mediated PCR. DNA nanoballs were generated via rolling circle amplification and sequenced using 150 bp paired-end reads. Raw sequencing images were processed using DNBSEQ’s base calling software with default parameters. Quality trimming of raw reads was conducted using SOAPnuke (v2.2.1) (88), and clean reads were aligned to the human reference genome (hg38) using the Burrows-Wheeler Aligner (89). Variant calling and annotation followed the Genome Analysis Toolkit (GATK, *v4*.*1*.*4*.*1*) (90) best practices, including base quality score recalibration, indel realignment, and variant quality score recalibration. The recalibration model was trained using four standard reference sets: HapMap3.3, dbSNP v150, 1000 Genomes Omni 2.5, and Phase I high-confidence SNPs. Variants were retained if they passed stringent quality thresholds: mean depth > 8×, Hardy–Weinberg equilibrium *P* > 1×10−^6^, and genotype call rate >90%. Samples were excluded if they had low sequencing depth (< 20×), excessive missingness, relatedness (Pi-hat > 0.1875), or population outlier status based on PCA in PLINK (*v1*.*9*). After quality control, the final sample included 835 unrelated individuals with 6.19 million common variants (minor allele frequency [MAF] ≥ 5%) and 3.64 million low-frequency variants (0.5% ≤ MAF < 5%). For more details, see ***Supplementary Material, Section S7*.*1***.

We conducted GWAS for both NPI and procrastination using linear regression models implemented in PLINK (v1.9). Trait scores were obtained from the non-planning subscale of the BIS-11 and GPS respectively. Each model controlled for sex, age, and the top 10 ancestry principal components to account for population stratification. Genome-wide significance was defined as *P* < 5 × 10^−5^. To further investigate the genetic basis of trait variation, we integrated GWAS summary statistics on DLPFC morphology obtained from the Chinese Imaging Genetics (CHIMGEN) study, which included 7,058 individuals of Han Chinese ancestry and analyzed 3,414 T1-weighted MRI-derived phenotypes (91). The imaging GWAS models were adjusted for age, sex, age*sex interaction, age-squared, and the top three ancestry principal components. Specifically, we focused on summary statistics for cortical surface area and thickness of the rostral middle frontal gyrus, derived from the Desikan–Killiany atlas via *FreeSurfer*. These publicly available summary statistics were retrieved from the GWAS Catalog (https://www.ebi.ac.uk/gwas) and served as a reference for follow-up analyses examining genetic convergence across impulsivity, procrastination, and brain morphology. For additional information, see ***Supplementary Material, Section S7*.*2***.

### Neuroanatomical loci calibration analysis

For impulsivity, we leveraged large-scale online meta-analytic synthesis via the *NeuroSynth* platform. Using the keywords “impulsivity”, “impulsive”, “impulsiveness”, and “impulse”, we retrieved uniformity test maps derived from a total of 198 peer-reviewed functional neuroimaging studies, encompassing 5,855 reported activation loci. These meta-analytic maps reflect the consistency of reported activation across studies referencing each term in their abstracts. With an expected false discovery rate (FDR) of 0.01, only the keywords “impulsivity” (k = 120) and “impulsive” (k = 78) yielded valid results; no studies met the inclusion threshold for “impulsiveness” or “impulse”. The two valid maps were binarized and combined using a voxel-wise union to generate a composite mask representing the core neuroanatomical correlates of impulsivity.

To identify brain regions reliably associated with procrastination, we conducted a systematic meta-analysis of structural neuroimaging studies following *PRISMA* guidelines (92). A comprehensive literature search was performed across *Web of Science, PsycINFO, and PubMed* using predefined morphological and neuroimaging-related keywords (Forward searching), supplemented by targeted author-based searching (Backward searching). This yielded 906 articles, of which 508 unique entries were screened using a semi-automated machine learning tool (*ASReview*). After full-text review and exclusion of studies not involving structural morphometric analysis of procrastination, five studies met all eligibility criteria and were included. Peak coordinates and effect sizes extracted from these studies were entered into Seed-based d Mapping (SDM, version 6.23) (93) to estimate voxel-wise associations with procrastination. The analysis was optimized for grey matter metrics, with preprocessing parameters including a 20 mm *FWHM* smoothing kernel and 2 mm voxel resolution. Statistical significance was determined using permutation-based family-wise error correction (*p* < 0.05, extent threshold ≥10 voxels). For additional information, see ***Supplementary Material, Section S5***.

To quantify the spatial overlap between impulsivity-procrastination volumetric mask, we first computed a volumetric conjunction mask derived from the meta-analytic synthesis of both traits. This volume-based mask was then projected onto the cortical surface using *FreeSurfer*’s *mri_vol2surf* utility, yielding hemisphere-specific surface representations. To enable anatomical localization within a standardized cortical parcellation, the resulting surface files were aligned to the DK atlas via *mri_surf2surf* utility, which resampled the surface data into fsaverage space. Subsequently, in *MATLAB*, we loaded the DK annotation files and assigned each vertex within the projected surface mask to its corresponding DK cortical region by referencing the annotation label table. This procedure allowed for the precise identification of DK-defined cortical areas exhibiting spatial overlap with the shared impulsivity–procrastination mask, thereby characterizing their putative surface-based neuroanatomical substrates.

### Normative modeling and neurodevelopmental deviation estimation

For the twin cohort, T1-weighted anatomical images were acquired using standardized 3T MRI protocols across all participants. Structural data were processed using the standard FreeSurfer pipeline (https://surfer.nmr.mgh.harvard.edu), which includes skull stripping, tissue segmentation, cortical surface reconstruction, and parcellation. Morphometric features—including regional cortical thickness (CT) and surface area (SA)—were extracted from 68 cortical regions defined by the DK atlas, along with global measures such as mean cortical thickness, total surface area.

To characterize neurodevelopmental deviations in cortical structure, we then implemented a normative modeling framework using the CentileBrain platform (https://centilebrain.org) (94). Normative reference models were pretrained on a large-scale, lifespan neuroimaging dataset curated by the ENIGMA Lifespan Working Group (N = 37,407; age range: 3–90 years), using multivariable fractional polynomial regression (MFPR). These models accounted for nonlinear age effects, sex, and site-specific variation harmonized via ComBat-GAM. Separate normative models were constructed for CT and SA, yielding expected values and variance estimates for each individual based on their demographic and scanner characteristics. The models enable the estimation of subject-level neuroanatomical deviations by comparing observed morphometric values to age- and sex-adjusted population norms.

To quantify individual-level deviations from normative trajectories, we then computed region-wise standardized deviation scores (*Z*) for each participant. These z-scores represent the degree of overdevelopment (*Z* > 0) or underdevelopment (*Z* < 0) relative to the expected value from the normative model, adjusted by the root mean square error (RMSE). Regional *z*-scores were averaged across participants to compute an unweighted Average Deviation Score (ADS), reflecting global patterns of deviation for CT and SA (Allen et al., 2024). As a supplementary analysis, we further classified statistically significant neurodevelopmental anomalies using a threshold of |*Z*| ≥ 1.96, corresponding to *p* < .05 (two-tailed) under a standard normal distribution, to identify morphometric deviations that exceeded normative expectations at the individual level (see ***Supplementary Material, Section S6***).

### Mendelian randomization analysis

We employed the *TwoSampleMR* package to investigate the causal relationships between NPI and DLPFC morphological traits, as well as between DLPFC morphological traits and procrastination. The primary analytical approach was the inverse-variance weighted (IVW) method. To validate the robustness of our findings, we conducted supplementary analyses using MR-Egger regression, weighted median, and weighted mode methods, each of which accounts for potential violations of instrumental variable assumptions in distinct ways. Heterogeneity among SNP-specific estimates was evaluated using Cochran’s *Q* test, and the MR-Egger intercept was used to detect horizontal pleiotropy. Sensitivity analyses, including leave-one-out analysis, were performed to identify influential instrumental variables by sequentially excluding each SNP and re-running the IVW analysis (see ***Supplementary Material, Section S7*.*3*)**. Effect estimates are presented as beta coefficients per 1 standard deviation increase in the exposure and its corresponding association with the outcome. Statistical significance was assessed using a Bonferroni-corrected threshold (*p* < 0.05). To ensure robust instrumental variable selection, we calculated *F-statistics* to evaluate the strength of the instruments and conducted statistical power analyses to confirm sufficient sensitivity for detecting causal effects.

## Acknowledgments

The authors gratefully acknowledge all the twin participants from the Beijing Twins Brain Behavior Association Project for their involvement in this study. We thank Ting Chen from the Institute of Psychology, Chinese Academy of Sciences, for her assistance with data collection. We also thank Jie Chen and Xinying Li from the Institute of Psychology, Chinese Academy of Sciences for their valuable suggestions on methodological design. This work was supported by the National Natural Science Foundation of China (32300907, 72033006), the Chongqing Natural Science Foundation (General Project), the PLA Talent Program Foundation (2022160258), and the AMU-RD Scholar Foundation (202211001).

## References

1. P. Steel, D. Taras, A. Ponak, J. Kammeyer-Mueller, Self-regulation of slippery deadlines: the role of procrastination in work performance. Frontiers in psychology 12, 783789 (2022).

2. R. Paulsen, Non-work at work: Resistance or what? Organization 22, 351–367 (2015).

3. J. Vveinhardt, W. Sroka, What determines employee procrastination and multitasking in the workplace: personal qualities or mismanagement? Journal of Business Economics and Management 23, 532–550 (2022).

4. American College Health Association, “American College Health Association National College Health Assessment III: Undergraduate Student Reference Group Executive Summary (Spring 2022)” (American College Health Association, 2022).

5. F. Johansson, et al., Associations between procrastination and subsequent health outcomes among university students in Sweden. JAMA network open 6, e2249346–e2249346 (2023).

6. J. R. Ferrari, C. A. Roster, Delaying Disposing: Examining the Relationship between Procrastination and Clutter across Generations. Curr Psychol 37, 426–431 (2018).

7. R. Gupta, D. A. Hershey, J. Gaur, Time Perspective and Procrastination in the Workplace: An Empirical Investigation. Curr Psychol 31, 195–211 (2012).

8. B. Nguyen, P. Steel, J. R. Ferrari, Procrastination’s Impact in the Workplace and the Workplace’s Impact on Procrastination. Int J Selection Assessment 21, 388–399 (2013).

9. P. Steel, D. Taras, A. Ponak, J. Kammeyer-Mueller, Self-regulation of slippery deadlines: the role of procrastination in work performance. Frontiers in Psychology 12, 783789 (2022).

10. F. Di Nocera, O. Ricciardi, G. Abate, A. Bevilacqua, Does the catechol-O-methyltransferase (COMT) Val158Met human polymorphism influence procrastination? Organisms. Journal of Biological Sciences 1, 27–36 (2017).

11. D. E. Gustavson, et al., Genetic and Environmental Associations Between Procrastination and Internalizing/Externalizing Psychopathology. Clinical Psychological Science 5, 798–815 (2017).

12. P. Steel, The nature of procrastination: a meta-analytic and theoretical review of quintessential self-regulatory failure. Psychological bulletin 133, 65 (2007).

13. D. E. Gustavson, A. Miyake, J. K. Hewitt, N. P. Friedman, Genetic Relations Among Procrastination, Impulsivity, and Goal-Management Ability: Implications for the Evolutionary Origin of Procrastination. Psychol Sci 25, 1178–1188 (2014).

14. D. E. Gustavson, A. Miyake, J. K. Hewitt, N. P. Friedman, Understanding the cognitive and genetic underpinnings of procrastination: Evidence for shared genetic influences with goal management and executive function abilities. Journal of Experimental Psychology: General 144, 1063 (2015).

15. F. Sirois, T. Pychyl, Procrastination and the Priority of Short-Term Mood Regulation: Consequences for Future Self. Social & Personality Psych 7, 115–127 (2013).

16. T. R. Zentall, Enhancing “self-control”: The paradoxical effect of delay of reinforcement. Learn Behav 48, 165–172 (2020).

17. P. Steel, The procrastination equation: How to stop putting things off and start getting stuff done (FT Press, 2012).

18. P. Steel, Arousal, avoidant and decisional procrastinators: Do they exist? Personality and individual differences 48, 926–934 (2010).

19. J. R. Ferrari, M. J. Olivette, Parental authority and the development of female dysfunctional procrastination. Journal of research in Personality 28, 87–100 (1994).

20. P. Liu, T. Feng, The overlapping brain region accounting for the relationship between procrastination and impulsivity: A voxel-based morphometry study. Neuroscience 360, 9–17 (2017).

21. P. Steel, The nature of procrastination: a meta-analytic and theoretical review of quintessential self-regulatory failure. Psychological bulletin 133, 65 (2007).

22. D. E. Gustavson, et al., Genetic and Environmental Associations Between Procrastination and Internalizing/Externalizing Psychopathology. Clinical Psychological Science 5, 798–815 (2017).

23. S. Ma, et al., Molecular and cellular evolution of the primate dorsolateral prefrontal cortex. Science 377, eabo7257 (2022).

24. T. M. Preuss, S. P. Wise, Evolution of prefrontal cortex. Neuropsychopharmacology 47, 3– 19 (2022).

25. F. A. Mansouri, E. Koechlin, M. G. Rosa, M. J. Buckley, Managing competing goals—a key role for the frontopolar cortex. Nature Reviews Neuroscience 18, 645–657 (2017).

26. Z. Chen, T. Feng, Neural connectome features of procrastination: Current progress and future direction. Brain and Cognition 161, 105882 (2022).

27. C. M. Garin, et al., An evolutionary gap in primate default mode network organization. Cell reports 39 (2022).

28. T. Xu, et al., Cross-species functional alignment reveals evolutionary hierarchy within the connectome. Neuroimage 223, 117346 (2020).

29. Y. Yin, et al., Exploring common and distinct neural basis of procrastination and impulsivity through elastic net regression. Cerebral Cortex 35, bhae503 (2025).

30. Z. Chen, P. Liu, C. Zhang, T. Feng, Brain morphological dynamics of procrastination: The crucial role of the self-control, emotional, and episodic prospection network. Cerebral Cortex 30, 2834–2853 (2020).

31. Y. Hu, P. Liu, Y. Guo, T. Feng, The neural substrates of procrastination: A voxel-based morphometry study. Brain and cognition 121, 11–16 (2018).

32. N. Pan, et al., Brain gray matter structures associated with trait impulsivity: A systematic review and voxel-based meta-analysis. Human Brain Mapping 42, 2214–2235 (2021).

33. B. Besteher, C. Gaser, I. Nenadić, Brain structure and trait impulsivity: A comparative VBM study contrasting neural correlates of traditional and alternative concepts in healthy subjects. Neuropsychologia 131, 139–147 (2019).

34. Z. Chen, T. Feng, Neural connectome features of procrastination: Current progress and future direction. Brain and Cognition 161, 105882 (2022).

35. E. H. Willbrand, et al., Variable Presence of an Evolutionarily New Brain Structure is Related to Trait Impulsivity. Biological Psychiatry: Cognitive Neuroscience and Neuroimaging (2024).

36. E. Morris, Editors’ Choice. Science 357 (2017).

37. J. C. Loehlin, N. G. Martin, The genetic correlation between procrastination and impulsivity. Twin research and human genetics 17, 512–515 (2014).

38. J. Fu, et al., Cross-ancestry genome-wide association studies of brain imaging phenotypes. Nature Genetics 56, 1110–1120 (2024).

39. D. C. Burk, B. B. Averbeck, Environmental uncertainty and the advantage of impulsive choice strategies. PLOS Computational Biology 19, e1010873 (2023).

40. J. Fenneman, W. E. Frankenhuis, P. M. Todd, In which environments is impulsive behavior adaptive? A cross-discipline review and integration of formal models. Psychological Bulletin 148, 555 (2022).

41. J. R. Stevens, D. W. Stephens, The adaptive nature of impulsivity. (2010).

42. P. Steel, Arousal, avoidant and decisional procrastinators: Do they exist? Personality and individual differences 48, 926–934 (2010).

43. P. Steel, The procrastination equation (Allen & Unwin, 2011).

44. P. Steel, C. J. König, Integrating Theories of Motivation. AMR 31, 889–913 (2006).

45. M. M. L. Rebetez, L. Rochat, C. Barsics, M. Van Der Linden, Procrastination as a Self-Regulation Failure: The Role of Impulsivity and Intrusive Thoughts. Psychol Rep 121, 26– 41 (2018).

46. C. Sümer, O. B. Büttner, I’ll do it–after one more scroll: the effects of boredom proneness, self-control, and impulsivity on online procrastination. Frontiers in Psychology 13, 918306 (2022).

47. J. D. Parker, R. M. Bagby, Impulsivity in adults: A critical review of measurement approaches. Impulsivity: Theory, assessment, and treatment 142–157 (1997).

48. J. C. Strickland, M. W. Johnson, Rejecting impulsivity as a psychological construct: A theoretical, empirical, and sociocultural argument. Psychological review 128, 336 (2021).

49. D. E. Gustavson, A. Miyake, J. K. Hewitt, N. P. Friedman, Genetic Relations Among Procrastination, Impulsivity, and Goal-Management Ability: Implications for the Evolutionary Origin of Procrastination. Psychol Sci 25, 1178–1188 (2014).

50. S. Chavanas, et al., Mutations in SPINK5, encoding a serine protease inhibitor, cause Netherton syndrome. Nature genetics 25, 141–142 (2000).

51. N. A. Le, et al., Regulation of serine protease inhibitor Kazal type-5 (SPINK5) gene expression in the keratinocytes. Environ Health Prev Med 19, 307–313 (2014).

52. C. J. Sumner, et al., A dominant mutation in FBXO38 causes distal spinal muscular atrophy with calf predominance. The American Journal of Human Genetics 93, 976–983 (2013).

53. D. Roedl, C. Traidl-Hoffmann, J. Ring, H. Behrendt, M. Braun-Falco, Serine protease inhibitor lymphoepithelial Kazal type-related inhibitor tends to be decreased in atopic dermatitis. (2009).

54. Q. Liu, et al., A functional polymorphism in the SPINK5 gene is associated with asthma in a Chinese Han Population. BMC Med Genet 10, 59 (2009).

55. A.P. Pérez-González, G. de Anda-Jáuregui, E. Hernández-Lemus, Differential Transcriptional Programs Reveal Modular Network Rearrangements Associated with Late-Onset Alzheimer’s Disease. International Journal of Molecular Sciences 26, 2361 (2025).

56. A. S. Rothmeier, W. Ruf, Protease-activated receptor 2 signaling in inflammation. Semin Immunopathol 34, 133–149 (2012).

57. A. Pegat, et al., Identification of rare variants in the FBXO38 gene of patients with chronic inflammatory demyelinating polyradiculoneuropathy. Journal of Neuroimmunology 392, 578381 (2024).

58. O. K. Ijomone, R. S. Oria, O. M. Ijomone, M. Aschner, J. Bornhorst, Dopaminergic Perturbation in the Aetiology of Neurodevelopmental Disorders. Mol Neurobiol 62, 2420– 2434 (2025).

59. P. P. Silveira, L. More, C. Gottfried, Gene and Environment Interactions in Neurodevelopmental Disorders. Frontiers in Behavioral Neuroscience 16, 893662 (2022).

60. L. T. Hunt, T. E. Behrens, T. Hosokawa, J. D. Wallis, S. W. Kennerley, Capturing the temporal evolution of choice across prefrontal cortex. Elife 4, e11945 (2015).

61. L. T. Hunt, T. E. Behrens, T. Hosokawa, J. D. Wallis, S. W. Kennerley, Capturing the temporal evolution of choice across prefrontal cortex. Elife 4, e11945 (2015).

62. S. Kim, D. Lee, Prefrontal cortex and impulsive decision making. Biological psychiatry 69, 1140–1146 (2011).

63. H. Raji, S. Dinesh, S. Sharma, Inside the impulsive brain: a narrative review on the role of neurobiological, hormonal and genetic factors influencing impulsivity in psychiatric disorders. Egypt J Neurol Psychiatry Neurosurg 61, 4 (2025).

64. G. Aydogan, et al., Genetic underpinnings of risky behaviour relate to altered neuroanatomy. Nature Human Behaviour 5, 787–794 (2021).

65. M. Kohno, et al., Functional genetic variation in dopamine signaling moderates prefrontal cortical activity during risky decision making. Neuropsychopharmacology 41, 695–703 (2016).

66. R. Le Bouc, M. Pessiglione, A neuro-computational account of procrastination behavior. Nature Communications 13, 5639 (2022).

67. T. R. Zentall, Basic behavioral processes involved in procrastination. Frontiers in Psychology 12, 769928 (2021).

68. T. Xu, F. M. Sirois, L. Zhang, Z. Yu, T. Feng, Neural basis responsible for self-control association with procrastination: Right MFC and bilateral OFC functional connectivity with left dlPFC. Journal of Research in Personality 91, 104064 (2021).

69. T. Xu, S. Zhang, F. Zhou, T. Feng, Stimulation of left dorsolateral prefrontal cortex enhances willingness for task completion by amplifying task outcome value. Journal of Experimental Psychology: General 152, 1122 (2023).

70. S. M. Kolk, P. Rakic, Development of prefrontal cortex. Neuropsychopharmacology 47, 41– 57 (2022).

71. R. E. Passingham, S. P. Wise, The neurobiology of the prefrontal cortex: anatomy, evolution, and the origin of insight (Oxford University Press (UK), 2012).

72. S. Zhang, P. Liu, T. Feng, To do it now or later: The cognitive mechanisms and neural substrates underlying procrastination. WIRES Cognitive Science 10, e1492 (2019).

73. S. Zhang, T. Feng, Modeling procrastination: Asymmetric decisions to act between the present and the future. Journal of Experimental Psychology: General 149, 311 (2020).

74. R. L. Bryck, P. A. Fisher, Training the brain: practical applications of neural plasticity from the intersection of cognitive neuroscience, developmental psychology, and prevention science. American psychologist 67, 87 (2012).

75. A. Gutchess, Plasticity of the aging brain: New directions in cognitive neuroscience. Science 346, 579–582 (2014).

76. D. Crews, Epigenetic modifications of brain and behavior: theory and practice. Hormones and behavior 59, 393–398 (2011).

77. H. Jeong, et al., Evolution of DNA methylation in the human brain. Nature communications 12, 2021 (2021).

78. E. B. Keverne, D. W. Pfaff, I. Tabansky, Epigenetic changes in the developing brain: Effects on behavior. Proc. Natl. Acad. Sci. U.S.A. 112, 6789–6795 (2015).

79. J. H. Patton, M. S. Stanford, E. S. Barratt, Factor structure of the barratt impulsiveness scale. J. Clin. Psychol. 51, 768–774 (1995).

80. L. Zhou, S. Xiao, X. He, J. Li, H. Liu, Reliability and validity of the Chinese version of the BIS-Chinese Journal of Clinical Psychology 14, 343–344 (2006).

81. C. H. Lay, At last, my research article on procrastination. Journal of research in personality 20, 474–495 (1986).

82. D. Borsboom, A. O. J. Cramer, Network Analysis: An Integrative Approach to the Structure of Psychopathology. Annu. Rev. Clin. Psychol. 9, 91–121 (2013).

83. Z. Chen, et al., Edge-centric connectome-genetic markers of bridging factor to comorbidity between depression and anxiety. Nature Communications 15, 1–15 (2024).

84. S. Boker, et al., OpenMx: an open source extended structural equation modeling framework. Psychometrika 76, 306–317 (2011).

85. C. A. M. Sutherland, et al., Individual differences in trust evaluations are shaped mostly by environments, not genes. Proc. Natl. Acad. Sci. U.S.A. 117, 10218–10224 (2020).

86. S. Clifford, K. Lemery-Chalfant, H. H. Goldsmith, The Unique and Shared Genetic and Environmental Contributions to Fear, Anger, and Sadness in Childhood. Child Development 86, 1538–1556 (2015).

87. B. B. McShane, U. Böckenholt, Single-paper meta-analysis: Benefits for study summary, theory testing, and replicability. Journal of Consumer Research 43, 1048–1063 (2017).

88. Y. Chen, et al., SOAPnuke: a MapReduce acceleration-supported software for integrated quality control and preprocessing of high-throughput sequencing data. Gigascience 7, gix120 (2018).

89. H. Li, R. Durbin, Fast and accurate short read alignment with Burrows–Wheeler transform. bioinformatics 25, 1754–1760 (2009).

90. M. A. DePristo, et al., A framework for variation discovery and genotyping using next-generation DNA sequencing data. Nature genetics 43, 491–498 (2011).

91. J. Fu, et al., Cross-ancestry genome-wide association studies of brain imaging phenotypes. Nature Genetics 1–11 (2024).

92. M. J. Page, et al., The PRISMA 2020 statement: an updated guideline for reporting systematic reviews. Bmj 372 (2021).

93. J. Radua, D. Mataix-Cols, Voxel-wise meta-analysis of grey matter changes in obsessive– compulsive disorder. The British Journal of Psychiatry 195, 393–402 (2009).

94. R. Ge, et al., Normative modelling of brain morphometry across the lifespan with CentileBrain: algorithm benchmarking and model optimisation. The Lancet Digital Health 6, e211– e221 (2024).

