## Supplementary Materials for "Procrastination partly reflects an evolutionary byproduct of non-planning impulsivity"

**This PDF file includes:**

Supporting text
Figures S1 to S8
Tables S1.1 to S7.5
SI References

**Supporting Information Text**

S2 Table S2: Extended Bayesian Information Criterion (EBIC) glasso network of Items in the GPS and the No-Planning Dimension of the BIS in the Twin Cohort. . 14

**Table S1:** Participant Demographics by Dataset.

| dataset | age (Mean ±SD) | Gender (female/male) | MZ | DZ | N |
|---|---|---|---|---|---|
| Longitudinal Twin Cohort | 19.61±2.43 | 86/68 | 40 pairs | 37 pairs | N= 154 (77 pairs) |
| Cross-sectional Adolescent Cohort | 20.04±1.68 | 92/235 | - | - | N= 327 |
| Cross-sectional Large-scale Adult Cohort | 18.45±1.34 | 1041/502 | - | - | N= 1,543 |
| GWAS dataset | 18.93±0.95 | 623/313 | - | - | N= 936 |
| ENIGMA lifespan sample | 3-90 years | 53%/47% | - | - | N= 37,407 |
| CHIMGEN-GWAS dataset | 23.68±2.48 | 4557/2501 | - | - | N= 7,058 |

#### S1 Comprehensive Analyses of Impulsivity Dimensions and Their Network Interrelations in Predicting Procrastination

#### S1.1 Dimensional Structure and Interrelations of BIS Subdimensions in Predicting Procrastination

To investigate whether impulsivity can prospectively predict procrastination nine years later, we first examined the relationships among the three BIS subdimensions (Attentional, Motor, and Non-planning impulsivity) and the total BIS score. This analysis aimed to clarify how each subdimension uniquely contributes to the overarching construct of impulsivity while also capturing their shared variance. **Pearson's correlation coefficients** were computed, revealing significant correlations between the BIS total score and each subdimension across all three cohorts, as well as intercorrelations among the subdimensions (**Figure S1**). These findings underscore the multifaceted nature of impulsivity, where distinct dimensions are interrelated yet retain unique contributions.

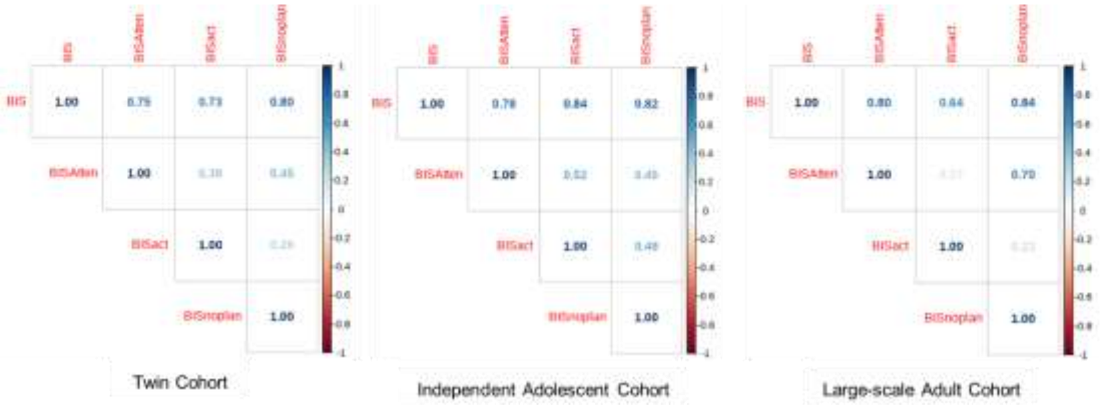

**Figure S1** Correlations Between BIS Total Score and Subdimensions Across Three Cohorts

To further evaluate the dimensional structure of the BIS, we conducted **principal component analysis (PCA)** across all three cohorts. The analysis was performed without rotation to examine whether the data naturally supported the theoretical three-factor structure. If three components with eigenvalues greater than 1 were extracted and each component corresponded closely to one of the BIS subdimensions—attentional, motor, and non-planning impulsivity—this would suggest that the subdimensions are both distinguishable and relatively independent. Indeed, the unrotated PCA consistently yielded three components across cohorts. However, the alignment between the extracted components and the theoretical subdimensions was not exact, as several items showed substantial cross-loadings across components. This pattern indicates that while the overall three-factor structure is recoverable (**Figure S2**), the components are not strictly orthogonal, supporting the interpretation that the BIS subdimensions reflect overlapping, obliquely related constructs.

In conclusion, the data exploration results support the presence of three primary dimensions of impulsivity, as predicted by the BIS framework. However, the observed oblique relationships highlight that these subdimensions are not entirely independent, underscoring the nuanced interplay between different aspects of impulsivity. Thus, controlling for the effects of the other impulsivity subdimensions was necessary to ensure clearer phenotypic associations when using each subdimension to predict procrastination level longitudinally.

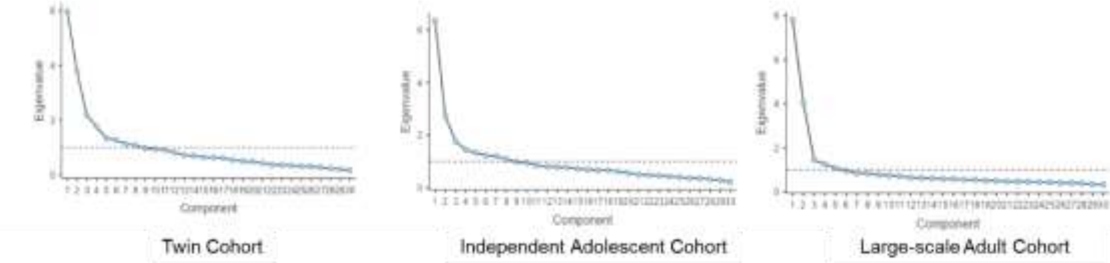

**Figure S2** Scree plot of principal component analysis on impulsivity subdimensions.

#### S1.2 Investigating the Predictive Role of Impulsivity Subdimensions Across Populations and Measurement Tools

In this study, we investigated the unique contributions of impulsivity subdimensions to procrastination across three distinct cohorts while accounting for key demographic covariates. For the 9-year longitudinal Twins Cohort, we addressed potential multicollinearity among the impulsivity subdimensions by orthogonalizing each subdimension. This was achieved by calculating residual scores for each subdimension through linear regression on the other two, ensuring that the residualized subdimensions captured unique, non-overlapping variance. We then employed **linear mixed-effects models (LMMs)**, incorporating family structure (familyID) as a random effect, to account for within-family dependencies and robustly estimate the predictive relationships between the orthogonalized subdimensions and procrastination.

Extending this framework, we conducted analogous analyses in an adolescent cohort and an independent adult cohort respectively to further evaluate the robustness of our findings across diverse populations and measurement tools. We first applied a similar orthogonalization process to the independent adolescent dataset, calculating residual scores for each impulsivity subdimension. Unlike the Twins Cohort, the independent adolescent cohort lacked hierarchical dependencies. In this case, **linear regression models (LMs)** were used to evaluate the unique contributions of each orthogonalized subdimension to procrastination, with gender and age included as covariates. The same orthogonalization process was performed to isolate the unique contributions of each impulsivity subdimension, followed by **LMs** assessing their relationship with procrastination in the independent adult cohort. Notably, unlike the Twins Cohort and adolescent cohort, which used the Barratt Impulsiveness Scale (BIS-11), the independent adult cohort employed the Chinese version of BIS-11, allowing us to evaluate measurement consistency and generalizability across cultural and linguistic contexts.

Across all three datasets, model comparison results consistently identified Non-planning Impulsivity (NPI), significantly outperformed other impulsivity subdimensions, such as Attentional Impulsivity and Motor Impulsivity, as best-fitting predictor of procrastination (S1 Table S1). Even after removing items with high bridge centrality in each dataset to control between-scale psychometric similarity, we obtained similar results (S1 Table S2, Figure 3). This finding underscores the unique and robust relationship between NPI and procrastination, highlighting its critical role in explaining procrastination behaviors. By systematically applying this methodology across three cohorts, we ensured that our findings are replicable and generalizable across developmental stages, population heterogeneity, and measurement tools. This comprehensive approach highlights not only the consistent predictive role of impulsivity subdimensions but also the robustness of the BIS framework across cultural adaptations.

**S1 Table S1:** Model comparison and predictive performance of impulsivity subdimensions on procrastination **before** removing high centrality items across three cohorts.

| dataset | Name | AIC (weights) | AIC$_C$ (weights) | BIC (weights) | R2 (cond.) | $R^2$ (marg.) | ICC | RMSE | Sigma |
|---|---|---|---|---|---|---|---|---|---|
| Longitudinal Twin Adolescent Cohort | GPS~ BISAtten | 1149.9 (0.572) | 1150.7 (0.572) | 1171.2 (0.572) | 0.440 | 0.111 | 0.370 | 6.835 | 8.096 |
| | GPS~ BISact | 1150.6 (0.398) | 1151.4 (0.398) | 1171.9 (0.398) | 0.384 | 0.108 | 0.309 | 7.252 | 8.401 |
| | GPS~ BISnon-planning | 1155.8 (0.030) | 1156.6 (0.030) | 1177.1 (0.030) | 0.406 | 0.079 | 0.355 | 7.065 | 8.324 |

| dataset | Name | AIC (weights) | AIC$_C$ (weights) | BIC (weights) | R2 | $R^2$ (adj.) | RMSE | Sigma |
|---|---|---|---|---|---|---|---|---|
| Cross-sectional Adolescent Cohort | GPS~ BISAtten | 2516.5 (0.946) | 2516.7 (0.946) | 2534.4 (0.946) | 0.082 | 0.074 | 11.174 | 11.243 |
| | GPS~ BISact | 2522.2 (0.054) | 2522.4 (0.054) | 2541.2 (0.054) | 0.066 | 0.057 | 11.273 | 11.342 |
| | GPS~ BISnon-planning | 2537.5 (<0.001) | 2537.7 (<0.001) | 2556.5 (<0.001) | 0.021 | 0.012 | 11.539 | 11. 611 |
| Cross-sectional Large-scale Adult Cohort | GPS~ BISAtten | 11783.0 (>0.999) | 11783.0 (>0.999) | 11809.7 (>0.999) | 0.108 | 0.106 | 10.979 | 10.994 |
| | GPS~ BISact | 11957.5 (<0.001) | 11957.5 (<0.001) | 11984.2 (<0.001) | 0.001 | <0.001 | 11.618 | 11.633 |
| | GPS~ BISnon-planning | 11894.1 (<0.001) | 11894.1 (<0.001) | 11920.8 (<0.001) | 0.042 | 0.040 | 11.382 | 11.397 |

**S1 Table S2:** Model comparison and predictive performance of impulsivity subdimensions on procrastination **after** removing high centrality items across three cohorts.

| dataset | Name | AIC (weights) | $AIC_C$ (weights) | BIC (weights) | R2 (cond.) | $R^2$ (marg.) | ICC | RMSE | Sigma |
|---|---|---|---|---|---|---|---|---|---|
| Longitudinal Twin Adolescent Cohort | GPS_del~ BISAtten | 1129.7 (0.769) | 1130.4 (0.769) | 1150.9 (0.769) | 0.417 | 0.124 | 0.335 | 6.610 | 7.733 |
| | GPS_del ~ BISact | 1136.0 (0.033) | 1136.7 (0.033) | 1157.2 (0.033) | 0.432 | 0.089 | 0.377 | 6.493 | 7.708 |
| | GPS_del ~ BISnon-planning_del | 1132.4 (0.198) | 1133.1 (0.198) | 1153.6 (0.198) | 0.445 | 0.108 | 0.377 | 6.413 | 7.615 |
| **dataset** | **Name** | **AIC (weights)** | **$AIC_C$ (weights)** | **BIC (weights)** | **R2** | **$R^2$ (adj.)** | | **RMSE** | **Sigma** |
| Cross-sectional Adolescent Cohort | GPS_del ~ BISAtten | 2478.2 (0.937) | 2478.4 (0.937) | 2497.1 (0.937) | 0.071 | 0.063 | | 10.538 | 10.603 |
| | GPS_del ~ BISact | 2494.5 (<.001) | 2494.7 (<.001) | 2513.4 (<.001) | 0.024 | 0.015 | | 10.804 | 10.871 |
| | GPS_del ~ BISnon-planning_del | 2483.6 (0.062) | 2483.8 (0.062) | 2502.5 (0.062) | 0.056 | 0.047 | | 10.626 | 10.692 |
| Cross-sectional Large-scale Adult Cohort | GPS_del ~ BISAtten | 11844.9 (< .001) | 11844.9 (< .001) | 11871.6 (< .001) | < .001 | -0.02 | | 11.202 | 11.216 |
| | GPS_del ~ BISact | 11768.3 (< .001) | 11768.4 (< .001) | 11795.0 (< .001) | 0.048 | 0.047 | | 10.927 | 10.942 |
| | GPS_del ~ BISnon-planning_del | 11729.8 (> .999) | 11729.9 (> .999) | 11756.5 (> .999) | 0.072 | 0.070 | | 10.792 | 10.806 |

**S1 Table S3:** Single Meta-Analytic Indicators for the Prediction of Procrastination by Non-Planning Impulsivity Before and After Removing High Betweenness Centrality Items Across Datasets.

| **Dataset** | **Before/after removing high-bridge items** | ***β*** | ***SE*** | ***t value*** | ***p*** |
|---|---|---|---|---|---|
| Longitudinal Twin Adolescent Cohort | before | 0.38 | 0.15 | 2.44 | 0.016 |
| | after | 0.34 | 0.18 | 1.89 | 0.060 |
| Cross-sectional Adolescent Cohort | before | 0.89 | 0.18 | 4.96 | < 0.001 |
| | after | 0.81 | 0.21 | 3.96 | < 0.001 |
| Cross-sectional Large-scale Adult Cohort | before | 0.40 | 0.03 | 13.66 | < 0.001 |
| | after | 0.95 | 0.09 | 10.92 | < 0.001 |

### S2 Full Findings to Network Analysis

Given the heterogeneity across populations, we conducted separate network analyses for each of the three datasets—the twin cohort, the cross-sectional adolescent cohort, and the large-scale adult cohort. Furthermore, to account for differences in measurement tools, we performed distinct network analyses for participants assessed with the BIS-11 (the twin cohort and the cross-sectional adolescent cohort) and those assessed with the Chinese version of BIS-11(the cross-sectional adolescent cohort), ensuring methodological rigor and consistency in interpreting the finding. Items surpassing the 90th percentile threshold were identified as exhibiting high bridge centrality, reflecting their critical role in bridging key constructs. These items highlight their pivotal role in bridging the relationship between impulsivity and procrastination, offering valuable insights into the interconnected dynamics of these constructs.

Network analyses based on BIS-11 are summarized in Figure S3. For the twin cohort, the Shannon entropy values for key items are as follows: "*I get easily bored when solving thought problems?*" (*SE* = 0.28) and "*I like puzzles?*" (*SE* = 0.30) from the BIS, along with "*I am continually saying I'll do it tomorrow.*" (*SE* = 0.28) from the GPS. S2 Table S**1** presents the corresponding 1-stage and 2-stage Bridge Expected Influence (BEI) values and Shannon entropy for each item in the GPS and the No-Planning subscale of the BIS-11, while S2 Table S**2** presents the EBICglasso network. For the cross-sectional adolescent cohort, the Shannon entropy values for key items are as follows: "*I plan tasks carefully?*" (*SE* = 0.33) and "*I am self controlled?*" (*SE* = 0.29) from the BIS, along with "I am continually saying I'll do it tomorrow" (*SE* = 0.30) from the GPS. For the combined dataset of the twin cohort and the cross-sectional adolescent cohort, the Shannon entropy values for key items are as follows: " *I get easily bored when solving thought problems?*" (*SE* = 0.29) and "*I am self controlled?*" (*SE* = 0.34) from the BIS, along with "I am continually saying I'll do it tomorrow" (*SE* = 0.32) from the GPS.

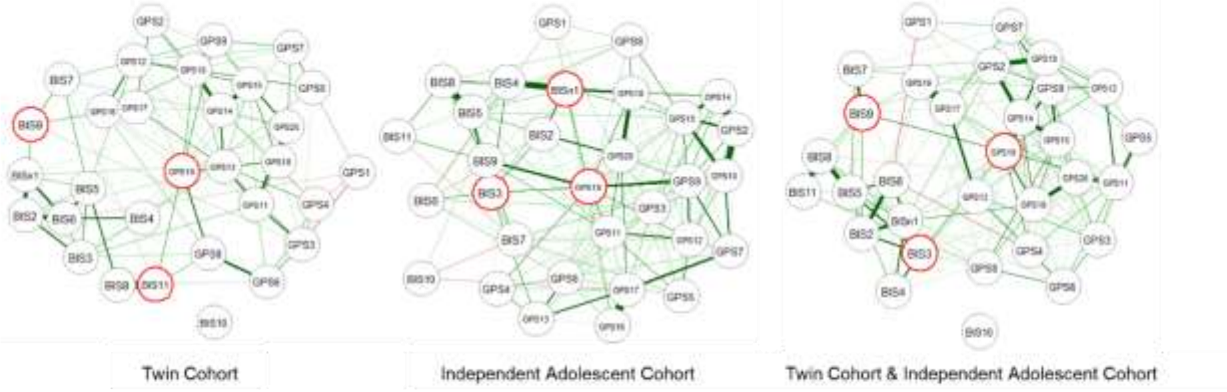

**Figure S3** Network Analyses Based on BIS-11: Results from the Twin Cohort, Cross-Sectional Adolescent Cohort, and Their Combined Dataset

Network analyses based on the Chinese version of BIS-11 are summarized in Figure S4. For the cross-sectional adolescent cohort, the Shannon entropy values for key items are as follows: "*我有规律地安排饮食起居? (I regularly schedule my daily routine.)*" (*SE* = 0.28) and "*我做事时能按计划完成 (I can complete tasks according to plan)?*" (*SE* = 0.36) from the BIS, along with " *I usually accomplish all the things I plan to do in a day.*" (*SE* = 0.26) from the GPS.

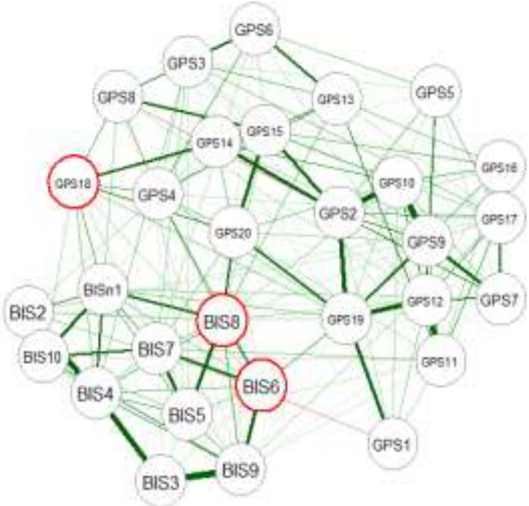

**Figure S4** Network Analyses Based on the Chinese version of BIS-11: Results from the cross-sectional adolescent cohort.

Although the network analysis results were somewhat inconsistent due to population heterogeneity and the heterogeneity of measurement tools, and the network model exhibited a high degree of node isolation, caution is required when interpreting the results. We prioritized the results derived from the twin cohort for subsequent analyses, as the imaging data were sourced from this dataset. Furthermore, in datasets measured using the BIS-11 and GPS, the item "*I am continually saying I'll do it tomorrow*" from the GPS consistently exhibited high bridge centrality, demonstrating its stable presence across different datasets.

After removing three items with high bridge centrality from the twin cohort, we conducted another PCA on the remaining GPS and NPI subscale items, using oblimin rotation and specifying a fixed number of two components to reflect the two hypothesized constructs. The results revealed a substantially clearer separation between the two factor structures: the remaining GPS items loaded predominantly on Component 1, while the BIS items loaded primarily on Component 2, with minimal cross-loadings and reduced overlap between components (**S2 Table S3**). This finding indicates that the psychometric distinction between procrastination and impulsivity was notably enhanced after item removal, thereby validating our decision to refine the measurement structure prior to downstream analyses.

**S2 Table S1:** Bridge Expected Influence and Shannon Entropy of Items in the GPS and the No-Planning Dimension of the BIS in the Twin Cohort.

| | Questions | 1-stage BEI | 2-stage BEI | Shannon Entropy |
|---|---|---|---|---|
| BIS11 | I like puzzles? | 0.198597528 | 0.340441419 | 0.302661528 |
| GPS19 | I am continually saying “I’ll do it tomorrow”. | 0.183449079 | 0.320589809 | 0.283388392 |
| BIS9 | I get easily bored when solving thought problems? | 0.172650356 | 0.325296565 | 0.275422919 |
| BIS4 | I save regularly? | 0.170030917 | 0.290644506 | 0.263772596 |
| BIS3 | I am self-controlled? | 0.159551298 | 0.313619881 | 0.260173906 |
| BIS7 | I say things without thinking? | 0.14516045 | 0.309117081 | 0.244020119 |
| GPS8 | I usually make decisions as soon as possible. | 0.137214691 | 0.249235985 | 0.21378424 |
| GPS13 | I prefer to leave early for an appointment. | 0.100854473 | 0.190223288 | 0.127387022 |
| BIS5 | I am a careful thinker? | 0.084642018 | 0.187295138 | 0.089360393 |
| GPS9 | I generally delay before starting on work I have to do. | 0.078981383 | 0.136706928 | 0.028188347 |
| GPS12 | In preparing for some deadline, I often waste time by doing other things. | 0.074460914 | 0.163026766 | 0.039749086 |
| GPS11 | When preparing to go out, I am seldom caught having to do something at the last minute. | 0.074336251 | 0.167863664 | 0.044110365 |
| GPS16 | I always seem to end up shopping for birthday or Christmas gifts at the last minute. | 0.06903215 | 0.141546385 | 0 |
| BISn1 | I plan tasks carefully? | 0.06281498 | 0.129236351 | 0 |
| GPS5 | A letter may sit for days after I write it before mailing it. | 0.061165409 | 0.082917318 | 0 |
| GPS20 | I usually take care of all the tasks I have to do before I settle down and relax for the evening. | 0.052069436 | 0.116617032 | 0 |
| GPS17 | I usually buy even an essential item at the last minute. | 0.046786915 | 0.112158112 | 0 |
| GPS18 | I usually accomplish all the things I plan to do in a day. | 0.045598015 | 0.119994291 | 0 |
| GPS3 | When I am finished with a library book, I return it right away regardless of the date it is due. | 0.042575115 | 0.088085382 | 0 |

| | | | | |
|---|---|---|---|---|
| BIS2 | I plan trips well ahead of time? | 0.040086599 | 0.146004531 | 0 |
| GPS6 | I generally return phone calls promptly. | 0.037726156 | 0.099392663 | 0 |
| GPS14 | I usually start an assignment shortly after it is assigned. | 0.027758583 | 0.082961352 | 0 |
| BIS8 | I like to think about complex problems? | 0.026411426 | 0.137833446 | 0 |
| GPS7 | Even with jobs that require little else except sitting down and doing them, I find they seldom get done for days. | 0.025467983 | 0.067793166 | 0 |
| GPS2 | I do not do assignments until just before they are to be handed in. | 0.009776734 | 0.023403235 | 0 |
| GPS15 | I often have a task finished sooner than necessary. | 0.003233903 | 0.059988156 | 0 |
| GPS4 | When it is time to get up in the morning, I most often get right out of bed. | 0 | 0.0265501 | 0 |
| GPS10 | I usually have to rush to complete a task on time. | 0 | 0.088073876 | 0 |
| BIS10 | I am interested in the present than the future? | 0 | 0 | 0 |
| BIS6 | I plan for job security? | 0.021686709 | 0.088192987 | 0 |
| GPS1 | I often find myself performing tasks that I had intended to do days before. | 0.032228327 | -0.069445604 | 0 |

**S2 Table S2:** Extended Bayesian Information Criterion (EBIC) glasso network of Items in the GPS and the No-Planning Dimension of the BIS in the Twin Cohort.

| | GPS 1 | GPS 2 | GPS 3 | GPS 4 | GPS 5 | GPS 6 | GPS 7 | GPS 8 | GPS 9 | GPS 10 | GPS 11 | GPS 12 | GPS 13 | GPS 14 | GPS 15 | GPS 16 | GPS 17 | GPS 18 | GPS 19 | GPS 20 | BISn1 | BIS2 | BIS3 | BIS4 | BIS5 | BIS6 | BIS7 | BIS8 | BIS9 | BIS10 | BIS11 |
|---|---|---|---|---|---|---|---|---|---|---|---|---|---|---|---|---|---|---|---|---|---|---|---|---|---|---|---|---|---|---|---|
| **GPS 1** | 0 | 0.040052931 | -0.156200121 | -0.096848394 | 0 | 0 | 0.147073685 | 0 | 0 | 0.004639859 | -0.01141982 | 0.01509562 | -0.006499323 | 0 | 0 | 0.006584744 | 0 | 0 | 0 | -0.031367711 | 0 | 0 | 0 | 0 | -0.011849901 | -0.058188396 | -0.022159898 | 0 | 0 | 0 | -0.03219404 |
| **GPS 2** | 0.040052931 | 0 | 0 | 0 | 0.054426195 | 0 | 0 | 0 | 0.01774074 | 0.153663608 | 0 | 0 | 0 | 0.118857794 | 0 | 0 | 0.141889069 | 0 | 0 | 0 | 0 | 0.001070892 | 0.017049228 | 0 | 0 | 0 | 0 | 0 | 0.038320801 | 0 | -0.077451364 |
| **GPS 3** | -0.156200121 | 0 | 0 | 0.092891827 | 0.060586731 | 0.166957732 | 0 | 0 | 0 | 0 | 0.183735126 | 0 | 0.093933554 | 0 | 0 | 0 | 0 | 0.04400901 | 0 | 0 | 0 | 0 | 0 | 0 | 0 | 0 | 0 | -0.040166087 | 0.067805827 | 0 | 0 |
| **GPS 4** | -0.096848394 | 0 | 0.092891827 | 0 | 0 | 0.139928674 | 0 | 0.004347825 | 0 | 0 | 0 | 0.022787275 | 0.133303813 | 0.135822699 | 0 | 0 | 0 | 0 | 0 | 0 | 0 | 0 | 0 | 0 | 0 | 0 | 0 | 0 | 0 | 0 | 0 |
| **GPS 5** | 0 | 0.054426195 | 0.060586731 | 0 | 0 | 0 | 0.201389588 | 0 | 0 | 0.166005841 | 0.017853922 | 0 | 0 | 0 | -0.087454574 | 0 | 0 | 0 | 0 | 0 | 0 | 0 | 0.011196092 | 0.101699554 | 0 | 0.006130292 | -0.027533389 | -0.068423947 | 0 | 0 | 0 |
| **GPS 6** | 0 | 0 | 0.166957732 | 0.139928674 | 0 | 0 | 0 | 0.237751988 | -0.060524714 | 0 | 0.083413258 | 0 | 0 | 0 | 0 | 0 | 0 | 0 | 0 | 0 | 0 | 0.005597996 | 0 | 0 | 0.008691288 | 0 | 0 | 0.073101979 | 0 | -0.035997055 | 0 |
| **GPS 7** | 0.147073685 | 0 | 0 | 0 | 0.201389588 | 0 | 0 | 0 | 0.120761939 | 0.022955455 | 0.020593156 | 0.063747545 | 0 | 0 | 0.052558257 | 0 | 0 | 0 | 0.048894843 | 0.116895063 | 0 | 0 | 0 | 0.043035594 | 0 | 0 | 0 | -0.000118229 | 0 | 0 | 0 |
| **GPS** | 0 | 0 | 0 | 0.0043 | 0 | 0.2377 | 0 | 0 | 0 | 0 | 0 | 0 | 0 | 0 | 0 | 0.0835 | 0 | 0 | 0.1752 | 0.0189 | 0 | 0 | 0.06585 | 0 | 0 | 0 | 0 | 0 | 0.00698 | 0 | 0.10589 |

| | | | | | | | | | | | | | | | | | | | | | | | | | | | | | | | |
|---|---|---|---|---|---|---|---|---|---|---|---|---|---|---|---|---|---|---|---|---|---|---|---|---|---|---|---|---|---|---|---|
| **8** | | | | 47825 | | 51988 | | | | | | | | | | 11555 | | | 47477 | 94423 | | | 3196 | | | | | | 7691 | | 2895 |
| **GPS 9** | 0 | 0.01774074 | 0 | 0 | 0 | -0.060524714 | 0.120761939 | 0 | 0 | 0.349303569 | 0 | 0.035883345 | 0 | 0 | 0.114946938 | 0 | 0 | 0.00208612 | 0.128468369 | 0.092376214 | 0 | -0.012562609 | 0.015335534 | 0 | 0 | 0 | 0.090348678 | 0 | 0 | 0 | 0 |
| **GPS 10** | 0.004639859 | 0.153663608 | 0 | 0 | 0.166005841 | 0 | 0.022955455 | 0 | 0.349303569 | 0 | 0 | 0.121138888 | 0.073076731 | 0.269034495 | 0 | 0.116082045 | 0 | 0 | 0.104306409 | 0 | 0 | 0 | 0 | 0 | 0 | 0 | 0 | 0 | 0 | 0 | 0 |
| **GPS 11** | -0.01141982 | 0 | 0.183735126 | 0 | 0.017853922 | 0.083413258 | 0.020593156 | 0 | 0 | 0 | 0 | 0.042871628 | 0.173162758 | 0 | 0.035894473 | 0.035840494 | 0.029369488 | 0.126840743 | 0 | 0.21798092 | 0 | 0.001918385 | 0.07738989 | 0 | 0 | 0 | 0 | 0 | 0.003061468 | 0 | 0 |
| **GPS 12** | 0.01509562 | 0 | 0 | 0.022787275 | 0 | 0 | 0.063747545 | 0 | 0.035883345 | 0.121138888 | 0.042871628 | 0 | 0 | 0 | 0.140598907 | 0.178736151 | 0 | 0 | 0.042278097 | 0 | 0 | 0 | 0 | 0 | 0.093727917 | 0 | 0 | 0 | 0 | 0 | 0 |
| **GPS 13** | -0.006499323 | 0 | 0.093933554 | 0.133303813 | 0 | 0 | 0 | 0 | 0 | 0.073076731 | 0.173162758 | 0 | 0 | 0.177725798 | 0.065331765 | 0 | 0.143564525 | 0.019737085 | 0 | 0.019666335 | 0 | 0.043993146 | 0 | 0.071112967 | 0 | 0 | 0 | 0 | 0 | 0 | 0 |
| **GPS 14** | 0 | 0.118857794 | 0 | 0.135822699 | 0 | 0 | 0 | 0 | 0 | 0.269034495 | 0 | 0 | 0.177725798 | 0 | 0.326009411 | 0 | 0.040873272 | 0.02011743 | 0.036613238 | 0.047129074 | 0.041034318 | 0 | 0 | 0 | 0 | 0.004818289 | 0 | 0 | 0 | 0 | 0 |
| **GPS 15** | 0 | 0 | 0 | 0 | -0.087454574 | 0 | 0.052558257 | 0 | 0.114946938 | 0 | 0.035894473 | 0.140598907 | 0.065331765 | 0.326009411 | 0 | 0.016298171 | 0 | 0.15104931 | 0 | 0.192811109 | 0 | 0 | 0 | 0 | 0 | 0 | 0.003951918 | 0 | 0 | 0 | 0.016422345 |
| **GPS 16** | 0.006584744 | 0 | 0 | 0 | 0 | 0 | 0 | 0.083511555 | 0 | 0.116082045 | 0.035840494 | 0.178736151 | 0 | 0 | 0.016298171 | 0 | 0.516183326 | 0 | 0.039318191 | 0 | -0.061834116 | 0 | 0 | 0 | 0 | 0 | 0 | 0 | 0.082955495 | 0 | 0 |
| **GPS 17** | 0 | 0.141889069 | 0 | 0 | 0 | 0 | 0 | 0 | 0 | 0 | 0.029369488 | 0 | 0.143564525 | 0.040873272 | 0 | 0.516183326 | 0 | 0 | 0.011380302 | 0 | 0 | 0 | 0 | 0 | 0 | 0 | 0 | 0 | 0.051888832 | 0 | 0 |
| **GPS** | 0 | 0 | 0.0440 | 0 | 0 | 0 | 0 | 0 | 0.0020 | 0 | 0.1268 | 0 | 0.0197 | 0.0201 | 0.1510 | 0 | 0 | 0 | 0.1499 | 0.1916 | 0.06 | 0 | 0 | 0 | 0.00 | 0 | 0 | 0 | 0 | - | 0 |

| | | | | | | | | | | | | | | | | | | | | | | | | | | | | | | | |
|---|---|---|---|---|---|---|---|---|---|---|---|---|---|---|---|---|---|---|---|---|---|---|---|---|---|---|---|---|---|---|---|
| 18 | | | 0901 | | | | | | 8612 | | 40743 | | 37085 | 1743 | 4931 | | | | 33045 | 42921 | 656256 | | | | 0583947 | | | | | 0.033632178 | |
| GPS19 | 0 | 0 | 0 | 0 | 0 | 0 | 0.048894843 | 0.175247477 | 0.128468369 | 0.104306409 | 0 | 0.042278097 | 0 | 0.036613238 | 0 | 0.039318191 | 0.011380302 | 0.149933045 | 0 | 0.047206021 | 0 | 0 | 0.026660374 | 0.020250453 | 0.006218354 | 0 | 0.022671621 | 0.007077131 | 0 | 0 | 0.137196248 |
| GPS20 | -0.031367711 | 0 | 0 | 0 | 0 | 0 | 0.116895063 | 0.018994423 | 0.092376214 | 0 | 0.21798092 | 0 | 0.019666335 | 0.047129074 | 0.192811109 | 0 | 0 | 0.191642921 | 0.047206021 | 0 | 0 | 0 | 0 | 0 | 0 | 0 | 0.066575939 | 0 | -0.014988064 | 0 | 0 |
| BISn1 | 0 | 0 | 0 | 0 | 0 | 0 | 0 | 0 | 0 | 0 | 0 | 0 | 0 | 0.041034318 | 0 | -0.061834116 | 0 | 0.06656256 | 0 | 0 | 0 | 0.260456865 | 0.025192349 | 0 | 0.091322886 | 0.209948267 | 0 | 0 | 0 | 0 | 0 |
| BIS2 | 0 | 0.001070892 | 0 | 0 | 0 | 0.005597996 | 0 | 0 | -0.012562609 | 0 | 0.001918385 | 0 | 0.043993146 | 0 | 0 | 0 | 0 | 0 | 0 | 0 | 0.260456865 | 0 | 0.182299946 | 0 | 0.060092145 | 0.343533186 | 0 | 0.041078554 | 0.147169988 | 0 | 0.036095479 |
| BIS3 | 0 | 0.017049228 | 0 | 0 | 0.011196092 | 0 | 0 | 0.065853196 | 0.015335534 | 0 | 0.07738989 | 0 | 0 | 0 | 0 | 0 | 0 | 0 | 0.026660374 | 0 | 0.025192349 | 0.182299946 | 0 | 0.127229147 | 0.138313227 | 0.117609647 | 0 | 0 | 0 | 0 | 0 |
| BIS4 | 0 | 0 | 0 | 0 | 0.101699554 | 0 | 0.043035594 | 0 | 0 | 0 | 0 | 0 | 0.071112967 | 0 | 0 | 0 | 0 | 0 | 0.020250453 | 0 | 0 | 0 | 0.127229147 | 0 | 0.030629235 | 0.192331672 | 0 | 0 | 0 | 0 | 0 |
| BIS5 | -0.011849901 | 0 | 0 | 0 | 0 | 0.008691288 | 0 | 0 | 0 | 0 | 0 | 0.093727917 | 0 | 0 | 0 | 0 | 0 | 0.000583947 | 0.006218354 | 0 | 0.091322886 | 0.060092145 | 0.138313227 | 0.030629235 | 0 | 0.233475454 | 0.13085609 | 0.187862414 | 0 | 0 | 0 |
| BIS6 | -0.058188396 | 0 | 0 | 0 | 0.006130292 | 0 | 0 | 0 | 0 | 0 | 0 | 0 | 0 | 0.004818289 | 0 | 0 | 0 | 0 | 0 | 0 | 0.209948267 | 0.343533186 | 0.117609647 | 0.192331672 | 0.233475454 | 0 | 0 | 0 | 0 | 0 | 0 |
| BIS7 | -0.022159898 | 0 | 0 | 0 | -0.027533389 | 0 | 0 | 0 | 0.090348678 | 0 | 0 | 0 | 0 | 0 | 0.003951918 | 0 | 0 | 0 | 0.022671621 | 0.066575939 | 0 | 0 | 0 | 0 | 0.13085609 | 0 | 0 | 0 | 0.167418951 | 0.06081486 | 0.056869186 |

| | | | | | | | | | | | | | | | | | | | | | | | | | | | | | | | |
|---|---|---|---|---|---|---|---|---|---|---|---|---|---|---|---|---|---|---|---|---|---|---|---|---|---|---|---|---|---|---|---|
| **BIS 8** | 0 | 0 | -0.040166087 | 0 | -0.068423947 | 0.073101979 | -0.000118229 | 0 | 0 | 0 | 0 | 0 | 0 | 0 | 0 | 0 | 0 | 0 | 0.007077131 | 0 | 0 | 0.041078554 | 0 | 0 | 0.187862414 | 0 | 0 | 0 | 0 | 0 | 0.437513813 |
| **BIS 9** | 0 | 0.038320801 | 0.067805827 | 0 | 0 | 0 | 0 | 0.006987691 | 0 | 0 | 0.003061468 | 0 | 0 | 0 | 0 | 0.082955495 | 0.051888832 | 0 | 0 | -0.014988064 | 0 | 0.147169988 | 0 | 0 | 0 | 0 | 0.167418951 | 0 | 0 | 0 | 0 |
| **BIS 10** | 0 | 0 | 0 | 0 | 0 | -0.035997055 | 0 | 0 | 0 | 0 | 0 | 0 | 0 | 0 | 0 | 0 | 0 | -0.033632178 | 0 | 0 | 0 | 0 | 0 | 0 | 0 | 0 | 0.06081486 | 0 | 0 | 0 | 0 |
| **BIS 11** | -0.03219404 | -0.077451364 | 0 | 0 | 0 | 0 | 0 | 0.105892895 | 0 | 0 | 0 | 0 | 0 | 0 | 0.016422345 | 0 | 0 | 0 | 0.137196248 | 0 | 0 | 0.036095479 | 0 | 0 | 0 | 0 | 0.056869186 | 0.437513813 | 0 | 0 | 0 |

**S2 Table S3:** Component Loadings from Principal Component Analysis of GPS and BIS No-Planning Items After Removing High Bridge Centrality Items

| | Component 1 | Component 2 | Uniqueness |
|---|---|---|---|
| **GPS1** | | -0.304 | 0.912 |
| **GPS2** | 0.51 | | 0.741 |
| **GPS3** | 0.342 | | 0.859 |
| **GPS4** | 0.38 | | 0.843 |
| **GPS5** | | | 0.918 |
| **GPS6** | | | 0.903 |
| **GPS7** | 0.49 | | 0.767 |
| **GPS8** | | | 0.886 |
| **GPS9** | 0.674 | | 0.564 |
| **GPS10** | 0.8 | | 0.38 |
| **GPS11** | 0.539 | | 0.633 |
| **GPS12** | 0.603 | | 0.636 |
| **GPS13** | 0.635 | | 0.58 |
| **GPS14** | 0.765 | | 0.399 |
| **GPS15** | 0.729 | | 0.471 |
| **GPS16** | 0.635 | | 0.608 |
| **GPS17** | 0.59 | | 0.66 |
| **GPS18** | 0.496 | | 0.674 |
| **GPS20** | 0.669 | | 0.55 |
| **BISnoplan1** | | 0.718 | 0.499 |
| **BISnoplan2** | | 0.8 | 0.359 |
| **BISnoplan3** | | 0.621 | 0.574 |
| **BISnoplan4** | | 0.477 | 0.742 |
| **BISnoplan5** | | 0.741 | 0.464 |
| **BISnoplan6** | | 0.801 | 0.353 |
| **BISnoplan7** | | | 0.876 |
| **BISnoplan8** | | 0.413 | 0.836 |
| **BISnoplan10** | | | 0.994 |

**S3 The Genetic Correlation Between Impulsivity and procrastination**

We first conducted model comparisons for the univariate and bivariate heritability models of procrastination and impulsivity. The comparison results are presented in S3 **Table S1**.

The genetic correlation between impulsivity and procrastination established in existing published studies, with the results of this study are detailed in S3 **Table S2**: Single Meta-Analytic Indicators of Heritability Estimates for Procrastination and Its Genetic Correlation with Impulsivity.

**S3 Table S1:** Model comparison results for univariate and bivariate heritability estimates of procrastination and non-planning impulsivity.

| Model Name | df | AIC | BIC |
|---|---|---|---|
| Procrastination | | | |
| ACE | 150 | 113.42 | -238.15 |
| **AE** | **151** | **111.42** | **-242.50** |
| CE | 151 | 119.89 | -234.03 |
| E | 152 | 132.03 | -224.23 |
| Non-planning impulsivity | | | |
| ACE | 150 | 130.08 | -221.49 |
| **AE** | **151** | **128.21** | **-225.70** |
| CE | 151 | 128.39 | -225.53 |
| E | 152 | 132.03 | -224.23 |
| Procrastination & Non-planning impulsivity | | | |
| ACE | 297 | 235.54 | -460.57 |
| **AE** | **300** | **229.62** | **-473.52** |
| CE | 300 | 238.41 | -464.73 |
| E | 3 | 251.71 | -458.46 |

**S3 Table S2:** Single Meta-Analytic Indicators of Heritability Estimates for Procrastination and Its Genetic Correlation with Impulsivity.

| Study | Author, year | MZ | DZ | age (Mean ±SD) | Gender (female/male) | $h^2$ | $r_g$ |
|---|---|---|---|---|---|---|---|
| *rg between procrastination and impulsivity: 3656 twin pairs (MZ= 2230 pairs, DZ= 1426 pairs) in total* | | | | | | | |
| 1 | Gustavson et al, 2014 | 181 pairs | 166 pairs | 22.66±1.12 | 364/299 | 46% | 1.0 |
| 2<br>3 | Loehlin & Martin, 2014 | Adult: 1310 pairs<br>Young: 702 pairs | Adult: 748 pairs<br>Young: 472 pairs | 42.5±13.0<br>23.2±2.19 | 2882/1234<br>1524/824 | 21% | 0.3 |
| 4 | Longitudinal Twin Adolescent Cohort | 37 pairs | 40 pairs | 19.61±2.43 | 86/68 | 63% | 0.51 |
| *$h^2$ of procrastination:* | | | | | | | |
| 1 | Gustavson et al, 2014 | 181 pairs | 166 pairs | 22.66±1.12 | 364/299 | 46% | 1.0 |
| 2 | Loehlin & Martin, 2014 | Adult: 1310 pairs<br>Young: 702 pairs | Adult: 748 pairs<br>Young: 472 pairs | 42.5±13.0<br>23.2±2.19 | 2882/1234<br>1524/824 | 21% | 0.3 |
| 3 | Longitudinal Twin Adolescent Cohort | 37 pairs | 40 pairs | 19.61±2.43 | 86/68 | 63% | 0.51 |
| 4 | Steel, P., 2007 | 118 pairs | 93 pairs | | | 22% | - |
| 5 | Gustavson et al., 2015 | 206 pairs | 179 pairs | | | 28% | - |
| 6 | Cesarini et al., 2012 | 1150 pairs | 2362 pairs | | | 18% | - |

### S4 Multivariate heritability analysis

To explore the role of individual morphological deviations in the genetic relationship between procrastination and impulsivity, we applied multivariate modeling to these three traits. Using the ACE framework, we decomposed the variance of these traits and evaluated their shared genetic and environmental contributions. In an iterative process, we compared the full trivariate model (Cholesky decomposition) with more restrictive models, including the Independent Pathway model and the Common Pathway model. These models incorporate both specific and shared variance but differ in how they distribute shared variance across the traits.

The Cholesky trivariate model decomposes the covariation between individual morphological deviation, impulsivity and procrastination into additive genetic (A), shared environmental (C), and non-shared environmental (E) factors, layering these factors sequentially into the model. This approach allows each factor to exert influence across an increasing sequence of traits. For example, genetic and environmental influences on the first trait may also affect the other traits, whereas the second trait’s influences affect only itself and subsequent traits. Based on our hypothesis that procrastination is a genetic byproduct of impulsivity, we ordered the traits in the Cholesky decomposition as individual morphological deviation, impulsivity and procrastination. Thus, the first set of latent ACE factors includes genetic, shared, and non-shared environmental contributions to individual morphological deviation, which may also influence impulsivity and procrastination. The second set of latent ACE factors applies to impulsivity independently of individual morphological deviation but may extend to procrastination. The third set of ACE factors represents unique influences on procrastination. It’s important to note that while the trait order affects interpretive clarity, it does not impact the model’s overall fit.

The Independent Pathway Model assumes that a single set of A, C, and E factors explains the shared variance among all traits, but the influence paths (or weights) of these factors vary for each trait. This model allows each trait to receive distinct variance contributions from the shared A, C, and E factors through different pathways. For example, the covariance between impulsivity and individual morphological deviation might be predominantly explained by A, whereas the covariance between impulsivity and procrastination might be more strongly attributed to E, and so on.

The Common Factor Model assumes that all traits are connected through a single common latent factor, characterized by a single set of A, C, and E variance components. However, each trait retains its unique genetic and environmental components. This model is advantageous for cases where the traits share a high degree of commonality, as it simplifies the covariance structure by assuming that the relationship between traits is entirely mediated by a single latent factor. For instance, the relationship among individual morphological deviation, impulsivity and procrastination may be primarily driven by a common factor, while each trait's unique components still influence its variance independently without affecting the covariance among the traits.

For each model type, we also compared ACE, AE, CE, and E models. We used -2 log-likelihood ratio tests for nested models and AIC as additional criteria, and fitted the multivariate models using the CSOLNP optimizer. To estimate the 95% confidence intervals (CIs) for model parameters, we employed likelihood-based confidence interval estimation.

Schematic diagrams of the three multivariate heritability models are presented in **Figure S5**.

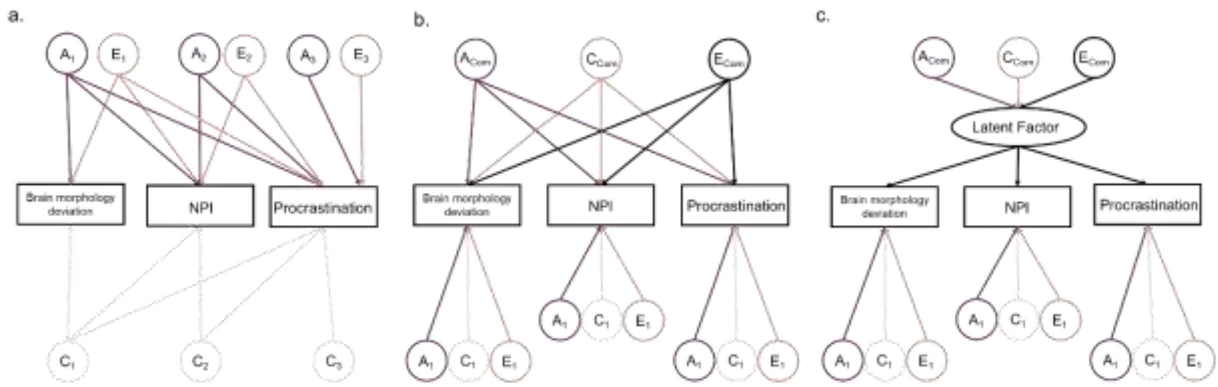

**Figure S5** Multivariate heritability model structures for brain morphology deviations, procrastination and impulsivity.

**S4 Table S1**: Model comparison results of multivariate ACE twin analysis for individual morphological deviation of global measures, impulsivity and procrastination.

| Name | No.parameters | -2 log-likelihood | df | AIC | No.parameters | -2 log-likelihood | df | AIC |
|---|---|---|---|---|---|---|---|---|
| | **global average deviation scores (global ADS)** | | | | **average deviation scores of cortical thickness (CT ADS)** | | | |
| Cholesky ACE | 21 | 1130.16 | 404 | 322.16 | 21 | 1140.59 | 405 | 330.59 |
| Cholesky AE | 15 | **1131.55** | 410 | **311.55** | 15 | **1147.76** | 411 | **325.76** |
| Cholesky CE | 15 | 1141.8 | 410 | 321.8 | 15 | 1149.08 | 411 | 327.08 |
| Cholesky E | 9 | 1180.92 | 416 | 348.92 | 9 | 1189.5 | 417 | 355.5 |
| IP ACE | 21 | 1130.17 | 404 | 322.17 | 21 | 1140.62 | 405 | 330.62 |
| IP AE | 15 | 1131.55 | 410 | 311.55 | 15 | 1148.73 | 411 | 326.73 |
| IP CE | 15 | 1141.8 | 410 | 321.8 | 15 | 1151.28 | 411 | 329.28 |
| IP E | 9 | 1181.33 | 416 | 349.33 | 9 | 1191.88 | 417 | 357.88 |
| CP ACE | 18 | 1134.68 | 408 | 318.68 | 18 | 1144.3 | 409 | 326.3 |
| CP AE | 14 | 1135.42 | 412 | 311.42 | 14 | 1150.87 | 413 | 324.87 |
| CP CE | 14 | 1143.37 | 412 | 319.37 | 14 | 1151.83 | 413 | 325.83 |
| CP E | 10 | 1181.33 | 416 | 349.33 | 10 | 1191.88 | 417 | 357.88 |
| | **average deviation scores of surface area (SA ADS)** | | | | | | | |
| Cholesky ACE | 21 | 1150.05 | 405 | 340.05 | | | | |
| Cholesky AE | 15 | **1150.43** | 411 | **328.43** | | | | |
| Cholesky CE | 15 | 1159.13 | 411 | 337.13 | | | | |
| Cholesky E | 9 | 1189.91 | 417 | 355.91 | | | | |
| IP ACE | 21 | 1150.43 | 405 | 340.43 | | | | |
| IP AE | 15 | 1150.83 | 411 | 328.83 | | | | |
| IP CE | 15 | 1160.83 | 411 | 338.83 | | | | |

| | | | | |
|---|---|---|---|---|
| IP E | 9 | 1189.91 | 417 | 355.91 |
| CP ACE | 18 | 1152.42 | 409 | 334.42 |
| CP AE | 14 | 1152.42 | 413 | 326.42 |
| CP CE | 14 | 1163.95 | 413 | 337.95 |
| CP E | 10 | 1189.91 | 417 | 355.91 |

**S4 Table S2**: Model comparison results of multivariate ACE twin analysis for individual morphological deviation of left rostral middle frontal, impulsivity and procrastination.

| Name | No.parameters | -2 log-likelihood | df | AIC | No.parameters | -2 log-likelihood | df | AIC |
|---|---|---|---|---|---|---|---|---|
| | The cortical thickness of left rostralmiddlefrontal | | | | The surface average of left rostralmiddlefrontal | | | |
| Cholesky ACE | 21 | 1154.16 | 405 | 344.16 | 21 | 1157.9 | 404 | 349.9 |
| Cholesky AE | 15 | **1154.75** | 411 | **332.75** | 15 | **1157.95** | 410 | **337.95** |
| Cholesky CE | 15 | 1163.18 | 411 | 341.18 | 15 | 1166.5 | 410 | 346.5 |
| Cholesky E | 9 | 1192.57 | 417 | 358.57 | 9 | 1183.26 | 416 | 351.26 |
| IP ACE | 21 | 1154.56 | 405 | 344.56 | 21 | 1157.91 | 404 | 349.91 |
| IP AE | 15 | 1155.26 | 411 | 333.26 | 15 | 1157.95 | 410 | 337.95 |
| IP CE | 15 | 1163.23 | 411 | 341.23 | 15 | 1166.51 | 410 | 346.51 |
| IP E | 9 | 1192.58 | 417 | 358.58 | 9 | 1183.27 | 416 | 351.27 |
| CP ACE | 18 | 1155.43 | 409 | 337.43 | 18 | 1158.6 | 408 | 342.6 |
| CP ACE | 18 | 1155.43 | 409 | 337.43 | 18 | 1158.6 | 408 | 342.6 |
| CP AE | 14 | 1156.01 | 413 | 330.01 | 14 | 1158.6 | 412 | 334.6 |
| CP CE | 14 | 1163.86 | 413 | 337.86 | 14 | 1166.56 | 412 | 342.56 |
| CP E | 10 | 1192.58 | 417 | 358.58 | 10 | 1183.27 | 416 | 351.27 |

**S4 Table S3**: Model comparison results of multivariate ACE twin analysis for individual morphological deviation of left medial orbitofrontal, impulsivity and procrastination.

| Name | No.parameters | -2 log-likelihood | df | AIC | No.parameters | -2 log-likelihood | df | AIC |
|---|---|---|---|---|---|---|---|---|
| | The cortical thickness of left medialorbitofrontal | | | | The surface average of left medialorbitofrontal | | | |
| Cholesky ACE | 21 | 1151.41 | 404 | 343.41 | 21 | 1164.27 | 405 | 354.27 |
| Cholesky AE | 15 | **1151.41** | 410 | **331.41** | 15 | **1167** | 411 | **345** |
| Cholesky CE | 15 | 1160.24 | 410 | 340.24 | 15 | 1171.25 | 411 | 349.25 |
| Cholesky E | 9 | 1178.7 | 416 | 346.7 | 9 | 1194.34 | 417 | 360.34 |
| IP ACE | 21 | 1151.84 | 404 | 343.84 | 21 | 1164.57 | 405 | 354.57 |
| IP AE | 15 | 1152.34 | 410 | 332.34 | 15 | 1167.67 | 411 | 345.67 |
| IP CE | 15 | 1160.93 | 410 | 340.93 | 15 | 1173.82 | 411 | 351.82 |
| IP E | 9 | 1178.84 | 416 | 346.84 | 9 | 1194.34 | 417 | 360.34 |
| CP ACE | 18 | 1152.21 | 408 | 336.21 | 18 | 1167.05 | 409 | 349.05 |
| CP ACE | 18 | 1152.21 | 408 | 336.21 | 18 | 1167.05 | 409 | 349.05 |
| CP AE | 14 | 1152.21 | 412 | 328.21 | 14 | 1168.02 | 413 | 342.02 |
| CP CE | 14 | 1160.96 | 412 | 336.96 | 14 | 1174.97 | 413 | 348.97 |
| CP E | 10 | 1178.84 | 416 | 346.84 | 10 | 1194.34 | 417 | 360.34 |

**S4 Table S4:** Cholesky model parameters calculated using multivariate ACE twin modelling of global morphological measures.

| | Additive Genetic | | | Shared Environment | | | Non-shared Environmental | | |
|---|---|---|---|---|---|---|---|---|---|
| **Original** | **1** | **2** | **3** | **1** | **2** | **3** | **1** | **2** | **3** |
| | | | | **global average deviation scores (global ADS)** | | | | | |
| **AE Path Coefficients** | | | | | | | | | |
| global ADS | 0.74 (0.56; 0.92) | - | - | 0 | - | - | 0.62 (0.5; 0.77) | - | - |
| BIS-noplanning | 0.13 (-0.15; 0.45) | 0.58 (-0.84; 0.84) | - | 0 | 0 | - | -0.28 (-0.53; -0.04) | 0.76 (0.61; 0.95) | - |
| GPS | 0.16 (-0.11; 0.43) | 0.27 (-0.85; 0.85) | 0.73 (-0.92; 0.92) | 0 | 0 | 0 | -0.11 (-0.31; 0.08) | 0.11 (-0.08; 0.32) | 0.6 (0.48; 0.75) |
| **AE (Co)variance components** | | | | | | | | | |
| global ADS | 0.59 (0.37; 0.74) | - | - | 0 | - | - | 0.41 (0.26; 0.63) | - | - |
| BIS-noplanning | NA | 0.35 (0.03; 0.6) | - | 0 | 0 | - | NA | 0.65 (0.4; 0.97) | - |
| GPS | NA | 0.6 (-0.28; 1.15) | 0.62 (0.39; 0.77) | 0 | 0 | 0 | NA | 0.4 (-0.15; 1.28) | 0.38 (0.23; 0.61) |
| | | | | **average deviation scores of cortical thickness (CT ADS)** | | | | | |
| **AE Path Coefficients** | | | | | | | | | |
| CT ADS | 0.73 (0.54; 0.91) | - | - | 0 | - | - | 0.67 (0.56; 0.83) | - | - |
| BIS-noplanning | -0.09 (-0.37; 0.2) | -0.59 (-0.84; -0.19) | - | 0 | 0 | - | -0.1 (-0.35; 0.14) | 0.8 (0.64; 1) | - |
| GPS | 0.14 (-0.13; 0.42) | -0.31 (-0.68; 0.08) | NA | 0 | 0 | 0 | -0.01 (-0.19; 0.18) | 0.14 (-0.04; 0.35) | 0.6 (0.49; 0.75) |
| **AE (Co)variance components** | | | | | | | | | |
| CT ADS | 0.54 (0.32; 0.69) | - | - | 0 | - | - | 0.46 (0.31; 0.68) | - | - |
| BIS-noplanning | NA | 0.36 (0.04; 0.6) | - | 0 | 0 | - | NA | 0.64 (0.4; 0.96) | - |
| GPS | NA | 0.6 (-0.3; 1.14) | 0.62 (0.39; 0.77) | 0 | 0 | 0 | NA | 0.4 (-0.14; 1.3) | 0.38 (0.23; 0.61) |
| | | | | **average deviation scores of surface area (SA ADS)** | | | | | |
| **AE Path Coefficients** | | | | | | | | | |
| SA ADS | 0.69 (0.46; 0.89) | - | - | 0 | NA | NA | 0.71 (0.58; 0.88) | - | - |
| BIS-noplanning | -0.06 (-0.37; 0.28) | 0.6 (0.18; 0.84) | - | 0 | 0 | NA | -0.11 (-0.37; 0.13) | 0.79 (0.64; 0.99) | - |

| | | | | | | | | | |
|---|---|---|---|---|---|---|---|---|---|
| GPS | -0.26 (-0.58; 0.03) | 0.28 (-0.11; 0.79) | 0.67 (-0.88; 0.88) | 0 | 0 | 0 | 0.04 (-0.16; 0.24) | 0.14 (-0.05; 0.35) | 0.61 (0.49; 0.76) |
| **AE (Co)variance components** | | | | | | | | | |
| SA ADS | 0.49 (0.23; 0.67) | - | - | 0 | - | - | 0.51 (0.33; 0.77) | - | - |
| BIS-noplanning | NA | 0.36 (0.04; 0.61) | - | 0 | 0 | - | NA | 0.64 (0.39; 0.96) | - |
| GPS | NA | 0.64 (-0.21; 1.21) | 0.61 (0.37; 0.76) | 0 | 0 | 0 | NA | 0.36 (-0.21; 1.21) | 0.39 (0.24; 0.63) |

*1, 2, 3 = first, second and third latent ACE factors.

NA = alpha level not reached; confidence interval could not be calculated.

**S4 Table S5:** Cholesky model parameters calculated using multivariate ACE twin modelling of global morphological measures.

| | Additive Genetic | | | Shared Environment | | | Non-shared Environmental | | |
|---|---|---|---|---|---|---|---|---|---|
| **Original** | **1** | **2** | **3** | **1** | **2** | **3** | **1** | **2** | **3** |
| | | | | **global average deviation scores (global ADS)** | | | | | |
| **AE Path Coefficients** | | | | | | | | | |
| global ADS | 0.74 (0.56; 0.92) | - | - | 0 | - | - | 0.62 (0.5; 0.77) | - | - |
| BIS-noplanning | 0.13 (-0.15; 0.45) | 0.58 (-0.84; 0.84) | - | 0 | 0 | - | -0.28 (-0.53; -0.04) | 0.76 (0.61; 0.95) | - |
| GPS | 0.16 (-0.11; 0.43) | 0.27 (-0.85; 0.85) | 0.73 (-0.92; 0.92) | 0 | 0 | 0 | -0.11 (-0.31; 0.08) | 0.11 (-0.08; 0.32) | 0.6 (0.48; 0.75) |
| **AE (Co)variance components** | | | | | | | | | |
| global ADS | 0.59 (0.37; 0.74) | - | - | 0 | - | - | 0.41 (0.26; 0.63) | - | - |
| BIS-noplanning | NA | 0.35 (0.03; 0.6) | - | 0 | 0 | - | NA | 0.65 (0.4; 0.97) | - |
| GPS | NA | 0.6 (-0.28; 1.15) | 0.62 (0.39; 0.77) | 0 | 0 | 0 | NA | 0.4 (-0.15; 1.28) | 0.38 (0.23; 0.61) |
| | | | | **average deviation scores of cortical thickness (CT ADS)** | | | | | |
| **AE Path Coefficients** | | | | | | | | | |
| CT ADS | 0.73 (0.54; 0.91) | - | - | 0 | - | - | 0.67 (0.56; 0.83) | - | - |
| BIS-noplanning | -0.09 (-0.37; 0.2) | -0.59 (-0.84; -0.19) | - | 0 | 0 | - | -0.1 (-0.35; 0.14) | 0.8 (0.64; 1) | - |
| GPS | 0.14 (-0.13; 0.42) | -0.31 (-0.68; 0.08) | NA | 0 | 0 | 0 | -0.01 (-0.19; 0.18) | 0.14 (-0.04; 0.35) | 0.6 (0.49; 0.75) |
| **AE (Co)variance components** | | | | | | | | | |
| CT ADS | 0.54 (0.32; 0.69) | - | - | 0 | - | - | 0.46 (0.31; 0.68) | - | - |
| BIS-noplanning | NA | 0.36 (0.04; 0.6) | - | 0 | 0 | - | NA | 0.64 (0.4; 0.96) | - |
| GPS | NA | 0.6 (-0.3; 1.14) | 0.62 (0.39; 0.77) | 0 | 0 | 0 | NA | 0.4 (-0.14; 1.3) | 0.38 (0.23; 0.61) |
| | | | | **average deviation scores of surface area (SA ADS)** | | | | | |
| **AE Path Coefficients** | | | | | | | | | |
| SA ADS | 0.69 (0.46; 0.89) | - | - | 0 | NA | NA | 0.71 (0.58; 0.88) | - | - |
| BIS-noplanning | -0.06 (-0.37; 0.28) | 0.6 (0.18; 0.84) | - | 0 | 0 | NA | -0.11 (-0.37; 0.13) | 0.79 (0.64; 0.99) | - |

| | | | | | | | | | |
|---|---|---|---|---|---|---|---|---|---|
| GPS | -0.26 (-0.58; 0.03) | 0.28 (-0.11; 0.79) | 0.67 (-0.88; 0.88) | 0 | 0 | 0 | 0.04 (-0.16; 0.24) | 0.14 (-0.05; 0.35) | 0.61 (0.49; 0.76) |
| **AE (Co)variance components** | | | | | | | | | |
| SA ADS | 0.49 (0.23; 0.67) | - | - | 0 | - | - | 0.51 (0.33; 0.77) | - | - |
| BIS-noplanning | NA | 0.36 (0.04; 0.61) | - | 0 | 0 | - | NA | 0.64 (0.39; 0.96) | - |
| GPS | NA | 0.64 (-0.21; 1.21) | 0.61 (0.37; 0.76) | 0 | 0 | 0 | NA | 0.36 (-0.21; 1.21) | 0.39 (0.24; 0.63) |

*1, 2, 3 = first, second and third latent ACE factors.

NA = alpha level not reached; confidence interval could not be calculated.

**S4 Table S6:** Cholesky model parameters calculated using multivariate ACE twin modelling of global morphological measures.

| | Additive Genetic | | | Shared Environment | | | Non-shared Environmental | | |
|---|---|---|---|---|---|---|---|---|---|
| **Original** | **1** | **2** | **3** | **1** | **2** | **3** | **1** | **2** | **3** |
| | | | | **global average deviation scores (global ADS)** | | | | | |
| **AE Path Coefficients** | | | | | | | | | |
| BIS-noplanning | 0.6 (0.18; 0.85) | - | - | 0 | - | - | 0.81 (0.65; 1.01) | - | - |
| GPS | 0.29 (-0.11; 0.69) | NA | - | 0 | 0 | - | 0.14 (-0.05; 0.35) | 0.6 (0.49; 0.75) | - |
| global ADS | 0.16 (-0.18; 0.78) | NA | 0.72 (-0.91; 0.91) | 0 | 0 | 0 | -0.21 (-0.41; -0.03) | -0.07 (-0.25; 0.12) | 0.57 (0.47; 0.72) |
| **AE (Co)variance components** | | | | | | | | | |
| BIS-noplanning | 0.35 (0.03; 0.6) | - | - | 0 | - | - | 0.65 (0.4; 0.97) | - | - |
| GPS | 0.6 (-0.28; 1.15) | 0.62 (0.39; 0.77) | - | 0 | 0 | - | 0.4 (-0.15; 1.28) | 0.38 (0.23; 0.61) | - |
| global ADS | NA | NA | 0.59 (0.37; 0.74) | 0 | 0 | 0 | NA | NA | 0.41 (0.26; 0.63) |
| | | | | **average deviation scores of cortical thickness (CT ADS)** | | | | | |
| **AE Path Coefficients** | | | | | | | | | |
| BIS-noplanning | 0.6 (0.2; 0.85) | - | - | 0 | - | - | 0.8 (0.65; 1) | - | - |
| GPS | 0.29 (-0.11; 0.66) | 0.74 (0.48; 0.93) | - | 0 | 0 | - | 0.14 (-0.04; 0.35) | 0.6 (0.49; 0.75) | - |
| CT ADS | -0.11 (-0.49; 0.27) | 0.18 (-0.07; 0.46) | 0.69 (0.45; 0.88) | 0 | 0 | 0 | -0.08 (-0.29; 0.12) | 0.01 (-0.19; 0.21) | 0.67 (0.56; 0.82) |
| **AE (Co)variance components** | | | | | | | | | |
| BIS-noplanning | 0.36 (0.04; 0.6) | - | - | 0 | - | - | 0.64 (0.4; 0.96) | - | - |
| GPS | 0.6 (-0.3; 1.14) | 0.62 (0.39; 0.77) | - | 0 | 0 | - | 0.4 (-0.14; 1.3) | 0.38 (0.23; 0.61) | - |
| CT ADS | NA | NA | 0.54 (0.32; 0.69) | 0 | 0 | 0 | NA | NA | 0.46 (0.31; 0.68) |
| | | | | **average deviation scores of surface area (SA ADS)** | | | | | |
| **AE Path Coefficients** | | | | | | | | | |
| BIS-noplanning | 0.6 (0.19; 0.85) | - | - | 0 | - | - | 0.8 (0.65; 1.01) | - | - |
| GPS | 0.31 (-0.08; 0.71) | NA | - | 0 | 0 | - | 0.13 (-0.06; 0.34) | 0.61 (0.49; 0.77) | NA |

| | | | | | | | | | |
|---|---|---|---|---|---|---|---|---|---|
| SA ADS | -0.07 (-0.42; 0.42) | -0.22 (-0.69; 0.05) | 0.65 (-0.86; 0.86) | 0 | 0 | 0 | -0.1 (-0.32; 0.12) | 0.07 (-0.15; 0.28) | 0.7 (0.57; 0.86) |
| **AE (Co)variance components** | | | | | | | | | |
| BIS-noplanning | 0.36 (0.04; 0.61) | - | - | 0 | - | - | 0.64 (0.39; 0.96) | - | - |
| GPS | 0.64 (-0.21; 1.21) | 0.61 (0.37; 0.76) | - | 0 | 0 | - | 0.36 (-0.21; 1.21) | 0.39 (0.24; 0.63) | - |
| SA ADS | NA | NA | 0.49 (0.23; 0.67) | 0 | 0 | 0 | NA | NA | 0.51 (0.33; 0.77) |

*1, 2, 3 = first, second and third latent ACE factors.

NA = alpha level not reached; confidence interval could not be calculated.

**S4 Table S7**: Cholesky model parameters calculated using multivariate ACE twin modelling of regional morphological measures.

| | Additive Genetic | | | Shared Environment | | | Non-shared Environmental | | |
|---|---|---|---|---|---|---|---|---|---|
| **Original** | **1** | **2** | **3** | **1** | **2** | **3** | **1** | **2** | **3** |
| | | | **the thickness average of left rostral middle frontal** | | | | | | |
| **AE Path Coefficients** | | | | | | | | | |
| Lrostral middle frontal thickavg | 0.7 (0.48; 0.9) | - | - | 0 | - | - | 0.71 (0.58; 0.88) | - | - |
| BIS-noplanning | -0.04 (-0.35; 0.27) | 0.6 (0.21; 0.85) | - | 0 | 0 | - | 0.07 (-0.18; 0.31) | 0.8 (0.64; 1) | - |
| GPS | 0.19 (-0.1; 0.49) | 0.29 (-0.11; 0.67) | 0.71 (0.4; 0.9) | 0 | 0 | 0 | -0.02 (-0.21; 0.17) | 0.15 (-0.04; 0.36) | 0.6 (0.49; 0.75) |
| **AE (Co)variance components** | | | | | | | | | |
| Lrostral middle frontal thickavg | 0.49 (0.25; 0.67) | - | - | 0 | - | - | 0.51 (0.33; 0.75) | - | - |
| BIS-noplanning | NA | 0.37 (0.05; 0.61) | - | 0 | 0 | - | NA | 0.63 (0.39; 0.95) | - |
| GPS | NA | NA | 0.62 (0.38; 0.77) | 0 | 0 | 0 | NA | NA | 0.38 (0.23; 0.62) |
| | | | **the surface average of left rostral middle frontal** | | | | | | |
| **AE Path Coefficients** | | | | | | | | | |
| Lrostral middle frontal surfavg | 0.22 (-0.6; 0.6) | - | - | 0 | - | - | 0.94 (0.78; 1.09) | - | - |
| BIS-noplanning | 0.11 (-0.84; 0.84) | 0.59 (-0.85; 0.85) | - | 0 | 0 | - | -0.1 (-0.33; 0.14) | 0.79 (0.64; 1) | - |
| GPS | 0.22 (-0.98; 0.98) | 0.26 (-0.98; 0.98) | 0.71 (-0.92; 0.92) | 0 | 0 | 0 | -0.07 (-0.25; 0.12) | 0.13 (-0.05; 0.34) | 0.6 (0.48; 0.75) |
| **AE (Co)variance components** | | | | | | | | | |
| Lrostral middle frontal surfavg | 0.05 (0; 0.34) | - | - | 0 | - | - | 0.95 (0.66; 1) | - | - |
| BIS-noplanning | NA | 0.36 (0.04; 0.6) | - | 0 | 0 | - | NA | 0.64 (0.4; 0.96) | - |
| GPS | NA | 0.61 (-0.27; 1.16) | 0.62 (0.39; 0.77) | 0 | 0 | 0 | NA | 0.39 (-0.16; 1.27) | 0.38 (0.23; 0.61) |
| | | | **the thickness average of left medial orbitofrontal** | | | | | | |
| **AE Path Coefficients** | | | | | | | | | |

| | | | | | | | | | |
|---|---|---|---|---|---|---|---|---|---|
| Lmedial orbitofrontal thickavg | 0.42 (-0.67; 0.67) | - | - | 0 | - | - | 0.86 (0.72; 1.03) | - | - |
| BIS-noplanning | -0.21 (-0.79; 0.79) | 0.57 (-0.84; 0.84) | - | 0 | 0 | - | 0.06 (-0.18; 0.28) | 0.8 (0.64; 1) | - |
| GPS | 0.04 (-0.89; 0.89) | 0.32 (-0.95; 0.95) | 0.72 (-0.92; 0.92) | 0 | 0 | 0 | 0 (-0.19; 0.18) | 0.14 (-0.04; 0.35) | 0.6 (0.49; 0.76) |
| **AE (Co)variance components** | | | | | | | | | |
| Lmedial orbitofrontal thickavg | 0.19 (0; 0.43) | - | - | 0 | - | - | 0.81 (0.57; 1) | - | - |
| BIS-noplanning | NA | 0.37 (0.05; 0.61) | - | 0 | 0 | - | NA | 0.63 (0.39; 0.95) | - |
| GPS | NA | 0.6 (-0.29; 1.15) | 0.62 (0.38; 0.77) | 0 | 0 | 0 | NA | 0.4 (-0.15; 1.29) | 0.38 (0.23; 0.62) |
| **The surface average of left medial orbitofrontal** | | | | | | | | | |
| **AE Path Coefficients** | | | | | | | | | |
| Lmedial orbitofrontal surfavg | -0.41 (-0.69; 0.69) | - | - | 0 | - | - | 0.91 (0.75; 1.09) | - | - |
| BIS-noplanning | -0.15 (-0.76; 0.76) | 0.57 (-0.84; 0.84) | - | 0 | 0 | - | -0.1 (-0.33; 0.14) | 0.8 (0.65; 1.01) | - |
| GPS | 0.13 (-0.91; 0.91) | 0.35 (-0.96; 0.96) | 0.69 (-0.91; 0.91) | 0 | 0 | 0 | 0.02 (-0.16; 0.22) | 0.14 (-0.05; 0.35) | 0.6 (0.49; 0.76) |
| **AE (Co)variance components** | | | | | | | | | |
| Lmedial orbitofrontal surfavg | 0.17 (0; 0.43) | - | - | 0 | - | - | 0.83 (0.57; 1) | - | - |
| BIS-noplanning | NA; | 0.34 (0.03; 0.59) | - | 0 | 0 | - | NA | 0.66 (0.41; 0.97) | - |
| GPS | NA | 0.62 (-0.24; 1.19) | 0.61 (0.38; 0.77) | 0 | 0 | 0 | NA | 0.38 (-0.19; 1.24) | 0.39 (0.23; 0.62) |

*1, 2, 3 = first, second and third latent ACE factors.
NA = alpha level not reached; confidence interval could not be calculated.

**S4 Table S8**: Cholesky model parameters calculated using multivariate ACE twin modelling of regional morphological measures.

| | Additive Genetic | | | Shared Environment | | | Non-shared Environmental | | |
|---|---|---|---|---|---|---|---|---|---|
| **Original** | **1** | **2** | **3** | **1** | **2** | **3** | **1** | **2** | **3** |
| | | | | **the thickness average of left rostral middle frontal** | | | | | |
| **AE Path Coefficients** | | | | | | | | | |
| BIS-noplanning | 0.61 (0.21; 0.86) | - | - | 0 | - | - | 0.8 (0.65; 1) | - | - |
| GPS | 0.28 (-0.12; 0.64) | 0.74 (0.5; 0.92) | - | 0 | 0 | - | 0.15 (-0.04; 0.36) | 0.6 (0.49; 0.75) | - |
| Lrostral middle frontal thickavg | -0.05 (-0.45; 0.34) | 0.2 (-0.07; 0.48) | 0.67 (0.37; 0.88) | 0 | 0 | 0 | 0.06 (-0.16; 0.28) | -0.04 (-0.26; 0.18) | 0.7 (0.58; 0.87) |
| **AE (Co)variance components** | | | | | | | | | |
| BIS-noplanning | 0.37 (0.05; 0.61) | - | - | 0 | - | - | 0.63 (0.39; 0.95) | - | - |
| GPS | NA | 0.62 (0.38; 0.77) | - | 0 | 0 | - | NA | 0.38 (0.23; 0.62) | - |
| Lrostral middle frontal thickavg | NA | NA | **0.49 (0.25; 0.67)** | 0 | 0 | 0 | NA | NA | **0.51 (0.33; 0.75)** |
| | | | | **the surface average of left rostral middle frontal** | | | | | |
| **AE Path Coefficients** | | | | | | | | | |
| BIS-noplanning | 0.6 (0.2; 0.85) | - | - | 0 | - | - | 0.8 (0.65; 1) | - | - |
| GPS | 0.29 (-0.1; 0.66) | 0.73 (0.47; 0.92) | - | 0 | 0 | - | 0.14 (-0.05; 0.35) | 0.6 (0.49; 0.75) | - |
| Lrostral middle frontal surfavg | 0.04 (-0.34; 0.42) | 0.05 (-0.24; 0.32) | -0.21 (-0.59; 0.59) | 0 | 0 | 0 | -0.12 (-0.38; 0.16) | -0.08 (-0.35; 0.21) | 0.93 (0.77; 1.08) |
| **AE (Co)variance components** | | | | | | | | | |
| BIS-noplanning | 0.36 (0.04; 0.6) | - | - | 0 | - | - | 0.64 (0.4; 0.96) | - | - |
| GPS | 0.61 (-0.27; 1.16) | 0.62 (0.39; 0.77) | - | 0 | 0 | - | 0.39 (-0.16; 1.27) | 0.38 (0.23; 0.61) | - |
| Lrostral middle frontal surfavg | NA | NA | 0.05 (0; 0.34) | 0 | 0 | 0 | NA | NA | **0.95 (0.66; 1)** |
| | | | | **the thickness average of left medial orbitofrontal** | | | | | |
| **AE Path Coefficients** | | | | | | | | | |

| | | | | | | | | | |
|---|---|---|---|---|---|---|---|---|---|
| BIS-noplanning | 0.61 (0.22; 0.86) | - | - | 0 | - | - | 0.8 (0.65; 1) | - | - |
| GPS | 0.29 (-0.11; 0.65) | 0.73 (0.49; 0.92) | - | 0 | 0 | - | 0.14 (-0.04; 0.35) | 0.6 (0.49; 0.76) | - |
| Lmedial orbitofrontal thickavg | -0.14 (-0.51; 0.21) | 0.08 (-0.17; 0.38) | 0.38 (-0.65; 0.65) | 0 | 0 | 0 | 0.06 (-0.19; 0.3) | -0.02 (-0.27; 0.23) | 0.85 (0.71; 1.03) |
| **AE (Co)variance components** | | | | | | | | | |
| BIS-noplanning | 0.37 (0.05; 0.61) | - | - | 0 | - | - | 0.63 (0.39; 0.95) | - | - |
| GPS | 0.6 (-0.29; 1.15) | 0.62 (0.38; 0.77) | - | 0 | 0 | - | 0.4 (-0.15; 1.29) | 0.38 (0.23; 0.62) | - |
| Lmedial orbitofrontal thickavg | NA | NA | 0.19 (0; 0.43) | 0 | 0 | 0 | NA | NA | **0.81 (0.57; 1)** |
| **The surface average of left medial orbitofrontal** | | | | | | | | | |
| **AE Path Coefficients** | | | | | | | | | |
| BIS-noplanning | 0.59 (0.17; 0.77) | - | - | 0 | - | - | 0.81 (0.64; 0.98) | - | - |
| GPS | 0.31 (-0.1; 0.71) | 0.72 (0.4; 0.84) | - | 0 | 0 | - | 0.14 (-0.06; 0.35) | 0.61 (0.47; 0.77) | - |
| Lmedial orbitofrontal surfavg | 0.11 (-0.27; 0.54) | -0.12 (-0.48; 0.16) | 0.38 (-0.65; 0.65) | 0 | 0 | 0 | -0.11 (-0.36; 0.16) | 0.06 (-0.2; 0.33) | 0.9 (0.75; 1) |
| **AE (Co)variance components** | | | | | | | | | |
| BIS-noplanning | 0.34 (0.03; 0.59) | - | - | 0 | - | - | 0.66 (0.41; 0.97) | - | - |
| GPS | 0.62 (-0.24; 1.19) | 0.61 (0.38; 0.77) | - | 0 | 0 | - | 0.38 (-0.19; 1.24) | 0.39 (0.23; 0.62) | - |
| Lmedial orbitofrontal surfavg | NA | NA | 0.17 (0; 0.43) | 0 | 0 | 0 | NA | NA | **0.83 (0.57; 1)** |

*1, 2, 3 = first, second and third latent ACE factors.
NA = alpha level not reached; confidence interval could not be calculated.

### S5 Generation of the Procrastination Mask

To investigate the neural correlates of procrastination, we conducted a mini meta-analysis on “Associations Between Procrastination Behavior and Brain Structure: A Morphological Meta-Analysis”. This analysis followed PRISMA guidelines and employed both forward and backward searching approaches to ensure comprehensive literature coverage (**Figure S6)**.

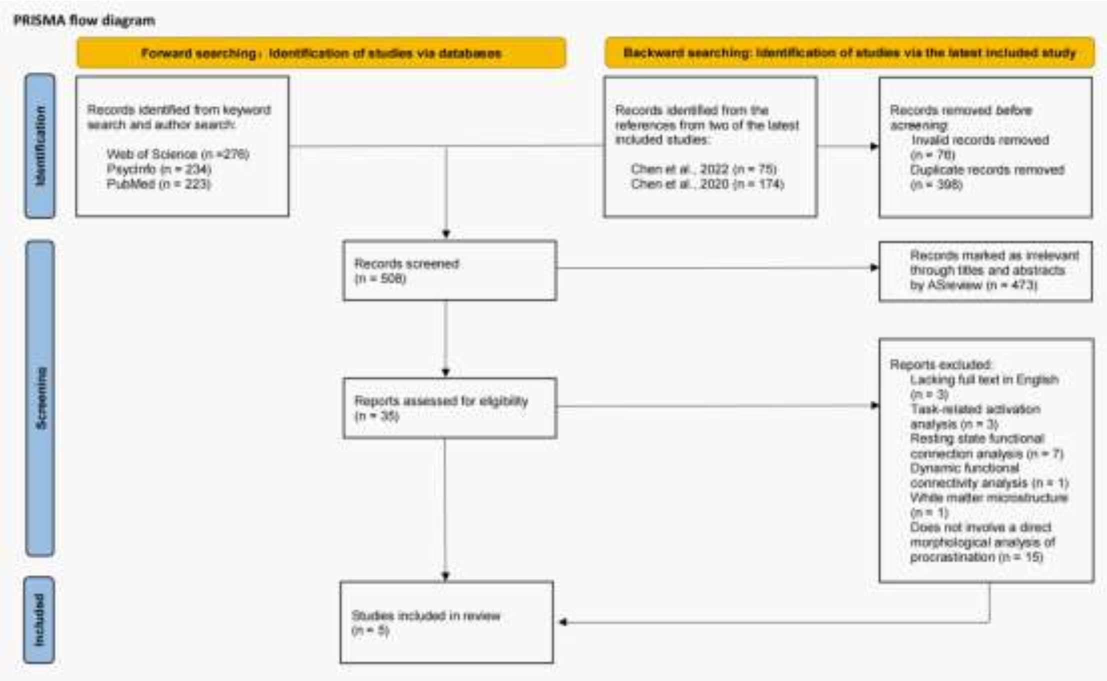

**Figure S6** PRISMA flow diagram for the mini meta-analysis on the brain structure of procrastination using voxel-based morphometry studies.

#### S5.1 Forward searching

For the forward searching, we systematically searched three major databases—Web of Science, PsycINFO, and PubMed—using a predefined set of keywords. The search was performed using the following keyword logic:

*procrastinat* AND (VBM OR voxel-based morphometry OR voxel-based analysis OR surface-based OR surface analysis OR surface morphometry OR surface mapping OR morpholog* OR structural MRI OR neuroimaging OR MRI OR magnetic resonance imaging OR brain imaging OR brain structure OR grey matter OR white matter OR cortical thickness OR brain volume OR surface OR cortical thickness OR diffusion tensor imaging OR DTI)*

Additionally, we searched all relevant publications by frequently cited authors in procrastination research, focusing on *Tingyong Feng*.

This forward search identified 132 articles from Web of Science, 98 from PsycINFO, and 93 from PubMed based on keywords, and 144, 136, and 130 articles respectively based on author-specific searches, yielding a total of 733 articles.

#### S5.2 Backward searching

Backward searching complemented this by reviewing the reference lists of the two most recent included studies:

*Chen, Z., Zhang, R., Xie, J., Liu, P., Zhang, C., Zhao, J., ... & Feng, T. (2022). Hybrid brain model accurately predict human procrastination behavior. Cognitive Neurodynamics, 16(5), 1107-1121.*

*Chen, Z., Liu, P., Zhang, C., & Feng, T. (2020). Brain morphological dynamics of procrastination: The crucial role of the self-control, emotional, and episodic prospection network. Cerebral Cortex, 30(5), 2834-2853.*

This step identified 75 and 174 references from these two studies, respectively, yielding a total of 249 references for further examination.

#### S5.3 Screening and inclusion process

After combining forward and backward searches in EndNote 20, a total of valid 906 articles were retrieved as of December 9, 2024, of which 398 duplicates were removed, leaving 508 unique articles for screening.

To refine the dataset, we employed ASReview (https://asreview.nl/), a machine learning tool for systematic reviews, which screened titles and abstracts to exclude irrelevant studies. This process eliminated 473 articles unrelated to the neural basis of procrastination. The remaining 35 articles underwent full-text review, and studies were excluded based on the following criteria: no full-text availability in English (3 articles), focus on task-related activation analysis (3 articles), focus on resting-state functional connectivity analysis (7 articles), focus on dynamic functional connectivity analysis (1 article), focus on white matter microstructures (1 article) and those examining procrastination as a dependent variable without direct morphological analysis (15 articles). After this rigorous screening, 5 studies were included for the meta-analysis.

#### S5.4 Seed-based d Mapping

We performed a voxel-based meta-analysis using Seed-based d Mapping (SDM) to identify procrastination-related brain regions. The MNI-based peak coordinates and corresponding t-statistics from the final 5 studies (S5 Table S1) served as inputs for the SDM analysis, generating the effect size map of procrastination.

Preprocessing was configured for grey matter analysis, employing a grey matter template and mask, an anisotropy of 1.0, a 20 mm isotropic full-width at half-maximum (FWHM) smoothing kernel, and a voxel size of 2 mm. Following preprocessing, meta-analytic mean calculations integrated effect sizes across studies. Family-wise error (FWE) correction was applied using 1000 subject-based permutations to generate a null distribution of maximum statistics, which was used to threshold the meta-analysis images, producing a corrected *p*-value map. A statistical threshold of $p < 0.05$ and an extent threshold of 10 voxels were applied to identify significant clusters. In the uncorrected *p*-value map, we observed two large clusters under both positive and negative SDM-Z thresholds, as detailed in S5 Table S2.

**S5 Table S1:** Peak Coordinates and Effect Sizes of Included Studies.

| Study | n | Age (years) | Gender (M/F) | Threshold | Brain Region | MNI | | | | t_thr |
|---|---|---|---|---|---|---|---|---|---|---|
| | | | | | | x | y | z | t | |
| (1) | n= 243 | 21.03± 2.05 | 112/131 | p < 0.001 without correction | Insula | -30 | 18 | -15 | 3.89 | 3.12 |
| | | | | | Anterior cingulate cortex | -6 | 42 | 2 | 3.66 | |
| | | | | | Parahippocampal gyrus | 24 | -6 | -38 | 3.87 | |
| | | | | | Middle frontal gyrus, dlPFC | -43 | 54 | 2 | -2.09 | |
| | | | | | Anterior cingulate cortex | -2 | 31 | 0 | 6.58 | |
| | | | | | Middle frontal gyrus, vmPFC | 4 | 41 | -13 | 2.98 | |
| | | | | | Orbital frontal cortex | -4 | 59 | -20 | 4.36 | |
| | | | | | Orbital frontal cortex | 6 | 42 | 2 | 4.01 | |
| | | | | | Orbital frontal cortex | -4 | 47 | -16 | 3.89 | |
| | | | | | Orbital frontal cortex | 9 | 51 | -2 | 3.73 | |
| (2) | n= 168 | 20.04± 1.83 | 42/126 | p < 0.05, non-stationary corrected | Hippocampus | 33 | -33 | -6 | 5.31 | 1.65 |
| | | | | | Precuneus | 11 | -72 | 20 | -4.99 | |
| | | | | p = 0.001 uncorrected | Superior temporal gyrus | -44 | -54 | -17 | -3.86 | 3.14 |
| (3) | n= 85 | 20.53± 2.07 | 30/55 | p < 0.05, corrected | Left middle frontal gyrus | -38 | 49 | 27 | -3.74 | 1.66 |
| | n= 84 | 19.51± 1.35 | 33/51 | | DLPFC | -33 | 36 | 37.5 | -5.73 | 1.66 |
| (4) | n=151 | 19.85± 1.60 | 99/52 | p < 0.05, corrected | Parahippocampal gyrus | 16 | -16 | -26 | 3.49 | 1.66 |
| | | | | | Orbital frontal cortex | -20 | 60 | -18 | 3.48 | |
| | | | | | Medial frontal gyrus | -16 | 70 | -4 | 3.71 | |
| | | | | | Lingual_L (aal)* | -10 | -92 | -8 | 3.44 | |

| | | | | | | | | | | |
|---|---|---|---|---|---|---|---|---|---|---|
| | | | | | Inferior frontal gyrus | -50 | 24 | -2 | -4.47 | |
| | | | | | Supramarginal_R (aal)* | 66 | -44 | 30 | -4.18 | |
| | | | | | Middle frontal gyrus | -40 | 38 | 24 | -3.58 | |
| (5) | $n_1$=81<br>$n_2$=81 | $n_1$: 21.19±1.85<br>$n_2$: 20.88±2.16 | $n_1$: 32/49<br>$n_2$: 32/49 | $p < .001$, corrected | DLPFC | -39 | 45 | 5 | -8.23 | 3.14 |
| | | | | | Hippocampus | 21 | -5 | -20 | 5.1 | |
| | | | | | Orbital frontal cortex | -2 | 33 | -14 | 4.92 | |

**S5 Table S2:** Significant Clusters Identified in the Uncorrected Procrastination Meta-analysis.

| Brain Region | MNI Coordinate | SDM-Z | P | Voxels |
|---|---|---|---|---|
| Blobs of ≥ 113 voxels with all voxels SDM-Z ≥ 1.645 and all peaks SDM-Z ≥ 1.878 | | | | |
| Right parahippocampal gyrus, BA 36 | 20,-10,-30 | 2.61 | 0.005 | 252 |
| Left superior frontal gyrus, medial orbital, BA 11 | 0,46,-10 | 1.88 | 0.030 | 113 |
| Blobs of ≥ 10 voxels with all voxels SDM-Z ≤ -1.646 and all peaks SDM-Z ≤ -1.979 | Show / Hide | | | |
| Left middle frontal gyrus, BA 46 | -34,42,30 | -3.23 | 0.001 | 432 |
| Left inferior frontal gyrus, triangular part, BA 45 | -48,36,-2 | -1.98 | 0.024 | 10 |

**S5 Table S3:** Significant Activation Clusters Identified in the NeuroSynth Meta-Analytic Map for Impulsivity.

| Cluster | X | Y | Z | Peak Intensity | Number of Voxels | Main Brain Region |
|---|---|---|---|---|---|---|
| 1 | 10 | 14 | -4 | 25.08 | 3204 | Right caudate |
| 2 | 50 | 10 | 26 | 15.02 | 265 | Right inferior frontal gyrus |
| 3 | 34 | -58 | 48 | 13.17 | 178 | Right angular gyrus |
| 4 | -46 | 8 | 26 | 12.31 | 214 | Left inferior frontal gyrus |
| 5 | -44 | -60 | 40 | 11.44 | 89 | Left inferior parietal lobule |
| 6 | 30 | -92 | 6 | 10.68 | 70 | Right middle occipital gyrus |
| 7 | -40 | 46 | 4 | 10.68 | 57 | Left inferior frontal gyrus |
| 8 | 46 | -48 | 52 | 10.57 | 81 | Right inferior parietal lobule |
| 9 | 50 | 30 | 30 | 9.60 | 30 | Right middle frontal gyrus |
| 10 | -14 | 64 | 16 | 6.64 | 24 | Left superior frontal gyrus |
| 11 | 38 | 22 | 44 | 6.64 | 98 | Right middle frontal gyrus |
| 12 | -58 | -46 | 4 | 5.78 | 56 | Left middle temporal gyrus |
| 13 | -34 | 46 | 26 | 5.77 | 23 | Left middle frontal gyrus |

*Note*: Only clusters with size greater than 20 voxels are reported.

**S5 Table S4:** Shared Neural Mask Between Impulsivity and Procrastination.

| Cluster | X | Y | Z | Peak Intensity | Number of Voxels | Main Brain Region |
|---|---|---|---|---|---|---|
| 1 | -40 | 42 | 20 | 2.43 | 6 | Middle frontal gyrus |
| 2 | -32 | 46 | 26 | 2.27 | 3 | Middle frontal gyrus |
| 3 | -42 | 44 | 26 | 2.07 | 3 | Middle frontal gyrus |
| 4 | 0 | 46 | -10 | 1.88 | 53 | Medial frontal gyrus |
| 5 | -44 | 38 | 20 | 1.71 | 1 | Middle frontal gyrus |
| 6 | -6 | 44 | -16 | 1.67 | 1 | Medial frontal gyrus |

### S6 Supplementary Analyses of Morphological Deviations in Shared Neural Loci: DLPFC and mPFC

To determine whether regional morphological deviations in cortical thickness (CT) and surface area (SA) significantly diverged from normative expectations, we conducted one-sample *t-tests* comparing mean deviation scores to a theoretical mean of zero. These analyses were performed independently for each of the 68 cortical regions defined by the Desikan–Killiany atlas. All statistical tests were carried out using *MATLAB's Statistical Toolbox (R2023b; ttest function)*, ensuring computational consistency and precision across regions. To account for multiple comparisons, the Benjamini–Hochberg procedure was applied to control the false discovery rate (FDR) at $q < 0.05$. In this cohort, 18 cortical regions demonstrated significant overdevelopment and 25 showed significant underdevelopment in CT morphology (S6 Table S**1**). For SA morphology, 18 regions were identified as significantly overdeveloped and 16 as underdeveloped (S6 Table S**2**). At the individual level, $Z$ scores ≥ 1.96 and ≤ −1.96 were used as thresholds to classify overdevelopment and underdevelopment, respectively. The proportion of participants (N = 140) exhibiting regional morphological deviations beyond these thresholds is also summarized in S6 Tables **S1–S2**. Given that this analysis was conducted on the same cohort as the main study, the observed findings align closely with those reported in the primary results (6).

**S6 Table S1:** Group-averaged deviations on CT for each region in this cohort.

| Brain Parcel | *t* value | *p* value | *BH q* value | Proportion |
|---|---|---|---|---|
| L_pericalcarine | 20.485 | 0 | 0 | 0.0000 |
| R_pericalcarine | 19.054 | 0 | 0 | 0.0000 |
| L_lingual | 14.187 | 0 | 0 | 0.0000 |
| R_lateraloccipital | 13.034 | 0 | 0 | 0.0000 |
| L_entorhinal | 12.789 | 0 | 0 | 0.0000 |
| L_lateraloccipital | 12.267 | 0 | 0 | 0.0000 |
| R_lingual | 11.684 | 0 | 0 | 0.0000 |
| R_cuneus | 9.923 | 0 | 0 | 0.7143 |
| R_entorhinal | 8.654 | 0 | 0 | 0.7143 |
| R_fusiform | 7.872 | 0 | 0 | 0.0000 |
| R_precentral | 6.792 | 0 | 0 | 1.4286 |
| R_caudalmiddlefrontal | 6.546 | 0 | 0 | 0.7143 |
| R_paracentral | 6.022 | 0 | 0 | 0.0000 |
| L_cuneus | 5.908 | 0 | 0 | 0.7143 |
| L_medialorbitofrontal | 5.497 | 0 | 0 | 0.0000 |
| L_inferiortemporal | 5.474 | 0 | 0 | 1.4286 |
| L_frontalpole | 4.663 | 0 | 0 | 2.8571 |
| L_fusiform | 3.582 | 0 | 0 | 1.4286 |
| L_precentral | 3.077 | 0.003 | 0.054 | 0.0000 |
| L_rostralmiddlefrontal | 2.676 | 0.008 | 0.144 | 0.7143 |
| L_temporalpole | 2.604 | 0.01 | 0.15 | 1.4286 |
| R_temporalpole | 2.427 | 0.016 | 0.224 | 2.1429 |
| L_paracentral | 2.37 | 0.019 | 0.2327 | 1.4286 |

| | | | | |
|---|---|---|---|---|
| R_medialorbitofrontal | 1.782 | 0.077 | 0.514 | 0.7143 |
| R_superiorparietal | 1.729 | 0.086 | 0.515 | 0.0000 |
| R_rostralmiddlefrontal | 1.639 | 0.104 | 0.52 | 4.2857 |
| R_transversetemporal | 1.311 | 0.192 | 0.576 | 0.7143 |
| L_caudalanteriorcingulate | 0.942 | 0.348 | 0.696 | 2.1429 |
| L_superiorparietal | 0.853 | 0.395 | 0.79 | 4.2857 |
| L_caudalmiddlefrontal | 0.819 | 0.414 | 0.828 | 2.1429 |
| R_parahippocampal | 0.768 | 0.444 | 0.888 | 0.7143 |
| L_parahippocampal | 0.486 | 0.628 | 1 | 2.1429 |
| L_rostralanteriorcingulate | -0.07 | 0.944 | 0.94 | 2.1429 |
| R_postcentral | -0.323 | 0.747 | 0.77 | 1.4286 |
| R_lateralorbitofrontal | -0.649 | 0.518 | 0.55 | 0.7143 |
| R_rostralanteriorcingulate | -0.685 | 0.494 | 0.54 | 3.5714 |
| L_transversetemporal | -0.823 | 0.412 | 0.47 | 1.4286 |
| L_postcentral | -1.348 | 0.18 | 0.21 | 1.4286 |
| R_caudalanteriorcingulate | -1.562 | 0.121 | 0.15 | 2.8571 |
| R_frontalpole | -1.669 | 0.097 | 0.12 | 7.8571 |
| R_inferiortemporal | -1.842 | 0.068 | 0.09 | 6.4286 |
| L_inferiorparietal | -1.942 | 0.054 | 0.07 | 0.7143 |
| L_superiorfrontal | -2.087 | 0.039 | 0.05 | 1.4286 |
| R_parstriangularis | -2.482 | 0.014 | 0.02 | 2.1429 |
| R_parsorbitalis | -2.498 | 0.014 | 0.02 | 0.7143 |
| R_superiorfrontal | -2.578 | 0.011 | 0.02 | 2.1429 |
| L_insula | -2.936 | 0.004 | 0.01 | 6.4286 |
| R_insula | -3.908 | 0 | 0.00 | 2.8571 |
| R_inferiorparietal | -4.995 | 0 | 0.00 | 1.4286 |
| L_parsorbitalis | -5.268 | 0 | 0.00 | 3.5714 |
| L_parstriangularis | -5.557 | 0 | 0.00 | 7.1429 |
| R_posteriorcingulate | -5.581 | 0 | 0.00 | 5.0000 |
| L_supramarginal | -5.634 | 0 | 0.00 | 4.2857 |
| L_bankssts | -5.7 | 0 | 0.00 | 5.7143 |
| R_precuneus | -6.62 | 0 | 0.00 | 2.8571 |
| R_parsopercularis | -6.632 | 0 | 0.00 | 4.2857 |
| L_lateralorbitofrontal | -7.403 | 0 | 0.00 | 7.8571 |
| L_middletemporal | -7.595 | 0 | 0.00 | 7.1429 |
| L_superiortemporal | -7.664 | 0 | 0.00 | 7.1429 |
| R_bankssts | -8.261 | 0 | 0.00 | 8.5714 |
| L_posteriorcingulate | -9.584 | 0 | 0.00 | 8.5714 |
| R_superiortemporal | -10.297 | 0 | 0.00 | 9.2857 |

| | | | | |
|---|---|---|---|---|
| R_supramarginal | -11.024 | 0 | 0.00 | 11.4286 |
| R_isthmuscingulate | -11.673 | 0 | 0.00 | 7.8571 |
| L_parsopercularis | -12.445 | 0 | 0.00 | 11.4286 |
| R_middletemporal | -12.842 | 0 | 0.00 | 20.7143 |
| L_precuneus | -13.089 | 0 | 0.00 | 10.0000 |
| L_isthmuscingulate | -14.243 | 0 | 0 | 15.7143 |

**S6 Table S2:** Group-averaged deviations on SA for each region in this cohort.

| Brain Parcel | *t* value | *p* value | *BH q* value | Proportion |
|---|---|---|---|---|
| R_supramarginal | 8.722 | 0 | 0.0000 | 0.0000 |
| R_medialorbitofrontal | 7.729 | 0 | 0.0000 | 0.0000 |
| R_insula | 7.667 | 0 | 0.0000 | 0.0000 |
| R_isthmuscingulate | 7.488 | 0 | 0.0000 | 0.7143 |
| L_inferiortemporal | 5.641 | 0 | 0.0000 | 1.4286 |
| R_paracentral | 5.42 | 0 | 0.0000 | 0.0000 |
| L_parsopercularis | 5.368 | 0 | 0.0000 | 0.0000 |
| L_precentral | 4.806 | 0 | 0.0000 | 0.7143 |
| R_lingual | 4.472 | 0 | 0.0000 | 1.4286 |
| L_frontalpole | 4.291 | 0 | 0.0000 | 1.4286 |
| L_precuneus | 4.167 | 0 | 0.0000 | 0.7143 |
| L_cuneus | 4.11 | 0 | 0.0000 | 0.0000 |
| L_superiorparietal | 3.763 | 0 | 0.0000 | 0.0000 |
| R_fusiform | 3.298 | 0.001 | 0.0028 | 1.4286 |
| L_paracentral | 2.996 | 0.003 | 0.0078 | 0.7143 |
| R_caudalmiddlefrontal | 2.911 | 0.004 | 0.0098 | 1.4286 |
| R_bankssts | 2.861 | 0.005 | 0.0115 | 2.1429 |
| L_pericalcarine | 2.786 | 0.006 | 0.0130 | 0.0000 |
| L_caudalanteriorcingulate | 2.24 | 0.027 | 0.0554 | 0.7143 |
| L_supramarginal | 2.119 | 0.036 | 0.0702 | 1.4286 |
| L_middletemporal | 1.999 | 0.048 | 0.0891 | 2.1429 |
| L_entorhinal | 1.859 | 0.065 | 0.1152 | 1.4286 |
| L_isthmuscingulate | 1.808 | 0.073 | 0.1238 | 0.0000 |
| L_posteriorcingulate | 1.719 | 0.088 | 0.1430 | 0.7143 |
| R_temporalpole | 1.592 | 0.114 | 0.1725 | 3.5714 |
| R_frontalpole | 1.586 | 0.115 | 0.1725 | 1.4286 |
| R_parahippocampal | 1.463 | 0.146 | 0.2075 | 0.7143 |
| R_pericalcarine | 1.453 | 0.149 | 0.2075 | 0.0000 |
| L_insula | 1.356 | 0.177 | 0.2380 | 1.4286 |
| L_parstriangularis | 1.322 | 0.188 | 0.2415 | 0.7143 |

| | | | | |
|---|---|---|---|---|
| R_transversetemporal | 1.312 | 0.192 | 0.2415 | 0.0000 |
| R_parstriangularis | 0.991 | 0.323 | 0.3937 | 2.8571 |
| L_rostralmiddlefrontal | 0.865 | 0.388 | 0.4585 | 5.7143 |
| L_parsorbitalis | 0.845 | 0.4 | 0.4588 | 2.8571 |
| R_precuneus | 0.471 | 0.639 | 0.7120 | 0.7143 |
| L_lingual | 0.31 | 0.757 | 0.8201 | 1.4286 |
| R_superiorfrontal | 0.191 | 0.849 | 0.8949 | 1.4286 |
| L_inferiorparietal | 0.066 | 0.948 | 0.9550 | 2.1429 |
| R_entorhinal | 0.057 | 0.955 | 0.9550 | 3.5714 |
| L_parahippocampal | -0.356 | 0.723 | 0.7230 | 5.0000 |
| L_fusiform | -0.623 | 0.534 | 0.5531 | 5.0000 |
| L_lateralorbitofrontal | -0.737 | 0.462 | 0.4962 | 2.1429 |
| R_precentral | -0.913 | 0.363 | 0.4049 | 1.4286 |
| R_rostralanteriorcingulate | -0.917 | 0.361 | 0.4049 | 1.4286 |
| R_rostralmiddlefrontal | -1.155 | 0.25 | 0.3021 | 2.1429 |
| L_rostralanteriorcingulate | -1.185 | 0.238 | 0.3001 | 2.1429 |
| R_superiorparietal | -1.194 | 0.235 | 0.3001 | 3.5714 |
| L_temporalpole | -1.226 | 0.222 | 0.3001 | 2.1429 |
| R_postcentral | -1.233 | 0.22 | 0.3001 | 5.7143 |
| L_medialorbitofrontal | -1.862 | 0.065 | 0.0992 | 4.2857 |
| R_parsorbitalis | -1.965 | 0.051 | 0.0822 | 2.8571 |
| R_inferiortemporal | -2.061 | 0.041 | 0.0699 | 2.8571 |
| R_parsopercularis | -2.695 | 0.008 | 0.0145 | 7.1429 |
| L_transversetemporal | -2.901 | 0.004 | 0.0077 | 3.5714 |
| L_superiortemporal | -2.914 | 0.004 | 0.0077 | 3.5714 |
| R_middletemporal | -3.068 | 0.003 | 0.0067 | 4.2857 |
| R_inferiorparietal | -3.229 | 0.002 | 0.0048 | 1.4286 |
| L_caudalmiddlefrontal | -3.791 | 0 | 0.0000 | 5.0000 |
| R_cuneus | -4.052 | 0 | 0.0000 | 2.8571 |
| R_lateralorbitofrontal | -4.421 | 0 | 0.0000 | 3.5714 |
| R_lateraloccipital | -4.662 | 0 | 0.0000 | 5.0000 |
| L_bankssts | -4.855 | 0 | 0.0000 | 6.4286 |
| R_superiortemporal | -5.94 | 0 | 0.0000 | 2.8571 |
| L_lateraloccipital | -6.597 | 0 | 0.0000 | 7.1429 |
| L_postcentral | -7.079 | 0 | 0.0000 | 7.1429 |
| L_superiorfrontal | -8.96 | 0 | 0.0000 | 3.5714 |
| R_posteriorcingulate | -10.058 | 0 | 0.0000 | 9.2857 |
| R_caudalanteriorcingulate | -10.577 | 0 | 0.0000 | 22.1429 |

### S7 Genomic Analysis and Causal Inference Methods

#### S7.1 Whole-genome sequence for the adult cohort

In the study cohort, a total of 936 samples underwent whole-genome sequencing with a mean sequencing depth of 27×. Genomic DNA was obtained from the BGI-genomics. DNA was prepared with QIAGEN DNeasy Blood & Tissue Kit. Library construction was performed at BGI-genomics. The qualified genomic DNA sample was randomly fragmented by Covaris technology, and the DNA fragments were selected by size. The end-repair of DNA fragments was added an 'A' base at the 3′-end of each strand. DNBSEQ adapters were ligated to both ends of the A-tailed fragments, followed by amplification by ligation-mediated PCR (LM-PCR), single strand separation and cyclization. The rolling circle amplification (RCA) was performed to produce DNA Nanoballs (DNBs). The qualified DNBs were loaded into the patterned nanoarrays and processed for 150 bp pair-end sequencing on the DNBSEQ platform. Sequencing-derived raw image files were processed by the DNBSEQ base calling software with default parameters.

All raw data were filtered using SOAPnuke (v.2.2.1) (7) for quality trimming. Subsequently, the clean reads from each sample were aligned to the human reference genome hg38 using Burrows Wheeler Aligner (BWA) (8). SNP/InDel calling and annotation were performed using GATK (v.4.1.4.1) (9).

The Base Recalibrator to generate recalibration tables for identification of known SNPs and insertions or deletions (INDELs) within the BAM files obtained from dbSNP (v.150). Subsequently, GATKlite was employed for subsequent base quality recalibration and removal of read pairs with improperly aligned segments, determined by Stampy. Variant discovery was conducted using GATK's HaplotypeCaller. The GVCFs containing SNVs and INDELs from the GATK HaplotypeCaller output were combined, genotyped, variant score recalibrated, and filtered through a series of steps, including CombineGVCFs, GenotypeGVCFs, VariantRecalibrator, and ApplyRecalibration.

In the VariantRecalibrator process, the variants served as inputs, and the model was trained using four standard SNP sets: (1) HapMap3.3 SNPs; (2) dbSNP build 150 SNPs; (3) 1000 Genomes Project SNPs from Omni 2.5 chip; and (4) 1000 G phase1 high confidence SNPs. Variant selection was optimized for Transition to Transversion (TiTv) ratios, with sensitivity thresholds of 99.9% for SNPs and 99% for INDELs applied after recalibration using the GATK ApplyRecalibration command. Post-recalibration, 53 million raw variants remained, comprising 43,602,291 SNPs and 9,524,247 INDELs.

A conservative inclusion threshold was applied for variants, including criteria such as mean depth >8×, Hardy-Weinberg equilibrium (HWE) $P > 10^{-6,}$ and genotype calling rate > 90%. Samples were required to meet criteria including mean sequencing depth > 20×, variant calling rate > 90%, absence of population stratification determined by principal components analysis (PCA) implemented in PLINK50 (v.1.9) (10), and exclusion of related individuals based on pairwise identity by descent (IBD) calculation (Pi-hat threshold of 0.1875) in PLINK. After sample matching, variant and sample quality control, a total of 835 individuals with 6.19 million common variants (MAF ≥ 5%) and 3.64 million low-frequency variants (0.5% ≤ MAF < 5%) from the discovery cohort remained for subsequent GWAS analysis.

#### S7.2 GWAS Data source

We first performed a Genome-Wide Association Study (GWAS) on both procrastination and impulsivity within the large-scale adult cohort. After sample matching, variant and sample quality control, a total of 835 individuals with 6.19 million common variants (MAF ≥ 5%) and 3.64 million low-frequency variants (0.5% ≤ MAF < 5%). GWAS analyses were conducted separately for procrastination scores (GPS scale) and non-planning impulsivity scores (BIS-11 scale), using sex, age, and the top 10 ancestry principal components as covariates. This analysis enabled us to identify genetic influences underlying impulsivity and procrastination and explore their potential molecular underpinnings.

Genome-wide significant associations related to DLPFC morphology were obtained from a recently published genome-wide association study on brain imaging phenotypes (11). This study conducted a comprehensive GWAS in a cohort of 7058 Chinese Han (CH) individuals from the Chinese Imaging Genetics (CHIMGEN) study, examining 3414 brain phenotypes. The gene associations were adjusted for covariates including age, sex, age*sex interaction, age-squared, and the top three genetic principal components specific to the CH-GWAS. The brain imaging phenotypes analyzed in this study included the T1-derived FreeSurfer DK atlas-based rostral middle frontal cortical surface area and thickness in both hemispheres. The GWAS summary statistics of the DLPFC morphology based on CH-GWAS are obtained from GWAS Catalog (https://www.ebi.ac.uk/gwas/home ).

#### S7.3 Mendelian Randomization Methods

In this study, we employed several complementary Mendelian Randomization (MR) methods to estimate causal relationships and ensure the robustness of our findings. These methods differ in their underlying assumptions regarding the validity of instrumental variables, the presence of horizontal pleiotropy, and the proportion or clustering of valid SNPs, allowing for a more comprehensive evaluation of potential causal effects under varying conditions.

The inverse-variance weighted (IVW) method served as the primary analytical approach. The IVW method assumes that all SNPs are valid and there is no horizontal pleiotropy and that the SNP-specific effects are homogenous. It is based on weighted linear regression, where the association of SNP $i$ with the outcome ($\beta_{SNP\rightarrow outcome,i}$) is regressed on its association with the exposure ($\beta_{SNP\rightarrow exposure,i}$). The inverse variance of the SNP-outcome effect of SNP $i$ is used as the weight ($\omega_i$), giving more influence to SNPs with more precise estimates (f1). By combining the individual SNP effects, the IVW method provides an overall causal estimate of the exposure on the outcome. The causal estimate is computed as followed (f2) while *n* is the total number of SNPs:

$$\omega_i = \frac{1}{\mathrm{SE}^2_{\mathrm{SNP\rightarrow outcome,i}}} \quad \text{(f1)}$$

$$\beta_{\mathrm{IVW}} = \frac{\sum_{i=1}^{n} \omega_i * \frac{\beta_{SNP\rightarrow outcome,i}}{\beta_{SNP\rightarrow exposure,i}}}{\sum_{i=1}^{n} \omega_i} \quad \text{(f2)}$$

The MR-Egger regression method was employed as a complementary analytical approach to account for horizontal pleiotropy. Similar to the IVW method, MR-Egger is based on a weighted linear regression framework, where the association of SNP $i$ with the outcome ($\beta_{SNP\rightarrow outcome,i}$) is regressed on its association with the exposure ($\beta_{SNP\rightarrow exposure,i}$). Each SNP's weight ($\omega_i$) is also calculated as the inverse variance of the SNP-outcome effect (f1). However, unlike the IVW method, MR-Egger does not constrain the intercept to zero, which means that it allows for the presence of horizontal pleiotropy. The intercept ($\alpha$) provides an estimate of the average pleiotropic effect across all SNPs, while the slope ($\beta$) represents the causal effect of the exposure on the outcome after adjusting for pleiotropy. By allowing a non-zero intercept, MR-Egger can detect and adjust for pleiotropic effects under the InSIDE assumption (instrument strength independent of direct effects). The MR-Egger regression model is defined as followed (f3) while $\in_i$ is the residual error for SNP $i$. When using Weighted Least Squares (WLS) for parameter estimation, the formulas can be expressed as followed (f4 and f5). However, MR-Egger has lower statistical power and precision compared to IVW, particularly in the presence of weak instruments; therefore, it requires a larger sample size and a greater number of valid SNPs to ensure reliable causal inference and to mitigate the impact of random error on the estimates.

$$\beta_{SNP\rightarrow outcome,i} = \alpha + \beta * \beta_{SNP\rightarrow exposure,i} + \in_i \quad \text{(f3)}$$

$$\hat{\beta} = \frac{\sum_{i=1}^{n} \omega_i * \beta_{SNP\rightarrow exposure,i} * \beta_{SNP\rightarrow outcome,i}}{\sum_{i=1}^{n} \omega_i * \beta^2_{SNP\rightarrow exposure,i}} \quad \text{(f4)}$$

$$\hat{\alpha} = \frac{\sum_{i=1}^{n} \omega_i * (\beta_{SNP\rightarrow outcome,i} - \hat{\beta} * \beta_{SNP\rightarrow exposure,i})}{\sum_{i=1}^{n} \omega_i} \quad \text{(f5)}$$

The weighted median estimator method assumes that at least 50% of the total weight comes from valid SNPs, which satisfy the relevance, independence, and exclusion restriction assumptions of MR. Then the weighted median is defined as the value at which the cumulative weight reaches 50% when SNP-specific causal effect estimates ($\beta_i$) are ordered from smallest to largest (f6 and f7). Each SNP's weight ($\omega_i$) is also calculated as the inverse variance of the SNP-outcome effect (f1). This method is particularly useful in settings where heterogeneity or pleiotropy is suspected, as it provides a robust causal estimate without requiring all SNPs to be valid instruments as long as the majority of the total weight comes from valid SNPs. However, it may lose efficiency compared to IVW when all SNPs are valid, as it does not fully leverage the precision of all SNPs. The weighted median estimator ($\beta_{WM}$) is computed as followed:

$$\beta_i = \frac{\beta_{SNP \to outcome,i}}{\beta_{SNP \to exposure,i}} \text{ (f6)}$$

$$\beta_{WM} = \text{Median}(\beta_1, \beta_2, \ldots, \beta_n \,; \omega_1, \omega_2, \ldots, \omega_n) \text{ (f7)}$$

The weighted mode estimator is designed to estimate causal effects by identifying the most frequent (modal) SNP-specific causal effect in the distribution of all SNPs. It assumes that the largest group (mode) of SNPs with similar causal effects represents valid instruments, while SNPs with pleiotropic or invalid effects are distributed outside this main cluster. This method is particularly useful when pleiotropy or heterogeneity is suspected, as it allows for the presence of invalid SNPs, provided the valid SNPs dominate the mode. By focusing on the mode, the weighted mode estimator minimizes the influence of invalid instruments and outliers, ensuring robustness against pleiotropy. While robust to pleiotropy, because the weighted mode estimator relies on the assumption that the majority of valid SNPs share a similar causal effect, it may lose efficiency compared to other methods, such as IVW, when all SNPs are valid instruments. Similar to the weighted median estimator method, the weighted mode estimator uses SNP-specific causal effect estimates (f6) but uses the inverse variance of SNP-exposure effects to identify the modal cluster of causal effect estimates (f8). The causal estimate is derived as the mode of the weighted distribution of these estimates. The weighted mode causal estimate ($\beta_{WM}$) is computed as followed (f9):

$$\omega_i = \frac{1}{\text{SE}^2_{\text{SNP}\to\text{exposure,i}}} \text{ (f8)}$$

$$\beta_{WM} = \text{Mode}(\beta_1, \beta_2, \ldots, \beta_n \,; \omega_1, \omega_2, \ldots, \omega_n) \text{ (f9)}$$

The results from these four complementary MR methods: IVW, MR-Egger regression, weighted median estimator, and weighted mode estimator, including the causal estimates ($\beta$), 95% confidence intervals (*CI*), and *p-values*, are summarized in S7 Table S1. To further illustrate the results, Figure S7 provides scatter plots showing the SNP-exposure and SNP-outcome associations, along with the fitted causal effect lines derived from the four MR methods for each exposure-outcome pair. Among these methods, only the IVW method yielded significant results, the other MR methods (MR-Egger, weighted median, and weighted mode) produced consistent but statistically non-significant causal estimates. This discrepancy likely reflects the reduced statistical power of the alternative methods compared to IVW, particularly in the presence of weak instruments or a limited number of SNPs. Therefore, IVW remains the most reliable approach for detecting significant causal effects under standard MR assumptions, providing robust evidence to support the observed associations.

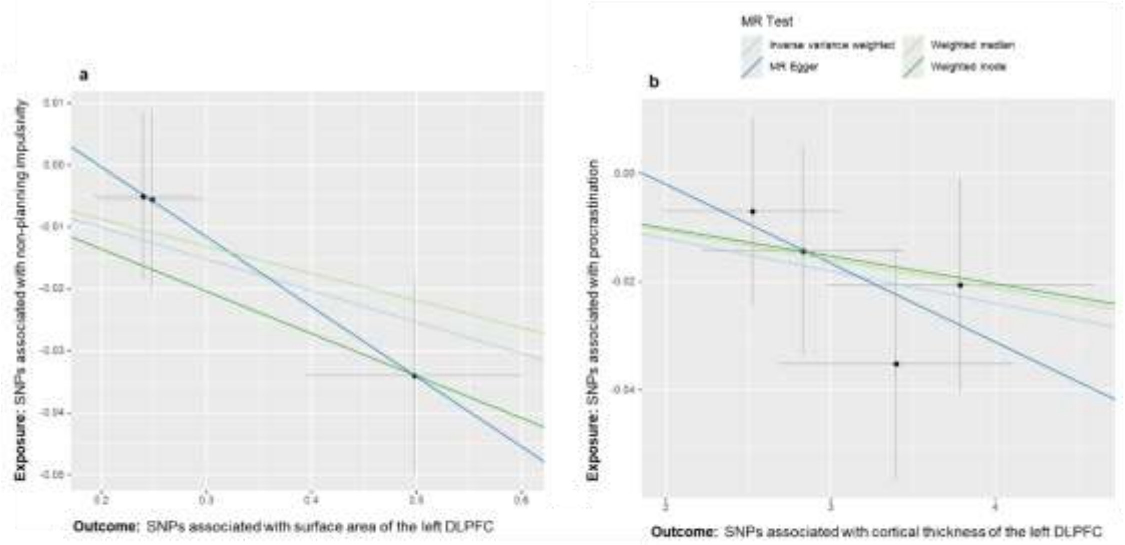

**Figure S7** The causal relationship between non-planning impulsivity and the surface area of the left DLPFC. (a), procrastination and the cortical thickness of the left DLPFC (b).

To further evaluate the robustness of the causal relationship between the exposure variables and the outcome variables, leave-one-out sensitivity analysis was performed. This analysis involved iteratively removing each SNP associated with impulsivity and recalculating the causal effect estimate using the remaining SNPs. The results are summarized in the accompanying plots. The red point ("All") represents the overall causal effect estimate and its 95% confidence interval (CI) when all SNPs were included. The black points represent the causal effect estimates and their 95% CIs after removing each SNP individually. The leave-one-out sensitivity analysis for these analyses further supported the robustness of the results, as removing any single SNP did not lead to substantial changes in the causal estimates. All estimates remained negative and clustered closely around the overall effect (Figure S8).

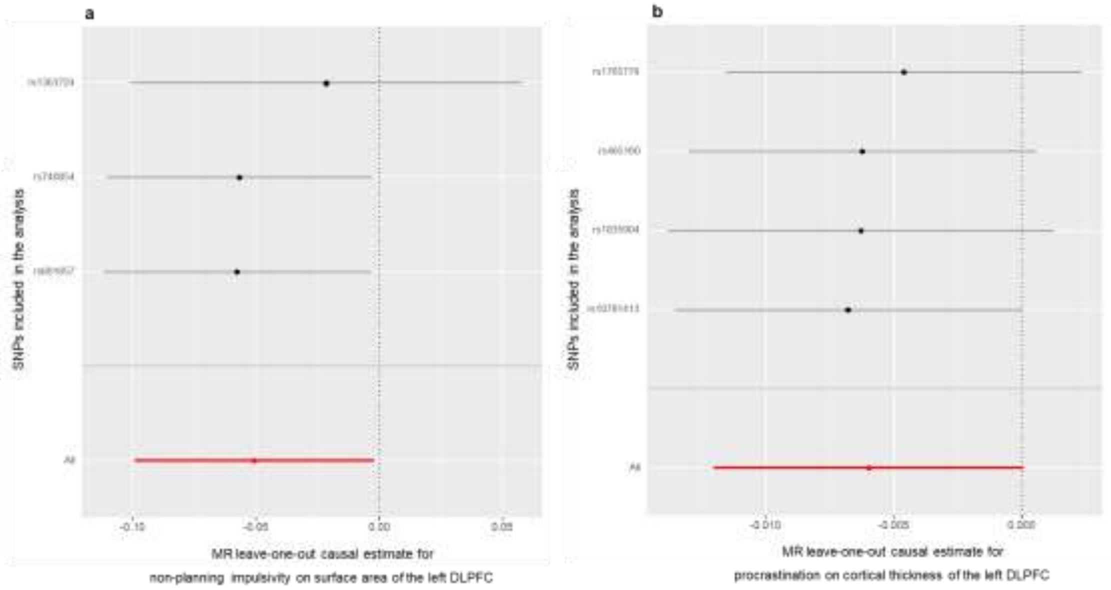

**Figure S8** Leave-one-out sensitivity analysis for the causal relationship between non-planning impulsivity and the surface area of the left DLPFC (a), procrastination and the cortical thickness of the left DLPFC (b)

**S7 Table S1:** Mendelian randomization results showing causal estimates for each exposure-outcome pair using four MR methods.

| Exposure | Outcome | Number of SNPs | Method | beta | 95% CI | p-value |
|---|---|---|---|---|---|---|
| non-planning impulsivity | the surface area of left DLPFC | 3 | IVW | -0.05 | (-0.10, -0.0020) | 0.041 |
| | | | MR-Egger regression | -0.11 | (-0.26, 0.030) | 0.364 |
| | | | Weighted median estimator | -0.04 | (-0.10, 0.015) | 0.145 |
| | | | Weighted mode estimator | -0.07 | (-0.13, -0.0066) | 0.162 |
| procrastination | the cortical thickness of left DLPFC | 4 | IVW | -0.01 | (-0.012, 6.44e-05) | 0.052 |
| | | | MR-Egger regression | -0.01 | (-0.052, 2.33e-02) | 0.530 |
| | | | weighted median estimator | -0.01 | (-0.012, 2.02e-03) | 0.156 |
| | | | weighted mode estimator | -0.01 | (-0.014, 4.14e-03) | 0.359 |

**S7 Table S2:** Candidate Genetic Loci for Procrastination Identified at $p < 5 \times 10^{-5}$.

| | SNP | Chromosome: start | Effect Allele | Other Allele | MAF | beta | pval | se |
|---|---|---|---|---|---|---|---|---|
| 1 | rs9305375 | 21:28619012 | C | G | 0.2305 | -3.264 | 8.835E-07 | 0.658595642 |
| 2 | rs1765779 | 1:231598827 | C | T | 0.1747 | -3.396 | 0.000002273 | 0.712846348 |
| 3 | rs12739703 | 1:217661602 | C | G | 0.0896 | 4.384 | 0.000002774 | 0.928616818 |
| 4 | rs7555956 | 1:217662005 | G | A | 0.09085 | 4.308 | 0.00000372 | 0.924463519 |
| 5 | rs10781413 | 9:77069932 | C | T | 0.4836 | -2.524 | 0.000003754 | 0.541863461 |
| 6 | rs465160 | 5:25843368 | C | T | 0.245 | -2.833 | 0.000004121 | 0.610823631 |
| 7 | rs1835904 | 5:174261696 | G | A | 0.3802 | 3.788 | 0.00000421 | 0.81096125 |
| 8 | rs5017213 | 1:217720162 | C | T | 0.08897 | 4.311 | 0.000004212 | 0.930498597 |
| 9 | rs801777 | 5:25842159 | T | C | 0.245 | -2.809 | 0.000004679 | 0.609195402 |
| 10 | rs6671096 | 1:217706641 | C | A | 0.1037 | 4.359 | 0.000005566 | 0.951538965 |
| 11 | rs1568425 | 3:99257875 | C | A | 0.1053 | 7.538 | 0.00000564 | 0.375024876 |
| 12 | rs1568424 | 3:99257876 | G | T | 0.1053 | 7.538 | 0.00000564 | 0.375024876 |
| 13 | rs387140 | 5:25844755 | C | T | 0.2456 | -2.803 | 0.000005673 | 0.613347921 |
| 14 | rs17724545 | 1:217662757 | C | T | 0.08908 | 4.247 | 0.000005848 | 0.930747315 |
| 15 | rs1025527 | 1:231614099 | G | A | 0.1723 | -3.232 | 0.000006295 | 0.710798329 |
| 16 | rs465365 | 5:25843875 | T | C | 0.2465 | -2.779 | 0.000006446 | 0.611845002 |
| 17 | rs11740209 | 5:25840027 | C | T | 0.2513 | -2.739 | 0.000006456 | 0.603038309 |
| 18 | rs7537954 | 1:217685966 | T | A | 0.09023 | 4.207 | 0.000006548 | 0.926856136 |
| 19 | rs67714799 | 13:49478319 | G | A | 0.1401 | 3.479 | 0.000006931 | 0.76850011 |
| 20 | rs406594 | 5:25844809 | G | A | 0.2469 | -2.764 | 0.000007142 | 0.611504425 |
| 21 | rs455792 | 5:25845121 | G | C | 0.2469 | -2.764 | 0.000007142 | 0.611504425 |

| | | | | | | | | |
|---|---|---|---|---|---|---|---|---|
| 22 | rs455846 | 5:25845231 | A | C | 0.2469 | -2.764 | 0.000007142 | 0.611504425 |
| 23 | rs455710 | 5:25845276 | G | A | 0.2469 | -2.764 | 0.000007142 | 0.611504425 |
| 24 | rs462495 | 5:25845672 | A | T | 0.2469 | -2.764 | 0.000007142 | 0.611504425 |
| 25 | rs398513 | 5:25836129 | G | A | 0.2434 | -2.76 | 0.000007991 | 0.614015573 |
| 26 | rs462672 | 5:25843268 | A | G | 0.2459 | -2.745 | 0.000008389 | 0.612040134 |
| 27 | rs9568210 | 13:49454900 | T | C | 0.1399 | 3.459 | 0.000008806 | 0.773133661 |
| 28 | rs464471 | 5:25843726 | T | A | 0.2462 | -2.733 | 0.000009085 | 0.611820013 |
| 29 | rs10757138 | 9:20547177 | A | G | 0.4426 | -2.43 | 0.000009167 | 0.544111061 |
| 30 | rs10065041 | 5:25840341 | T | C | 0.2509 | -2.739 | 0.000009209 | 0.61343785 |
| 31 | rs1232001 | 18:3523663 | C | T | 0.3408 | 2.592 | 0.000009219 | 0.580385132 |
| 32 | rs396045 | 5:25841504 | A | G | 0.2469 | -2.717 | 0.000009614 | 0.609876543 |
| 33 | rs457354 | 5:25844187 | G | A | 0.2459 | -2.735 | 0.000009633 | 0.613916947 |
| 34 | rs466188 | 5:25844207 | A | G | 0.2459 | -2.735 | 0.000009633 | 0.613916947 |
| 35 | rs9562880 | 13:49453710 | T | C | 0.1405 | 3.415 | 0.000009885 | 0.767588222 |
| 36 | rs2310368 | 4:185941822 | T | C | 0.06328 | 4.924 | 0.00001027 | 1.109009009 |
| **37** | **rs9325070** | **5:148097209** | **G** | **A** | **0.2989** | **-2.547** | **0.00001034** | **0.573777878** |
| **38** | **rs9791022** | **5:148097621** | **G** | **A** | **0.3026** | **-2.557** | **0.00001052** | **0.576550169** |
| 39 | rs2094709 | 9:77070282 | A | G | 0.4028 | 2.456 | 0.00001091 | 0.554777502 |
| 40 | rs117578951 | 12:54824399 | T | C | 0.05583 | 5.067 | 0.0000112 | 1.146120787 |
| 41 | rs117848353 | 12:54797366 | G | A | 0.05764 | 5.109 | 0.00001129 | 1.155882353 |
| 42 | rs3010550 | 6:167303140 | T | C | 0.3497 | 2.46 | 0.00001133 | 0.556687033 |
| 43 | rs9562882 | 13:49458965 | C | G | 0.1405 | 3.396 | 0.00001136 | 0.768673608 |
| 44 | rs17027918 | 12:98199072 | T | C | 0.1178 | -3.662 | 0.00001165 | 0.829820983 |

| | | | | | | | | |
|---|---|---|---|---|---|---|---|---|
| 45 | rs11109384 | 12:98199363 | T | C | 0.1178 | -3.662 | 0.00001165 | 0.829820983 |
| 46 | rs73064102 | 19:49749226 | T | C | 0.1284 | -3.465 | 0.00001179 | 0.785714286 |
| 47 | rs9568211 | 13:49455244 | A | G | 0.141 | 3.383 | 0.00001181 | 0.767120181 |
| 48 | rs9562881 | 13:49455648 | T | C | 0.141 | 3.383 | 0.00001181 | 0.767120181 |
| 49 | rs66771344 | 13:49458340 | A | G | 0.141 | 3.383 | 0.00001181 | 0.767120181 |
| 50 | rs9568216 | 13:49469278 | T | C | 0.141 | 3.383 | 0.00001181 | 0.767120181 |
| 51 | rs55995775 | 9:77068047 | G | C | 0.468 | 2.397 | 0.00001263 | 0.545392491 |
| 52 | rs8102970 | 19:49755480 | G | A | 0.1322 | -3.375 | 0.00001297 | 0.768967874 |
| 53 | rs72633255 | 16:534107 | C | T | 0.4519 | 2.332 | 0.00001361 | 0.532663317 |
| 54 | rs892356 | 1:231611204 | T | C | 0.1727 | -3.091 | 0.00001387 | 0.706675812 |
| 55 | rs2356384 | 1:231607935 | A | G | 0.1748 | -3.08 | 0.00001418 | 0.704966812 |
| 56 | rs7548847 | 1:231612243 | A | G | 0.1748 | -3.078 | 0.00001423 | 0.70467033 |
| 57 | rs9562884 | 13:49467265 | T | C | 0.1405 | 3.352 | 0.00001443 | 0.76792669 |
| 58 | rs9568209 | 13:49453566 | C | G | 0.1401 | 3.348 | 0.00001528 | 0.769301471 |
| 59 | rs444099 | 5:25826509 | C | A | 0.245 | -2.67 | 0.00001558 | 0.614075437 |
| 60 | rs35672711 | 9:77068726 | T | C | 0.4687 | 2.37 | 0.0000159 | 0.545705733 |
| 61 | rs9568215 | 13:49468847 | C | T | 0.1388 | 3.386 | 0.0000161 | 0.780184332 |
| 62 | rs415568 | 5:25830609 | G | A | 0.2469 | -2.641 | 0.00001612 | 0.608525346 |
| 63 | rs13428269 | 2:75168653 | T | C | 0.1405 | -3.39 | 0.00001623 | 0.781466113 |
| 64 | rs7025158 | 9:20542634 | C | T | 0.4428 | -2.36 | 0.00001633 | 0.544154946 |
| 65 | rs4555176 | 16:76450679 | T | A | 0.2792 | -2.723 | 0.00001637 | 0.627853355 |
| 66 | rs446987 | 5:25829446 | A | G | 0.2443 | -2.664 | 0.00001659 | 0.614674665 |
| 67 | rs17381926 | 13:49519954 | C | T | 0.141 | 3.312 | 0.00001769 | 0.766844177 |

| | | | | | | | | |
|---|---|---|---|---|---|---|---|---|
| 68 | rs10414681 | 19:49756465 | T | C | 0.1518 | -3.132 | 0.00001853 | 0.726850777 |
| 69 | rs4887979 | 16:78753994 | T | C | 0.3134 | 2.46 | 0.00001903 | 0.571827057 |
| 70 | rs6668915 | 1:231602144 | C | A | 0.1924 | -2.929 | 0.00001976 | 0.682114578 |
| 71 | rs439958 | 5:25818423 | G | C | 0.2443 | -2.642 | 0.0000199 | 0.61556384 |
| 72 | rs11791670 | 9:20545478 | G | A | 0.4404 | -2.343 | 0.00001997 | 0.545899348 |
| 73 | rs56737298 | 13:49489665 | T | C | 0.135 | 3.357 | 0.00002014 | 0.782517483 |
| 74 | rs3904084 | 1:225649736 | G | C | 0.1078 | -3.703 | 0.00002018 | 0.863371415 |
| 75 | rs249224 | 5:80552069 | C | G | 0.06901 | 4.396 | 0.00002021 | 1.02494754 |
| 76 | rs28506894 | 19:49750361 | G | C | 0.1531 | -3.088 | 0.00002083 | 0.721158337 |
| 77 | rs801778 | 5:25842057 | G | C | 0.2431 | -2.632 | 0.00002087 | 0.614666044 |
| 78 | rs11141289 | 9:86180304 | T | C | 0.06721 | -4.757 | 0.000021 | 1.111448598 |
| 79 | rs6674294 | 1:231602639 | C | T | 0.192 | -2.936 | 0.00002112 | 0.686141622 |
| 80 | rs12134398 | 1:231600584 | A | C | 0.193 | -2.917 | 0.0000212 | 0.681860683 |
| 81 | rs6679985 | 1:231602083 | C | G | 0.193 | -2.917 | 0.0000212 | 0.681860683 |
| 82 | rs6682685 | 1:231602464 | A | G | 0.193 | -2.917 | 0.0000212 | 0.681860683 |
| 83 | rs4070570 | 4:185942031 | A | G | 0.06391 | 4.721 | 0.00002175 | 1.105102996 |
| 84 | rs7529790 | 1:231612752 | A | C | 0.1705 | -3.026 | 0.00002219 | 0.708997188 |
| 85 | rs6656071 | 1:231601954 | T | C | 0.1926 | -2.911 | 0.00002265 | 0.682852451 |
| 86 | rs12228375 | 12:90535875 | A | C | 0.1913 | 2.998 | 0.00002342 | 0.704582844 |
| 87 | rs249225 | 5:80552148 | A | G | 0.06955 | 4.345 | 0.00002368 | 1.021631789 |
| 88 | rs6552487 | 4:181146944 | T | C | 0.3653 | -2.361 | 0.00002465 | 0.556445911 |
| 89 | rs11952614 | 5:80548073 | T | C | 0.06955 | 4.333 | 0.00002532 | 1.022657541 |
| 90 | rs10894795 | 11:100356042 | G | A | 0.06093 | 4.777 | 0.00002538 | 1.127448667 |

| | | | | | | | | |
|---|---|---|---|---|---|---|---|---|
| 91 | rs1655299 | 1:231596415 | T | C | 0.175 | -3.051 | 0.00002564 | 0.720595182 |
| 92 | rs150754743 | 3:137152185 | T | A | 0.148 | 4.115 | 0.00002742 | 0.971206042 |
| 93 | rs1562335 | 2:238236743 | C | T | 0.3526 | 2.448 | 0.00002787 | 0.580782918 |
| 94 | rs2001342 | 19:49757511 | T | C | 0.1566 | -3.004 | 0.00002945 | 0.714897668 |
| 95 | rs9568240 | 13:49532425 | G | T | 0.1385 | 3.233 | 0.00003004 | 0.770128633 |
| 96 | rs7596687 | 2:238236620 | C | T | 0.3473 | 2.423 | 0.00003006 | 0.577179609 |
| 97 | rs57893931 | 14:90078247 | T | C | 0.2375 | -2.648 | 0.00003155 | 0.632584806 |
| 98 | rs146068884 | 13:49504715 | G | A | 0.1212 | 3.292 | 0.00003311 | 0.788502994 |
| 99 | rs7867710 | 9:20602189 | C | T | 0.3526 | 2.315 | 0.00003328 | 0.554623862 |
| 100 | rs420870 | 5:25825083 | T | C | 0.2393 | -2.581 | 0.00003339 | 0.61849988 |
| 101 | rs73190705 | 13:49519868 | G | A | 0.1399 | 3.225 | 0.00003349 | 0.772825306 |
| 102 | rs7331182 | 13:49523754 | T | C | 0.1391 | 3.211 | 0.00003352 | 0.769654842 |
| 103 | rs34415923 | 18:1498479 | T | G | 0.193 | 2.875 | 0.00003382 | 0.689448441 |
| 104 | rs7245110 | 18:1500406 | G | A | 0.193 | 2.875 | 0.00003382 | 0.689448441 |
| 105 | rs72629390 | 14:90078701 | C | T | 0.2384 | -2.624 | 0.00003384 | 0.629256595 |
| 106 | rs10811376 | 9:20617115 | G | A | 0.3296 | 2.359 | 0.00003397 | 0.565843128 |
| 107 | rs10811364 | 9:20548755 | C | T | 0.4417 | -2.265 | 0.00003404 | 0.543295754 |
| 108 | rs12499597 | 4:181157857 | C | T | 0.4655 | 2.267 | 0.00003437 | 0.544036477 |
| 109 | rs11689432 | 2:238232864 | G | A | 0.3515 | 2.399 | 0.00003442 | 0.575852136 |
| 110 | rs746956 | 5:173707325 | A | G | 0.1692 | -2.937 | 0.00003501 | 0.705670351 |
| 111 | rs10072660 | 5:173707892 | T | C | 0.1692 | -2.937 | 0.00003501 | 0.705670351 |
| 112 | rs117201820 | 1:75486685 | A | G | 0.06524 | -4.556 | 0.00003555 | 1.095455638 |
| 113 | rs10964600 | 9:20549745 | C | A | 0.441 | -2.271 | 0.00003622 | 0.546701974 |

| | | | | | | | | |
|---|---|---|---|---|---|---|---|---|
| 114 | rs9315086 | 13:30864668 | C | T | 0.4956 | 2.287 | 0.00003671 | 0.550951578 |
| 115 | rs6659854 | 1:231603245 | A | C | 0.1903 | -2.848 | 0.00003691 | 0.68626506 |
| 116 | rs112946632 | 2:238240172 | C | A | 0.1168 | 3.54 | 0.00003745 | 0.853629129 |
| 117 | rs6744132 | 2:238235901 | G | A | 0.3521 | 2.395 | 0.00003772 | 0.577804584 |
| 118 | rs11811226 | 1:217846741 | T | A | 0.1259 | 3.286 | 0.00003773 | 0.792762364 |
| 119 | rs10915853 | 1:225654688 | T | C | 0.1062 | -3.594 | 0.00003883 | 0.868535524 |
| 120 | rs144312974 | 2:141898962 | T | C | 0.05827 | 4.723 | 0.00003904 | 1.141648538 |
| 121 | rs7240 | 2:238238488 | C | T | 0.3507 | 2.386 | 0.00003921 | 0.57688588 |
| 122 | rs7972619 | 12:94490207 | G | C | 0.468 | -2.232 | 0.00003938 | 0.539782346 |
| 123 | rs11646434 | 16:54715721 | G | A | 0.309 | 2.387 | 0.00003964 | 0.577546576 |
| **124** | **rs6887442** | **5:148112163** | **C** | **T** | **0.2994** | **-2.392** | **0.00004084** | **0.579738245** |
| 125 | rs12507155 | 4:181157743 | A | C | 0.4661 | 2.254 | 0.00004087 | 0.546291808 |
| 126 | rs10775550 | 19:29429295 | G | A | 0.2015 | 2.732 | 0.0000417 | 0.662945887 |
| 127 | rs57923757 | 16:54719206 | T | C | 0.4768 | 2.222 | 0.00004237 | 0.539582322 |
| 128 | rs9523738 | 13:92637538 | T | C | 0.3904 | -2.256 | 0.00004256 | 0.548104956 |
| 129 | rs10046877 | 9:20517596 | T | G | 0.4905 | 2.195 | 0.00004297 | 0.533543996 |
| 130 | rs80102896 | 5:115303884 | C | A | 0.08709 | 3.841 | 0.0000431 | 0.933868223 |
| 131 | rs9314349 | 8:27616685 | G | A | 0.1341 | 3.159 | 0.00004401 | 0.768800195 |
| 132 | rs2455886 | 3:105930971 | G | C | 0.4304 | 2.243 | 0.00004432 | 0.546140735 |
| 133 | rs12675095 | 8:5302690 | A | G | 0.1198 | -3.984 | 0.00004437 | 0.969107273 |
| 134 | rs68189740 | 1:217752425 | A | G | 0.09524 | 3.668 | 0.00004457 | 0.893326839 |
| 135 | rs10495067 | 1:217665359 | C | A | 0.09348 | 3.736 | 0.00004471 | 0.910109622 |
| 136 | rs200991568 | 6:168809277 | T | A | 0.05201 | -4.975 | 0.00004489 | 1.212231969 |

| | | | | | | | | |
|---|---|---|---|---|---|---|---|---|
| 137 | rs67428610 | 1:217761389 | G | A | 0.09461 | 3.672 | 0.00004513 | 0.894954911 |
| 138 | rs9661331 | 1:231616560 | T | C | 0.1625 | -3.007 | 0.000046 | 0.733772572 |
| 139 | rs9516104 | 13:92636681 | C | T | 0.3891 | -2.247 | 0.00004624 | 0.548450085 |
| 140 | rs9577115 | 13:40702417 | T | C | 0.2449 | -2.623 | 0.0000467 | 0.640537241 |
| **141** | **rs17107747** | **5:148109281** | **G** | **T** | **0.299** | **-2.376** | **0.00004726** | **0.580645161** |
| 142 | rs61688043 | 16:54719290 | A | C | 0.4793 | 2.207 | 0.00004773 | 0.539740768 |
| 143 | rs3823774 | 7:124233465 | G | A | 0.293 | 2.371 | 0.00004773 | 0.579848374 |
| 144 | rs57312975 | 11:111330650 | A | G | 0.05263 | 4.806 | 0.00004779 | 1.175348496 |
| 145 | rs11108046 | 12:95447891 | C | G | 0.07832 | -4.102 | 0.00004821 | 1.003670174 |
| 146 | rs9568231 | 13:49509462 | T | C | 0.1328 | 3.223 | 0.00004846 | 0.788790994 |
| **147** | **rs111890148** | **5:148112503** | **T** | **C** | **0.3001** | **-2.361** | **0.0000496** | **0.578676471** |
| 148 | rs12725155 | 1:217765888 | G | A | 0.09473 | 3.654 | 0.00004979 | 0.895807796 |
| **149** | **rs72660259** | **5:148112821** | **G** | **T** | **0.3005** | **-2.36** | **0.00004983** | **0.57857318** |

**S7 Table S3:** Candidate Genetic Loci for Non-planning Impulsivity Identified at $p < 5 \times 10^{-5}$.

| | SNP | Chromosome: start | Effect Allele | Other Allele | MAF | beta | pval | se |
|---|---|---|---|---|---|---|---|---|
| 1 | rs681657 | 13:42088464 | G | A | 0.4634 | 0.2397 | 0.000001107 | 0.048828682 |
| 2 | rs669642 | 13:42086709 | G | C | 0.464 | 0.2405 | 0.000001125 | 0.0490316 |
| 3 | rs2253650 | 13:42082705 | C | T | 0.4646 | 0.2392 | 0.00000134 | 0.049117043 |
| 4 | rs35208748 | 13:42084615 | A | T | 0.4635 | 0.2385 | 0.000001385 | 0.0490438 |
| 5 | rs1170203 | 13:42087243 | G | C | 0.4634 | 0.2381 | 0.000001436 | 0.049032125 |
| 6 | rs740854 | 17:12545875 | G | A | 0.3084 | -0.248 | 0.000001438 | 0.051081359 |
| 7 | rs9902200 | 17:12547675 | C | G | 0.307 | -0.2481 | 0.000001549 | 0.051260331 |

| | | | | | | | | |
|---|---|---|---|---|---|---|---|---|
| 8 | rs2013875 | 17:12545631 | T | A | 0.3082 | -0.247 | 0.00000159 | 0.051085832 |
| 9 | rs10491189 | 17:12548037 | G | A | 0.3079 | -0.2469 | 0.000001676 | 0.051181592 |
| 10 | rs9525568 | 13:42089115 | G | A | 0.3721 | -0.246 | 0.000001708 | 0.051037344 |
| 11 | rs73187215 | 13:42072633 | G | A | 0.4417 | 0.2382 | 0.000002062 | 0.049811794 |
| 12 | rs9562372 | 13:42075110 | C | T | 0.4107 | -0.2395 | 0.000002631 | 0.050623547 |
| 13 | rs12051552 | 17:12544218 | G | A | 0.3096 | -0.2418 | 0.000002732 | 0.051196274 |
| 14 | rs9532991 | 13:42080053 | A | G | 0.3741 | -0.2407 | 0.000002743 | 0.050974163 |
| 15 | rs9566906 | 13:42068410 | A | G | 0.4114 | -0.238 | 0.000002914 | 0.050530786 |
| 16 | rs1363724 | 5:148086066 | T | C | 0.4784 | 0.4979 | 0.000002963 | 0.102978283 |
| **17** | **rs9325070** | **5:148097209** | **G** | **A** | **0.3012** | **-0.244** | **0.000003054** | **0.051914894** |
| 18 | rs7993986 | 13:42087606 | A | G | 0.3754 | -0.2383 | 0.000003311 | 0.050886184 |
| 19 | rs9532990 | 13:42079762 | T | C | 0.3749 | -0.2378 | 0.000003448 | 0.050877193 |
| 20 | rs9566910 | 13:42076866 | A | G | 0.4103 | -0.2359 | 0.000003601 | 0.05056806 |
| 21 | rs73128139 | 2:235093183 | T | C | 0.06886 | -0.4476 | 0.000005192 | 0.097580118 |
| 22 | rs9913289 | 17:12558720 | C | A | 0.2812 | -0.2378 | 0.000005358 | 0.051910063 |
| 23 | rs757277 | 16:2951034 | A | G | 0.1175 | 0.3423 | 0.000006136 | 0.075214239 |
| 24 | rs57768809 | 16:2953946 | A | G | 0.1383 | 0.3195 | 0.00000731 | 0.07079548 |
| 25 | rs148127914 | 22:27742399 | T | C | 0.06707 | -0.4234 | 0.000007431 | 0.093880266 |
| 26 | rs141462441 | 22:27736681 | G | C | 0.06378 | -0.4327 | 0.000007551 | 0.096006213 |
| **27** | **rs9791022** | **5:148097621** | **G** | **A** | **0.3036** | **-0.235** | **0.000008135** | **0.05233853** |
| 28 | rs4467588 | 4:146996669 | A | G | 0.2764 | 0.2452 | 0.000008248 | 0.054646757 |
| 29 | rs7999315 | 13:42072457 | T | G | 0.4706 | 0.2221 | 0.000008288 | 0.049509585 |
| 30 | rs1170183 | 13:42076597 | C | A | 0.4706 | 0.222 | 0.000008496 | 0.049542513 |

| | | | | | | | | |
|---|---|---|---|---|---|---|---|---|
| 31 | rs1443607 | 18:7856858 | T | G | 0.05288 | -0.4796 | 0.000008672 | 0.10714924 |
| 32 | rs9303057 | 17:12542900 | G | A | 0.3141 | -0.2276 | 0.000009044 | 0.050951422 |
| 33 | rs17022238 | 4:146994415 | C | T | 0.2766 | 0.2436 | 0.000009274 | 0.054594352 |
| 34 | rs2118258 | 4:146996478 | A | G | 0.2766 | 0.2436 | 0.000009274 | 0.054594352 |
| 35 | rs28628802 | 4:146994293 | T | G | 0.2758 | 0.243 | 0.000009557 | 0.054545455 |
| 36 | rs79944343 | 19:47975257 | T | C | 0.2116 | 0.2674 | 0.000009797 | 0.060103394 |
| 37 | rs17022216 | 4:146984223 | A | G | 0.2754 | 0.2428 | 0.000009838 | 0.054574062 |
| 38 | rs141939843 | 22:27742580 | G | A | 0.06766 | -0.4159 | 0.0000101 | 0.093607923 |
| 39 | rs12918697 | 16:2952850 | C | G | 0.1325 | 0.3188 | 0.00001016 | 0.071769473 |
| 40 | rs13150180 | 4:146997133 | T | C | 0.2761 | 0.2425 | 0.00001037 | 0.054654046 |
| 41 | rs9525566 | 13:42069058 | A | G | 0.4701 | 0.2193 | 0.00001058 | 0.049469885 |
| 42 | rs117104172 | 22:27737576 | A | G | 0.06587 | -0.419 | 0.0000106 | 0.094539711 |
| 43 | rs117514820 | 22:27739715 | T | C | 0.06587 | -0.419 | 0.0000106 | 0.094539711 |
| 44 | rs149581578 | 22:27739806 | T | C | 0.06587 | -0.419 | 0.0000106 | 0.094539711 |
| 45 | rs9532985 | 13:42070241 | A | C | 0.4706 | 0.2195 | 0.00001063 | 0.049526173 |
| 46 | rs35488106 | 4:147003215 | T | G | 0.276 | 0.2418 | 0.00001069 | 0.054582393 |
| 47 | rs721038 | 12:117419512 | G | A | 0.05516 | -0.4763 | 0.00001071 | 0.10751693 |
| 48 | rs28735087 | 4:146994311 | T | A | 0.2764 | 0.2415 | 0.00001097 | 0.054576271 |
| 49 | rs71616517 | 4:147000737 | A | G | 0.2764 | 0.2418 | 0.00001097 | 0.05465642 |
| 50 | rs1443611 | 18:7849916 | C | A | 0.05216 | -0.481 | 0.00001113 | 0.108798914 |
| 51 | rs140753781 | 22:27740183 | T | C | 0.06483 | -0.4209 | 0.00001116 | 0.095204705 |
| 52 | rs76613669 | 4:146997843 | T | C | 0.2766 | 0.241 | 0.00001155 | 0.054611375 |
| 53 | rs1443610 | 18:7856337 | G | A | 0.0521 | -0.4801 | 0.00001159 | 0.108816863 |

| | | | | | | | | |
|---|---|---|---|---|---|---|---|---|
| 54 | rs1443609 | 18:7856405 | C | T | 0.0521 | -0.4801 | 0.00001159 | 0.108816863 |
| 55 | rs7223735 | 17:12559996 | G | T | 0.283 | -0.2278 | 0.00001169 | 0.051655329 |
| 56 | rs1443612 | 18:7849575 | A | G | 0.05241 | -0.4812 | 0.0000117 | 0.109115646 |
| 57 | rs1412934 | 13:94241452 | T | C | 0.1945 | 0.2691 | 0.00001171 | 0.061020408 |
| 58 | rs56914158 | 15:31949047 | T | A | 0.2521 | -0.243 | 0.00001195 | 0.055152065 |
| 59 | rs1530389 | 18:7850719 | C | T | 0.05222 | -0.4792 | 0.00001219 | 0.108884344 |
| 60 | rs386283 | 13:42368127 | A | G | 0.1954 | -0.272 | 0.00001225 | 0.061818182 |
| 61 | rs28430726 | 4:146994333 | G | C | 0.2758 | 0.24 | 0.00001236 | 0.054570259 |
| 62 | rs1443613 | 18:7847574 | T | C | 0.05222 | -0.4787 | 0.00001245 | 0.108894449 |
| 63 | rs61040170 | 17:12549009 | G | A | 0.291 | -0.2305 | 0.0000125 | 0.052434031 |
| 64 | rs147648611 | 22:27740381 | T | C | 0.06535 | -0.4171 | 0.0000125 | 0.094881711 |
| 65 | rs9917863 | 4:146986593 | G | C | 0.2755 | 0.24 | 0.00001269 | 0.054644809 |
| 66 | rs149285517 | 4:146993355 | T | C | 0.1709 | 0.2918 | 0.00001298 | 0.066514703 |
| 67 | rs7240568 | 18:7847293 | C | T | 0.05241 | -0.4783 | 0.0000132 | 0.109101277 |
| 68 | rs9918010 | 4:146986972 | A | G | 0.2755 | 0.2385 | 0.00001371 | 0.054514286 |
| 69 | rs62184861 | 2:203811709 | G | A | 0.08443 | 0.3846 | 0.00001377 | 0.087928669 |
| 70 | rs80108322 | 10:18737742 | C | T | 0.2368 | -0.2573 | 0.00001384 | 0.058838326 |
| 71 | rs178289 | 22:20987880 | G | C | 0.3291 | 0.2267 | 0.00001433 | 0.051935853 |
| 72 | rs882064 | 17:10251446 | A | G | 0.2132 | 0.2577 | 0.00001449 | 0.059064864 |
| 73 | rs76642275 | 10:18768074 | G | T | 0.2303 | -0.2593 | 0.00001482 | 0.059499771 |
| 74 | rs72809149 | 17:12540561 | C | T | 0.3162 | -0.2205 | 0.00001515 | 0.050654721 |
| 75 | rs28652568 | 1:170942424 | T | C | 0.4213 | 0.2467 | 0.00001528 | 0.056647532 |
| 76 | rs11927760 | 3:85026941 | A | G | 0.3838 | -0.217 | 0.00001566 | 0.049942463 |

| | | | | | | | | |
|---|---|---|---|---|---|---|---|---|
| 77 | rs2267115 | 22:27759464 | A | G | 0.3635 | 0.2229 | 0.00001569 | 0.051276743 |
| 78 | rs2439652 | 5:3204807 | C | T | 0.09484 | -0.3552 | 0.00001651 | 0.081975537 |
| 79 | rs17056526 | 5:159074842 | T | C | 0.315 | 0.227 | 0.00001797 | 0.052619379 |
| 80 | rs55937021 | 17:12537772 | A | G | 0.3157 | -0.2188 | 0.00001804 | 0.05073035 |
| 81 | rs7998896 | 13:111897554 | G | A | 0.2964 | 0.2257 | 0.00001821 | 0.052354442 |
| 82 | rs1348407 | 18:7849171 | C | T | 0.05168 | -0.4716 | 0.00001826 | 0.109394572 |
| 83 | rs9561517 | 13:94244440 | T | G | 0.1952 | 0.261 | 0.00001913 | 0.060697674 |
| 84 | rs56262049 | 13:42070569 | C | T | 0.2964 | -0.2269 | 0.00001993 | 0.052878117 |
| 85 | rs12584909 | 13:42076496 | A | G | 0.2964 | -0.2269 | 0.00001993 | 0.052878117 |
| 86 | rs7970835 | 12:95556850 | T | C | 0.06498 | -0.4331 | 0.00002026 | 0.101002799 |
| 87 | rs11732596 | 4:9567737 | T | C | 0.3845 | -0.2247 | 0.00002119 | 0.052524544 |
| 88 | rs12413388 | 10:18775012 | A | G | 0.2362 | -0.2506 | 0.00002162 | 0.058661049 |
| 89 | rs2973823 | 5:97710594 | G | T | 0.1918 | -0.2626 | 0.00002203 | 0.061527648 |
| 90 | rs11507721 | 9:88674429 | T | C | 0.213 | 0.6997 | 0.00002221 | 0.159931429 |
| 91 | rs16917797 | 10:18772960 | T | C | 0.2371 | -0.2488 | 0.00002273 | 0.058390049 |
| 92 | rs7732695 | 5:113159724 | C | T | 0.1847 | -0.2699 | 0.00002294 | 0.063371683 |
| 93 | rs78395975 | 20:54827004 | C | T | 0.09461 | 0.3436 | 0.00002355 | 0.080790031 |
| 94 | rs6098277 | 20:54828412 | C | T | 0.09461 | 0.3436 | 0.00002355 | 0.080790031 |
| 95 | rs6098278 | 20:54828732 | G | C | 0.09461 | 0.3436 | 0.00002355 | 0.080790031 |
| 96 | rs6098279 | 20:54828759 | C | T | 0.09461 | 0.3436 | 0.00002355 | 0.080790031 |
| 97 | rs6098281 | 20:54828842 | C | A | 0.09461 | 0.3436 | 0.00002355 | 0.080790031 |
| 98 | rs7560067 | 2:235089214 | C | T | 0.06527 | -0.425 | 0.00002371 | 0.099976476 |
| 99 | rs10169551 | 2:235090233 | G | C | 0.06527 | -0.425 | 0.00002371 | 0.099976476 |

| | | | | | | | | |
|---|---|---|---|---|---|---|---|---|
| 100 | rs75960612 | 17:51463294 | T | C | 0.0503 | -0.4696 | 0.00002471 | 0.110702499 |
| 101 | rs6091985 | 20:54829400 | C | G | 0.09472 | 0.3418 | 0.00002633 | 0.080861131 |
| 102 | rs6506536 | 18:7845329 | T | G | 0.05282 | -0.4587 | 0.00002659 | 0.108568047 |
| 103 | rs2287769 | 5:148111736 | T | C | 0.3012 | -0.2215 | 0.00002699 | 0.052475717 |
| 104 | rs9268784 | 6:32453015 | G | A | 0.4534 | 0.215 | 0.00002705 | 0.050899621 |
| 105 | rs16917798 | 10:18773880 | G | C | 0.2371 | -0.2484 | 0.00002708 | 0.058848614 |
| 106 | rs16917736 | 10:18737378 | A | G | 0.2356 | -0.2476 | 0.00002712 | 0.058672986 |
| 107 | rs6506538 | 18:7855870 | G | A | 0.05102 | -0.4644 | 0.00002727 | 0.110073477 |
| **108** | **rs111890148** | **5:148112503** | **T** | **C** | **0.3012** | **-0.2214** | **0.00002752** | **0.052501779** |
| 109 | rs9272367 | 6:32636848 | T | C | 0.07296 | 0.5037 | 0.00002773 | 0.11904987 |
| 110 | rs9663531 | 10:18786943 | G | A | 0.2383 | -0.2446 | 0.00002793 | 0.058044613 |
| 111 | rs1029372 | 9:133613320 | G | A | 0.4994 | 0.2064 | 0.00002833 | 0.049026128 |
| 112 | rs4863426 | 4:188965408 | G | A | 0.1151 | -0.3284 | 0.00002896 | 0.078097503 |
| 113 | rs7569005 | 2:235088912 | A | G | 0.06475 | -0.4218 | 0.00002928 | 0.100356888 |
| 114 | rs45552531 | 22:27746936 | T | G | 0.06707 | -0.395 | 0.0000295 | 0.094025232 |
| 115 | rs3826742 | 19:7848866 | C | T | 0.4339 | -0.2025 | 0.00002967 | 0.048214286 |
| 116 | rs2033605 | 7:12700591 | C | T | 0.2222 | 0.2421 | 0.00003015 | 0.057697807 |
| 117 | rs8089088 | 18:7836178 | T | C | 0.05396 | -0.4463 | 0.00003019 | 0.106363203 |
| 118 | rs146674603 | 4:147997344 | G | A | 0.07006 | 0.39 | 0.00003067 | 0.093034351 |
| 119 | rs1862443 | 5:148109518 | C | T | 0.3007 | -0.2209 | 0.00003089 | 0.052720764 |
| 120 | rs10898007 | 11:82617297 | T | C | 0.2988 | 0.2224 | 0.00003141 | 0.053129479 |
| 121 | rs16917808 | 10:18781406 | C | T | 0.2395 | -0.2427 | 0.00003161 | 0.057992832 |
| 122 | rs16917807 | 10:18781335 | C | T | 0.2383 | -0.2427 | 0.00003184 | 0.058020559 |

| | | | | | | | | |
|---|---|---|---|---|---|---|---|---|
| 123 | rs74418544 | 10:18731916 | G | A | 0.2365 | -0.2452 | 0.00003194 | 0.058618217 |
| 124 | rs13246307 | 7:83317060 | G | T | 0.06358 | -0.9036 | 0.00003197 | 0.211022887 |
| 125 | rs13112205 | 4:154345207 | T | C | 0.2515 | 0.2363 | 0.00003219 | 0.05651758 |
| 126 | rs13112673 | 4:154345287 | A | G | 0.2515 | 0.2363 | 0.00003219 | 0.05651758 |
| 127 | rs6098271 | 20:54827341 | A | T | 0.09401 | 0.3386 | 0.00003237 | 0.081024168 |
| 128 | rs6098276 | 20:54828254 | A | G | 0.09401 | 0.3386 | 0.00003237 | 0.081024168 |
| 129 | rs6098280 | 20:54828801 | T | A | 0.09401 | 0.3386 | 0.00003237 | 0.081024168 |
| **130** | **rs72660259** | **5:148112821** | **G** | **T** | **0.3016** | **-0.2194** | **0.00003262** | **0.052513164** |
| 131 | rs3815740 | 5:148111630 | G | A | 0.3016 | -0.2193 | 0.00003306 | 0.05253953 |
| 132 | rs7771461 | 6:124047665 | A | G | 0.4587 | 0.2075 | 0.00003345 | 0.049736337 |
| 133 | rs16917823 | 10:18786029 | T | C | 0.2389 | -0.2421 | 0.00003346 | 0.058029722 |
| 134 | rs12657808 | 5:148103561 | G | C | 0.3008 | -0.2186 | 0.00003396 | 0.052447217 |
| **135** | **rs17107747** | **5:148109281** | **G** | **T** | **0.2995** | **-0.2197** | **0.00003402** | **0.052711132** |
| 136 | rs828407 | 5:3199691 | A | G | 0.1012 | -0.3335 | 0.00003444 | 0.080072029 |
| 137 | rs6098275 | 20:54828175 | A | G | 0.09581 | 0.3353 | 0.00003468 | 0.080542878 |
| 138 | rs146365946 | 20:54855626 | C | T | 0.09353 | 0.3388 | 0.00003473 | 0.081383618 |
| 139 | rs3777142 | 5:148107956 | A | T | 0.3018 | -0.2184 | 0.00003538 | 0.052512623 |
| 140 | rs3815741 | 5:148109012 | C | T | 0.3018 | -0.2184 | 0.00003538 | 0.052512623 |
| 141 | rs1862442 | 5:148109707 | T | C | 0.3018 | -0.2184 | 0.00003538 | 0.052512623 |
| 142 | rs1862440 | 5:148109841 | G | A | 0.3018 | -0.2184 | 0.00003538 | 0.052512623 |
| 143 | rs58154955 | 5:148110246 | G | T | 0.3018 | -0.2184 | 0.00003538 | 0.052512623 |
| 144 | rs17107754 | 5:148110313 | C | T | 0.3018 | -0.2184 | 0.00003538 | 0.052512623 |
| 145 | rs72660256 | 5:148110454 | G | T | 0.3018 | -0.2184 | 0.00003538 | 0.052512623 |

| | | | | | | | | |
|---|---|---|---|---|---|---|---|---|
| 146 | rs58670387 | 5:148110597 | A | G | 0.3018 | -0.2184 | 0.00003538 | 0.052512623 |
| 147 | rs2287767 | 5:148111989 | G | A | 0.3018 | -0.2184 | 0.00003538 | 0.052512623 |
| 148 | rs2287766 | 5:148112014 | T | C | 0.3018 | -0.2184 | 0.00003538 | 0.052512623 |
| 149 | rs34975049 | 5:148112318 | G | T | 0.3018 | -0.2184 | 0.00003538 | 0.052512623 |
| 150 | rs55816145 | 5:148113099 | A | C | 0.3018 | -0.2184 | 0.00003538 | 0.052512623 |
| 151 | rs41291433 | 5:148113116 | T | G | 0.3018 | -0.2184 | 0.00003538 | 0.052512623 |
| 152 | rs17107762 | 5:148113347 | G | A | 0.3018 | -0.2184 | 0.00003538 | 0.052512623 |
| 153 | rs72660260 | 5:148113431 | A | C | 0.3018 | -0.2184 | 0.00003538 | 0.052512623 |
| 154 | rs17107763 | 5:148113690 | T | C | 0.3018 | -0.2184 | 0.00003538 | 0.052512623 |
| 155 | rs3815739 | 5:148113821 | A | T | 0.3018 | -0.2184 | 0.00003538 | 0.052512623 |
| 156 | rs3815738 | 5:148114037 | T | A | 0.3018 | -0.2184 | 0.00003538 | 0.052512623 |
| 157 | rs1443608 | 18:7856564 | C | A | 0.05384 | -0.4704 | 0.00003604 | 0.113158528 |
| 158 | rs13113312 | 4:154345579 | T | G | 0.2515 | 0.235 | 0.00003637 | 0.056599229 |
| 159 | rs9566908 | 13:42071845 | T | C | 0.3663 | -0.2131 | 0.00003718 | 0.051386544 |
| 160 | rs55862047 | 5:148109967 | G | T | 0.3013 | -0.2177 | 0.00003849 | 0.052597246 |
| **161** | **rs6887442** | **5:148112163** | **C** | **T** | **0.3005** | **-0.2175** | **0.00003854** | **0.052548925** |
| 162 | rs2021802 | 13:42078029 | T | C | 0.3665 | -0.2127 | 0.00003855 | 0.051389224 |
| 163 | rs12651103 | 4:154349807 | T | C | 0.2503 | 0.2338 | 0.00003881 | 0.056514382 |
| 164 | rs2798713 | 9:15397246 | G | A | 0.1029 | -1.314 | 0.00003907 | 0.250428816 |
| 165 | rs4773782 | 13:94231472 | G | A | 0.203 | 0.2447 | 0.00003928 | 0.059192066 |
| 166 | rs145307807 | 22:27741212 | T | C | 0.06775 | -0.3829 | 0.0000396 | 0.092666989 |
| 167 | rs1844548 | 10:18764967 | T | G | 0.2371 | -0.2422 | 0.00003974 | 0.058615682 |
| 168 | rs13101995 | 4:63617607 | C | G | 0.1401 | 0.2984 | 0.00003998 | 0.072251816 |

| | | | | | | | | |
|---|---|---|---|---|---|---|---|---|
| 169 | rs56814474 | 5:113161220 | G | A | 0.185 | -0.2616 | 0.0000405 | 0.063387449 |
| 170 | rs10199494 | 2:19116578 | A | T | 0.1277 | -0.2964 | 0.00004054 | 0.071819724 |
| 171 | rs76568969 | 14:49913745 | C | T | 0.08898 | 0.3569 | 0.00004061 | 0.086479283 |
| 172 | rs17311657 | 4:13073519 | C | T | 0.1545 | -0.279 | 0.00004086 | 0.067636364 |
| 173 | rs61600033 | 5:148110878 | A | G | 0.3018 | -0.216 | 0.00004091 | 0.052363636 |
| 174 | rs3100870 | 14:49913221 | C | T | 0.08802 | 0.3568 | 0.00004111 | 0.086517944 |
| 175 | rs8012916 | 14:49913632 | C | T | 0.06944 | 0.4034 | 0.00004132 | 0.097817653 |
| 176 | rs78486695 | 5:113163223 | A | G | 0.1832 | -0.2628 | 0.00004132 | 0.063755459 |
| 177 | rs72660258 | 5:148111298 | G | C | 0.3011 | -0.2166 | 0.00004185 | 0.052585579 |
| 178 | rs79825075 | 1:43271271 | T | C | 0.2114 | -0.2426 | 0.00004192 | 0.058897791 |
| 179 | rs7681342 | 4:154349308 | T | G | 0.2491 | 0.2336 | 0.00004252 | 0.05675413 |
| 180 | rs111765057 | 10:18789516 | A | G | 0.2386 | -0.2385 | 0.00004313 | 0.058000973 |
| 181 | rs79494654 | 20:50290285 | A | G | 0.1479 | -0.2813 | 0.00004319 | 0.068409533 |
| 182 | rs72936545 | 2:203813532 | C | A | 0.08683 | 0.3579 | 0.00004349 | 0.087080292 |
| 183 | rs12589830 | 14:49921185 | A | G | 0.06527 | 0.4029 | 0.00004373 | 0.098053054 |
| 184 | rs3784599 | 15:31067254 | T | G | 0.1447 | -0.286 | 0.00004381 | 0.06960331 |
| 185 | rs9524382 | 13:94239256 | A | G | 0.1996 | 0.2464 | 0.00004471 | 0.060038986 |
| 186 | rs36188935 | 18:7834309 | A | G | 0.05241 | -0.4485 | 0.00004516 | 0.109336909 |
| 187 | rs722885 | 12:28081090 | G | A | 0.4137 | 0.2031 | 0.00004536 | 0.049536585 |
| 188 | rs1994607 | 10:18756660 | G | C | 0.2353 | -0.2412 | 0.0000454 | 0.058829268 |
| 189 | rs11774735 | 8:4922059 | C | A | 0.3453 | 0.2076 | 0.00004554 | 0.050646499 |
| 190 | rs16917752 | 10:18755788 | G | A | 0.2365 | -0.2404 | 0.00004558 | 0.058648451 |
| 191 | rs16917771 | 10:18761005 | A | G | 0.2365 | -0.2404 | 0.00004558 | 0.058648451 |

| | | | | | | | | |
|---|---|---|---|---|---|---|---|---|
| 192 | rs7689027 | 4:154349156 | A | T | 0.2497 | 0.2315 | 0.00004589 | 0.056490971 |
| 193 | rs78936021 | 5:113162988 | C | G | 0.1814 | -0.2603 | 0.00004593 | 0.063534293 |
| 194 | rs2039130 | 10:18759818 | G | C | 0.2359 | -0.2403 | 0.00004603 | 0.058652673 |
| 195 | rs2287770 | 5:148111655 | G | A | 0.301 | -0.2156 | 0.00004684 | 0.052675299 |
| 196 | rs1348104 | 18:7842734 | C | A | 0.052 | -0.4491 | 0.00004706 | 0.109750733 |
| 197 | rs522115 | 6:83702974 | C | T | 0.1287 | -0.3025 | 0.00004784 | 0.073997065 |
| 198 | rs2377577 | 18:7836714 | T | C | 0.05288 | -0.444 | 0.00004819 | 0.10866373 |
| 199 | rs12587527 | 14:49921196 | C | T | 0.08683 | 0.3566 | 0.00004866 | 0.087316357 |
| 200 | rs2165932 | 5:113163222 | A | G | 0.1838 | -0.2589 | 0.00004928 | 0.063440333 |
| 201 | rs118148284 | 22:27734298 | A | G | 0.06287 | -0.3934 | 0.00004959 | 0.096445207 |
| 202 | rs7166431 | 15:31063601 | T | C | 0.142 | -0.2855 | 0.0000499 | 0.070009809 |

**S7 Table S4:** Gene Ontology (GO) Annotations for SPINK5.

| Aspect | Term |
|---|---|
| Cellular Component | cell cortex |
| | cytoplasm |
| | cytosol |
| | endoplasmic reticulum |
| | endoplasmic reticulum membrane |
| | epidermal lamellar body |
| | extracellular region |
| | intracellular membrane-bounded organelle |
| | perinuclear region of cytoplasm |
| Molecular Function | serine-type endopeptidase inhibitor activity |
| Biological Process | cell differentiation |
| | epidermal cell differentiation |
| | epithelial cell differentiation |
| | extracellular matrix organization |
| | hair cell differentiation |
| | negative regulation of angiogenesis |
| | negative regulation of antibacterial peptide production |
| | negative regulation of immune response |
| | negative regulation of proteolysis |
| | regulation of cell adhesion |
| | regulation of T cell differentiation |
| | regulation of timing of anagen |

**S7 Table S5:** Gene Ontology (GO) Annotations for FBXO38-DT.

| Aspect | Term |
|---|---|
| Cellular Component | cytoplasm |
| | cytosol |
| | nucleus |
| | SCF ubiquitin ligase complex |
| Molecular Function | ubiquitin-like ligase-substrate adaptor activity |
| Biological Process | adaptive immune response |
| | positive regulation of neuron projection development |
| | positive regulation of T cell activation |
| | positive regulation of T cell mediated immune response to tumor cell |
| | protein K48-linked ubiquitination |
| | SCF-dependent proteasomal ubiquitin-dependent protein catabolic process |

**SI References**

1. Z. Chen, P. Liu, C. Zhang, T. Feng, Brain morphological dynamics of procrastination: The crucial role of the self-control, emotional, and episodic prospection network. *Cerebral Cortex* **30**, 2834–2853 (2020).
2. R. Zhang, Z. Chen, T. Xu, L. Zhang, T. Feng, The overlapping region in right hippocampus accounting for the link between trait anxiety and procrastination. *Neuropsychologia* **146**, 107571 (2020).
3. P. Liu, T. Feng, The overlapping brain region accounting for the relationship between procrastination and impulsivity: A voxel-based morphometry study. *Neuroscience* **360**, 9–17 (2017).
4. Y. Hu, P. Liu, Y. Guo, T. Feng, The neural substrates of procrastination: A voxel-based morphometry study. *Brain and cognition* **121**, 11–16 (2018).
5. Z. Chen, *et al.*, Hybrid brain model accurately predict human procrastination behavior. *Cogn Neurodyn* **16**, 1107–1121 (2022).
6. Y. Hu, *et al.*, Mapping neurogenetic characteristics of psychopathological procrastination using normative modeling in a prospective twin cohort. *bioRxiv* 2025–06 (2025).
7. Y. Chen, *et al.*, SOAPnuke: a MapReduce acceleration-supported software for integrated quality control and preprocessing of high-throughput sequencing data. *Gigascience* **7**, gix120 (2018).
8. H. Li, R. Durbin, Fast and accurate short read alignment with Burrows–Wheeler transform. *bioinformatics* **25**, 1754–1760 (2009).
9. M. A. DePristo, *et al.*, A framework for variation discovery and genotyping using next-generation DNA sequencing data. *Nature genetics* **43**, 491–498 (2011).
10. S. Purcell, *et al.*, PLINK: a tool set for whole-genome association and population-based linkage analyses. *The American journal of human genetics* **81**, 559–575 (2007).
11. J. Fu, *et al.*, Cross-ancestry genome-wide association studies of brain imaging phenotypes. *Nature Genetics* 1–11 (2024).
